# Early evolution of BA.2.86 sheds light on the origins of highly divergent SARS-CoV-2 lineages

**DOI:** 10.1101/2024.07.18.604213

**Authors:** Marina Escalera-Zamudio, Cedric C S Tan, Lucy van Dorp, François Balloux

## Abstract

Since the emergence of SARS-CoV-2 in late 2019, highly divergent variants with novel constellations of mutations have periodically emerged to displace previously co-circulating virus lineages. The evolutionary mechanisms behind this process remain unclear, but the prevailing hypothesis supports emergence through single events followed by a rapid accumulation of mutations in individual long-term infections. In July 2023, the Omicron-descending BA.2.86 Variant of Interest (VOI) emerged to outcompete other virus lineages, with descending sublineages still dominating as of December 2024. We use BA.2.86 as a case study to test the hypothesis that highly divergent variants emerge during a single chronic infection. By applying a comprehensive suite of evolutionary analyses to independent high-resolution datasets, we identify a cluster of approximately 100 BA.2 genomes that fall directly ancestral and shorten the branch leading to BA.2.86, carrying some BA.2.86-specific lineage defining mutations. These genomes, sampled from multiple countries and at different timepoints, share a single ancestor dating back to December 2021. Additionally, we detect over 2600 earlier BA.2, BA.2.75, and XBB.1.5 genomes with multiple mutations later associated with BA.2.86. These genomes represent a standing genetic variation that likely facilitated a gradual evolution in the direct ancestry of this variant. We further uncover a complex evolutionary process involving recombination between the BA.2.75 and XBB.1.5 parental lineages. Together, our results challenge the notion of a single emergence event, and instead support a more complex scenario, with BA.2.86 likely having evolved over multiple transmission chains.

## INTRODUCTION

The evolution of SARS-CoV-2 has been characterised by the recurrent emergence of highly divergent variants that replace previously dominating virus lineages in circulation. This process has consistently led to complete turnovers in the virus population. The repeated emergence of viral lineages with enhanced epidemiological success has been associated with the acquisition of constellations of mutations conferring more transmissible and/or immune evasive phenotypes^1^. Nonetheless, the underlying evolutionary mechanisms driving this process have been rarely formally investigated. Consequently, how these highly divergent SARS-CoV-2 variants emerge remains only partially understood. Investigating the drivers behind the emergence and evolution of highly divergent SARS-CoV-2 variants is not only essential for a retrospective analysis of the COVID-19 pandemic, but can also provide insights into how virus population turnovers occur, informing future surveillance and outbreak control measures.

From a phylogenetic perspective, the emergence of highly divergent variants is distinguished by a long branch separating a newly designated clade from its direct ancestors^2,3^. These long branches reflect a rapid accumulation of mutations relative to the most closely related parental lineages, or could alternatively reflect a significant undersampling of viral diversity. In this light, several evolutionary scenarios have been proposed to explain how highly divergent SARS-CoV-2 lineages emerge (extensively reviewed by Markov et al)^4^. The two main hypotheses are: 1) single emergence events linked to individual infection cases, where multiple mutations rapidly accumulate over a short period of time (*i.e.,* accelerated evolution), and 2) a gradual accumulation of mutations over time across independent transmission chains. Hypothesis 1 is supported by the observation of multiple mutations acquired during chronic infections corresponding to immunocompromised patients^5–7^, highlighting the potential for a rapid accumulation of amino acid changes within single infections. Hypothesis 2 is supported by reports of undetected circulation (*i.e.,* cryptic circulation) of virus lineages within areas with poor genomic surveillance, which are only identified later in regions with a more robust detection capacity^8^.

The World Health Organization’s (WHO) criteria for classifying highly divergent variants involves naming emerging viral subpopulations as ‘Variants of Interest’ (VOI), ‘Variants under Monitoring’ (VUM) or ‘Variants of Concern’ (VOC), ranked according to their potential epidemiological risk^9^. The last VOC designation corresponds to the Omicron variant (labelled as lineage B.1.1.529 following the Pango nomenclature system^10^), detected in January 2022^9,11^. With over 50 ‘novel’ mutations, B.1.1.529 rapidly spread to dominate globally. Nineteen months later, the detection of BA.2.86 in July 2023 —directly descending from Omicron — marked the first instance of apparent accelerated evolution since the emergence of B.1.1.529^12^. Relative to its parental lineage (BA.2, also an Omicron descending lineage), BA.2.86 carried 45 mutations, including over 30 amino acid changes in the Spike protein alone^13^. BA.2.86 was first identified in Denmark, the UK, the USA, and Israel^12,13^ and quickly spread worldwide to be recognized by the WHO as a VOI by November 2023^14^. BA.2.86 further diversified to give rise to the JN.1* lineages, characterized by a single additional non-synonymous mutation within the Spike protein (L455S), and three synonymous mutations in other genes^15^. Further continuous evolution has been evidenced by the emergence of the so-called “FLiRT variants”^13^ (designated as KP.*), which acquired convergent mutations at positions 456, 346, and 572 of the Spike protein. With an increasing prevalence, FLiRT variants dominated across the USA in July 2024, and were (partially) responsible for the infection wave recorded during that Summer^16^. BA.2.86-descending sublineages still dominated worldwide as of December 2024^17^.

Here, we use BA.2.86 as a case study to test the hypothesis of a single emergence event for this highly divergent SARS-CoV-2 variant. We compiled four high-resolution datasets from the GISAID repository^18–21^, comprising genomes assigned to BA.2.86* and to its most closely related ancestral lineages (BA.2, BA.2.75, and XBB.1.5). These datasets capture the evolutionary history of the BA.2* Omicron-descending lineages, leading to the emergence and evolution of BA.2.8 up to May 2024. Applying a suite of evolutionary analyses to these independent datasets, we investigated the mechanisms underlying the emergence of this highly divergent variant. Our results reveal that BA.2.86 is unlikely to have emerged through a single event following accelerated evolution associated with a long-term infection case. Instead, our results are consistent with the hypothesis of a gradual accumulation of mutations across multiple transmission chains, with other processes, such as recombination, contributing to its emergence. Together, our findings provide insights into the broader mechanisms that may drive the evolution and emergence of highly divergent SARS-CoV-2 lineages.

## RESULTS

### Identification of BA.2.86-like evolutionary intermediates

To capture the evolution of lineage BA.2.86 over time, we used longitudinally retrieved publicly available genome assemblies from GISAID to reconstruct four phylogenetic trees comprising the evolution of the BA.2* Omicron-descending sublineages (*Supplementary Figure 1a-e*). Our ‘initial dataset’ (n=100. Compiled as of August 21^st^ 2023) recapitulates the early characterization of BA.2.86^12^, as shown in the Nextstrain reference phylogeny (visualised between August 2023 and May 2024)^2^, where this lineage directly descends from the BA.2 clade separated by a long branch (*Supplementary Figure 1a*). However, following a wider sampling of the expanding BA.2.86* diversity represented in our ‘full dataset’ (n=7,760. Compiled as of March 15^th^ 2024), we identified a cluster of 106 BA.2 genomes that fall directly ancestral to BA.2.86 and shorten this long branch (**Fig. 1a**), representing putative evolutionary intermediates leading to the emergence of BA.2.86 (henceforth termed ‘BA.2.86-like intermediates’) (*Supplementary Data 1–Supplementary Table 1*), (*Supplementary Data 1–Supplementary Table 1*). A consistent phylogenetic placement of this cluster and all clades comprising other BA.2* Omicron-descending sublineages was observed across all high-resolution datasets, compiled using various subsampling schemes and inclusion criteria, including a streamlined quality-controlled version of the data used for downstream analyses (*Supplementary Figure 1b-d*).

**Figure 1.**
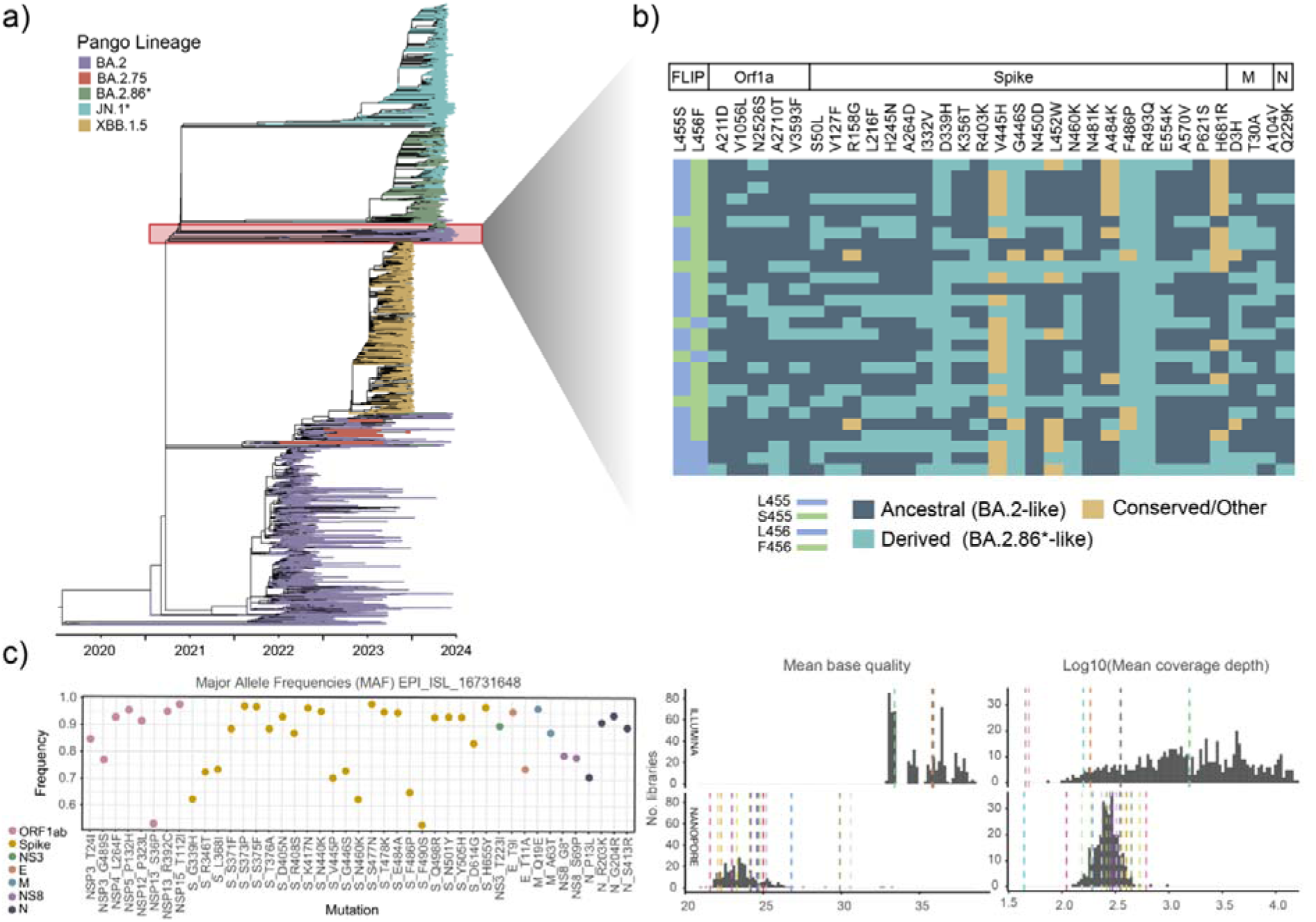
Identification of BA.2.86-like evolutionary intermediates. (a) The time-scaled ML consensus tree corresponding to our “full dataset” with branches coloured according to ‘Pango’ lineage. The phylogenetic placement of putative BA.2.86-like intermediates is represented by a cluster of around 100 BA.2-assigned genomes (highlighted with a red box) effectively shortening the branch leading to the BA.2.86* clade. (b) Heatmap showing a variable presence/absence pattern of BA.2.86-specific lineage defining mutations (LDMs) for a representative set of these genomes, rendering them genetically ‘BA.2.86-like’. LDMs are shown as ‘derived’ traits (teal), while ‘ancestral’ traits (i.e., those present in other closely related BA.2* lineages but not in BA.2.86*) are shown in dark blue. Conserved or other states are shown in ochre. (c) An example distribution of Major Allele Frequencies (MAF) for assembly EPI_ISL_16731648, with no evidence of autocorrelation. Only six alleles correspond to LDMs occurring within Spike (G339H, V445P, G446S, N460K, E484A and F486P), all corresponding to MAFs >60%. Distribution plots showing the mean base quality and mean coverage depth values for the 23 assemblies analysed, contrasted to a background distribution of 901 ‘high-quality’ SARS-CoV-2 sequencing libraries generated within the same timeframe. For all metrics, mean values generally fall within the background distributions, indicating that at least a fraction of these BA.2 genomes are unlikely to be artefactual.

To better understand the contribution of non-synonymous lineage-defining mutations specific to BA.2.86 (named here LDMs, defined as of August 2023^13^. See Methods section) to the phylogenetic placement of these BA.2.86-like intermediates, we further inspected their mutational profile. By annotating LDMs as traits at the tips of the phylogenetic tree, we observed that these genomes display a variable presence pattern for at least 32 LDMs: 5 occurring in ORF1ab (A211D, V1056L, N2526S, A2710T and V3593F), 23 in Spike (S50L, V127F, R158G, L216F, H245N, A264D, I332V, D339H, K356T, R403K, V445H, G446S, N450D, L452W, N460K, N481K, A484K, F486P, R493Q, E554K, A570V, P621S and H681R), 3 in the M gene (D3H, T30A, A104V), and 1 in the N gene (Q229K) (**Fig. 1b**). The subsequent reconstruction of LDMs as ancestral states supports a variable presence pattern at multiple nodes of this cluster, evidencing these BA.2.86-like genomes as intermediate evolutionary stages (*Supplementary Data 2*). Overall, the presence of multiple LDMs in this cluster indicates that these genomes are genetically similar to BA.2.86, with LDMs directly contributing to their phylogenetic placement.

To further assess if these BA.2.86-like intermediates could be artefactual (*i.e.,* assemblies linked to mixed infections or low-quality/contaminated libraries), we re-analysed the raw sequencing data (when available) for a subset of these to assess their quality. We successfully retrieved data for 23/106 (22%) assemblies sampled from the USA, Norway, Ireland, Spain, and New Zealand, with collection dates ranging from January 2023 to February 2024 (earliest: 2023-01-30, latest: 2024-02-13) (*Supplementary Data 1 - Supplementary Table 1*). We first examined major/minor allele frequency (MAF) distributions. While we acknowledge that distinguishing between MAF patterns in mixed infections from those representing genuine intrahost diversity in single infections remains challenging^22,23^, we found no evidence for mixed infection or contamination, as many LDMs within these assemblies represent major alleles with frequencies above 80% (MAF >80%). For example, in 17/23 of the re-analysed libraries, at least one LDM corresponds to a MAF >80%, and in one library (corresponding to assembly EPI_ISL_18746333) 20/45 LDMs correspond to MAFs >80%. These results indicate that SNP calling in consensus genomes corresponding to BA.2.86-like intermediates is not likely to be a result of mixed infections.

For these sequencing libraries, we then computed the mean base quality, mean coverage depth and mean mapping quality, along with the number of single amino acid variants (SAVs) with a frequency >10% (as a rough metric for intrahost diversity). We then compared these metrics to a background distribution of publicly available SARS-CoV-2 sequencing libraries (n=901) generated within the same timeframe of our data (*Supplementary Figure 2*). Quality scores for all assemblies analysed generally fall within these background distributions (**Fig. 1c**), indicating that most assemblies corresponding to BA.2.86-like intermediates do not derive from low-quality sequencing libraries. Thus, our results indicate that the presence of putative evolutionary intermediates leading to the emergence of BA.2.86 is unlikely to be artefactual.

### Gradual evolution preceding the emergence of BA.2.86

Based on available genome-associated metadata (*Supplementary Data 1–Supplementary Table 1*), BA.2.86-like intermediates were sampled from the USA (40%), Malaysia (15%), Italy (12%), Spain (10%), and Ireland (8%), with 14% of these displaying collection dates preceding the detection of BA.2.86 (*i.e.,* EPI_ISL_16731648; sampled from the USA on 2023-01-15) (*Supplementary Text 1*). The circulation of BA.2.86-like intermediates across multiple geographic regions over an extended time period suggests that the acquisition of LDMs potentially occurred across independent transmission chains preceding BA.2.86 (**Fig. 2a**). Nonetheless, relying solely on tip dates can lead to false age estimates, as unsampled lineages and uncertainty in sampling needs to be accounted for^24^. Thus, we employed a time-scaled analysis to infer the node ages corresponding to (i) the tMRCA (root) of the tree, (ii) the tMRCA of the cluster representing evolutionary intermediates, and (iii) the tMRCA of the BA.2.86* clade (**Fig. 2a**). However, incomplete sampling across geographic regions hinders our ability to infer the geographic origins of any virus lineage^25^.

**Figure 2.**
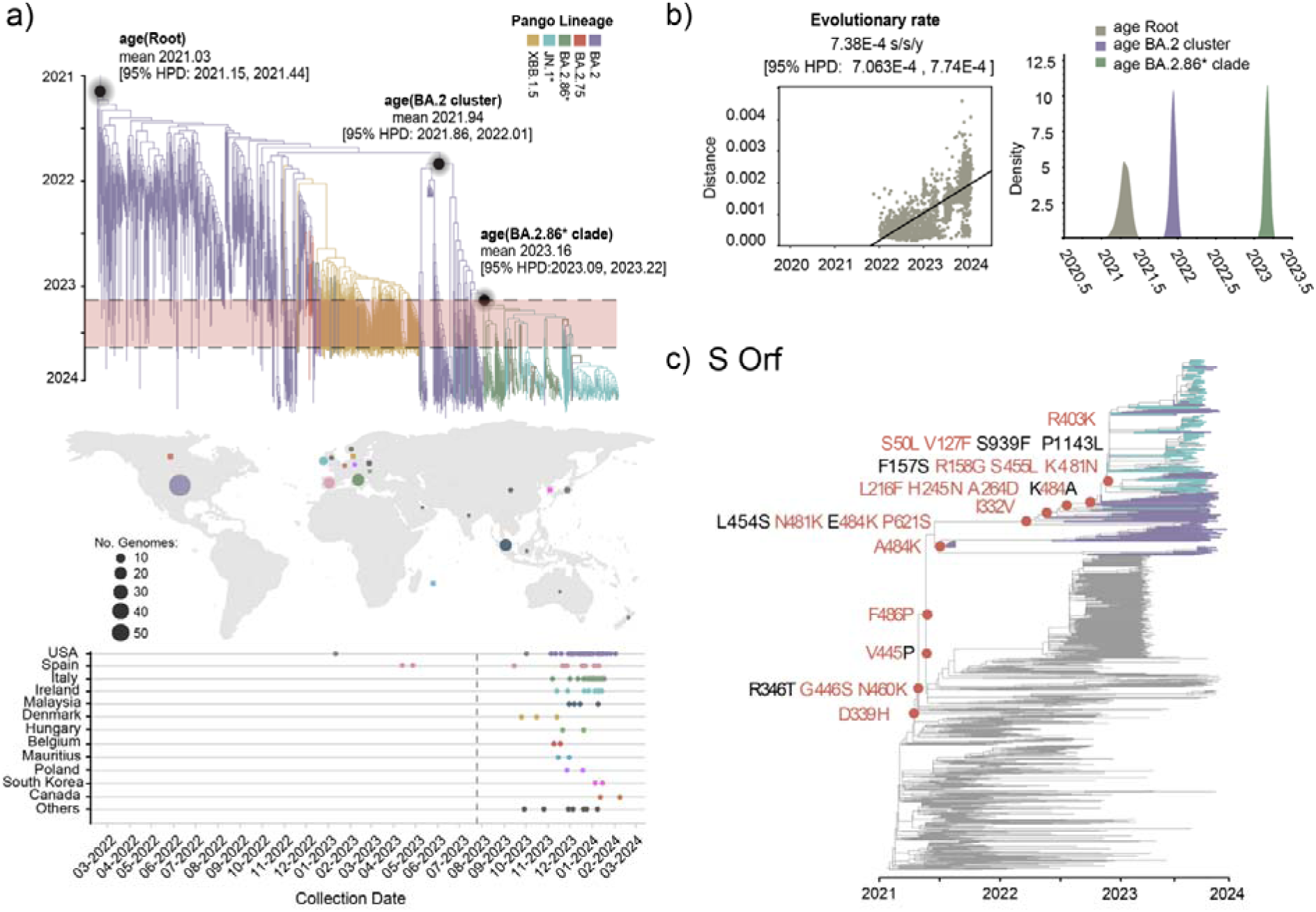
Multiple emergence and gradual evolution preceding BA.2.86. (a) The MCC tree derived from our “time-scaled” dataset, with branches coloured according to ‘Pango’ lineage. Mean node ages and 95% HPD intervals are shown for the tMRCA of the tree root, for the tMRCA of the BA.2 cluster directly ancestral to BA.2.86, and for the tMRCA of the BA.2.86* clade. The upper dashed line marks the inferred emergence date of BA.2.86 (February 2023), while the lower dashed line indicates lineage detection date (July 2023). The difference reveals a six-month lag indicative of cryptic circulation. Genomes within this cluster were sampled from multiple countries, some with collection dates preceding the detection of BA.2.86, suggesting evolution across multiple transmission chains. Sampling density and location is represented by circle sizes within the map, colour-coded by country. The temporal range of collection dates per tips is indicated below, with a dashed line marking the detection of BA.2.86. (b) The root-to-tip divergence plot indicating the inferred evolutionary rate with the corresponding 95% highest posterior density (HPD) interval. Density plots for mean node age estimates displaying the 95% HPD intervals for the tMRCA of the tree root (show in grey), the tMRCA of the cluster of BA.2.86-like intermediates (BA.2 genomes, in purple), and for the tMRCA of the BA.2.86* clade (in green). (c) Time-calibrated ML tree derived from our “time-scaled dataset” depicting the full mutational trajectory at an amino acid level inferred for the Spike protein. Branches are coloured according to ‘Pango’ lineage, highlighting only isolates within the cluster of BA.2.86-like evolutionary intermediates (in purple) and the BA.2.86* clade (in teal). Seventeen LDMs map onto deeper nodes of the tree (shown in red circles), with 13 of these observed as sequentially fixed in the cluster of BA.2.86-like intermediates, prior to the emergence of BA.2.86. Amino acid changes shown in black (R346T, L454S, F157S, S939F, P1143L) correspond to those co-occurring with LDMs, or sites showing patterns of reversion/toggling (484 and 445).

Derived from our ‘time-scaled dataset’, we inferred an overall evolutionary rate of 7.38E-4 substitutions/site/year [95% HPD: 7.063E-4, 7.74E-4], consistent with previous estimates^26,27^ (**Fig. 2b**). The mean age of the tMRCA (time to thee most recent common ancestor) at the root of the Omicron-descending BA.2* tree (excluding Wu-1) dates back to January 2021 (2021.03, 95% HPD: 2021.15, 2021.44), in line with the detection date reported for Omicron^11^. The mean age of the tMRCA for the cluster of BA.2.86-like intermediates goes back to December 2021 (2021.94, 95% HPD: 2021.86, 2022.01), whilst the mean age of the tMRCA of the BA.2.86* clade is centred around February 2023 (2023.16, 95% HPD:2023.09, 2023.22). A six-month lag between the inferred emergence and detection of BA.2.86 supports the observation of cryptic circulation (**Fig. 2b**). Under a constant virus evolutionary rate, a variant would need to accumulate at least 12 amino acid changes to remain undetected for approximately six months before being detected^4^. Consistently, the tMRCA of the cluster of BA.2.86-like intermediates indicates an emergence of around 14 months prior to the detection of BA.2.86, providing a sufficiently long timeframe to allow for a gradual accumulation of LDMs.

To investigate if gradual evolution lead to the emergence of BA.2.86, we reconstructed the full mutational trajectories for ORFs where LDMs are known to occur (ORF1a, S, M and N), mapping these onto the ‘time-scaled dataset’ (n=947) Maximum Likelihood (ML) tree (*Supplementary Figure 3*). Using the mutational trajectory of Spike as an illustrative example, we observe that multiple LDMs were sequentially acquired in BA.2.86-like intermediates prior to the emergence of BA.2.86 (**Fig. 2c**). Interestingly, some LDMs (G446S, N460K, N481K, P621S, R158G, S455L, K481N, S50L, and V127F) co-occur with other non-LDMs (R346T, L454S, F157S, S939F and P1143L), suggestive of possible epistatic interactions. Conversely, other sites, such as 484 and 445, are characterized by toggling and/or reversions. Of note, some LDMs (*e.g.,* D339H) precede the cluster of BA.2.86-like intermediates, occurring on early nodes that correspond to the distally positioned BA.2 clade, dating back to 2021. These results are consistent with a gradual accumulation of LDMs preceding the emergence of BA.2.86.

### A complex evolutionary process driving LDM emergence

To further examine the spatial and temporal distribution of LDMs over the broader evolution of all BA.2* lineages, we annotated all non-BA.2.86*-assigned genomes in our “full dataset” exhibiting at least three LDMs (*Supplementary Data 1-Supplementary Table 2*). While not directly related to BA.2.86, we identified over 2600 earlier BA.2, BA.2.75, and XBB.1.5 isolates (33.5% of genomes sampled within the “full dataset”) displaying constellations of LDMs later recognized as specific to BA.2.86. These isolates were sampled globally, with collection dates up to 19 months prior to the detection of BA.2.86 (**Fig. 3a**). When analysing LDM frequencies relative to isolate collection dates, we observed that a baseline number of LDMs (≥5) was already present in a significant proportion of genomes sampled before July 2023, followed by a rapid increase in their number after this date (**Fig. 3b**). Supporting this observation, by annotating LDMs as traits at the tips of the phylogenetic tree, we detected a consistent presence/absence pattern within a subset of these BA.2, BA.2.75, and XBB.1.5 genomes collected between January 2023 and January 2024 (*Supplementary Figure 4*). To validate our observations, assuming that a lower sequence quality could artificially inflate LDMs frequencies, we plotted mutation counts normalised by genome length against ambiguous content per genome. We observed a negative correlation between sequence quality and SAV counts (including LDMs), rather than an increase (*Supplementary Figure 5*). This indicates that the presence of LDMs were not likely driven by low sequence quality. Our findings further suggest that a fraction of LDMs were already acquired in all virus lineages distantly related to BA.2.86, and thus represent a pre-existing standing genetic variation that likely contributed to the gradual evolution of BA.2.86.

**Figure 3.**
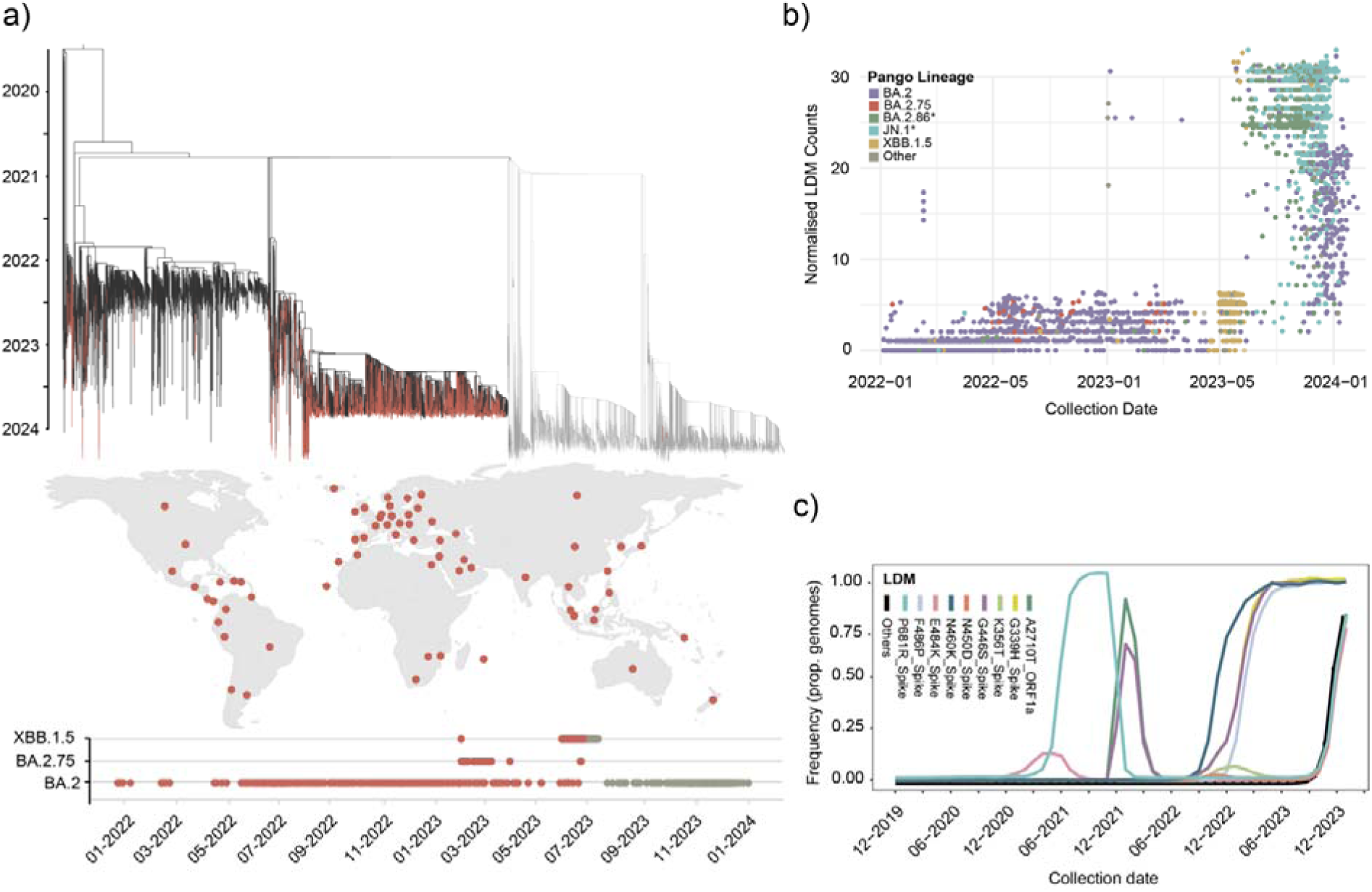
A complex evolutionary process driving LDM emergence. (a) Over 2600 earlier BA.2*-assigned genomes displaying ≥3 BA.2.86-specific LDMs are indicated as red branched on the “full dataset” time-scaled ML consensus tree. Genomes sampled after the detection date of BA.2.86 are shown in grey. The corresponding tip dates and sampling locations of isolates are shown as red dots on the map and the corresponding plot below, indicating that these genomes were collected globally and up to 19 months prior to the detection of BA.2.86. (b) Scatter plot representing normalised LDM counts relative to collection date per genome, with dots coloured according to ‘Pango’ lineage. A baseline number of LDMs (≥5) was already present in multiple BA.2*-assigned genomes collected prior to July 2023. After this date, LDM frequency increased to over 30, observed mostly in BA.2.86* genomes, but also in a considerable number of other BA.2* genomes. (c) LDM counts relative to the proportion of genomes available from GISAID from December 2019 up to March 2024 (n= ∼15,000,000, deposited as of June 2024). LDM frequencies over time are highlighted in different colours, with those displaying a similar pattern aggregated within the ‘Others’ category (shown in black). Four LDMs (A2710T in ORF1a, and P681R, E484K and G446S in Spike) show fluctuating frequencies across time, consistent with homoplasy. Four other mutations comprise an earliest constellation of LDMs (G339H, N460K, G446S and F486P in Spike) displaying a gradual emergence to reach fixation. LDMs falling within the ‘Others’ category represent a second constellation of LDMs with a frequency pattern consistent with a rapid acquisition and fixation, associated with linkage and/or recombination.

Moreover, plotting LDM frequency over the full timescale of the pandemic revealed a complex evolutionary pattern. It is evident that at least four LDMs (A2710T in ORF1a, and P681R, E484K and G446S in Spike) emerged as early as December 2020, with fluctuating frequencies across time (**Fig. 3c**). This observation is consistent with some LDMs identified as recurrent homoplasies throughout the full evolutionary history of SARS-CoV-2. Accordingly, across the global dataset available from Nextstrain (with 3821 genomes collected between December 2019 and June 2024. Accessed as of June 26^th^ 2024), we identified seven LDMs co-occurring with high frequencies in other virus lineages not directly related to BA.2.86. Specifically, we identified ORF1ab/T2710 in BA.1 (<u>Nextstrain_2710</u>), S/G158 and S/R681 in Delta (<u>Nextstrain_158</u>, <u>Nextstrain_681</u>), S/H339 and S/P486 in the XBB* lineages (<u>Nextstrain_339</u>, <u>Nextstrain_486</u>), and S/S446 and S/K460 in XBB* and other BA* lineages (<u>Nextstrain_446</u>, <u>Nextstrain_460</u>). We further identified four other mutations comprising a first constellation of LDMs (G339H, N460K, G446S and F486P in Spike) arising after June 2022 (**Fig. 3c**). These LDMs are characterized by a sequential pattern of emergence with a steady increase in frequency leading to their fixation by December 2022. This observation is consistent with gradual evolution following a soft-selective sweep^28^, in which multiple LDMs may have an additive effect on viral fitness^29^. Finally, we observed a second constellation of LDMs (exemplified by P681R, E484K in Spike, and ‘Others’), displaying a synchronous emergence after June 2023, and followed by a sudden increase in frequency leading to their eventual fixation. This pattern is consistent with a hard selective sweep, associated with the acquisition of this LDMs constellation through linkage and/or recombination.

### Evidence for recombination in the emergence of BA.2.86

As early as August 2023, it was suggested that recombination could have played a role in the emergence of BA.2.86^12^, but this hypothesis was never formally tested. To assess for recombination, we used an alignment-wide test based on topological incongruence (GARD) applied to our latest- and earliest-sampled data (n=100, reduced datasets represented by D4 and D1). In both cases, we detect consistent evidence for recombination between lineages BA.2.75 and BA.2, leading to the emergence of BA.2.86. Congruent with the reference phylogeny, the GARD Partition 1 trees show lineage BA.2.86 as directly descending from the BA.2 clade. In contrast, within the GARD Partition 2 trees, the BA.2.86 lineage is sister to clade BA.2.75 (**Fig. 4**). The topological incongruency observed is indicative of complete clade misplacement, rather than for a subset of internal branches. Both analyses revealed a single most likely breakpoint at genome site 21,314 (supported by at least four different models. Δ c-AIC vs the null model: 2306.91. Δ c-AIC vs the single tree multiple partition: 2392.67) (*Supplementary Data 8 and 9*). Thus, our finding, together with a synchronous emergence and sudden increase in frequency observed for a second constellation of LDMs, supports that recombination likely played a role in the emergence of BA.2.86. Recombination also provides a plausible explanation for how LDMs suddenly expanded after July 2023.

**Figure 4.**
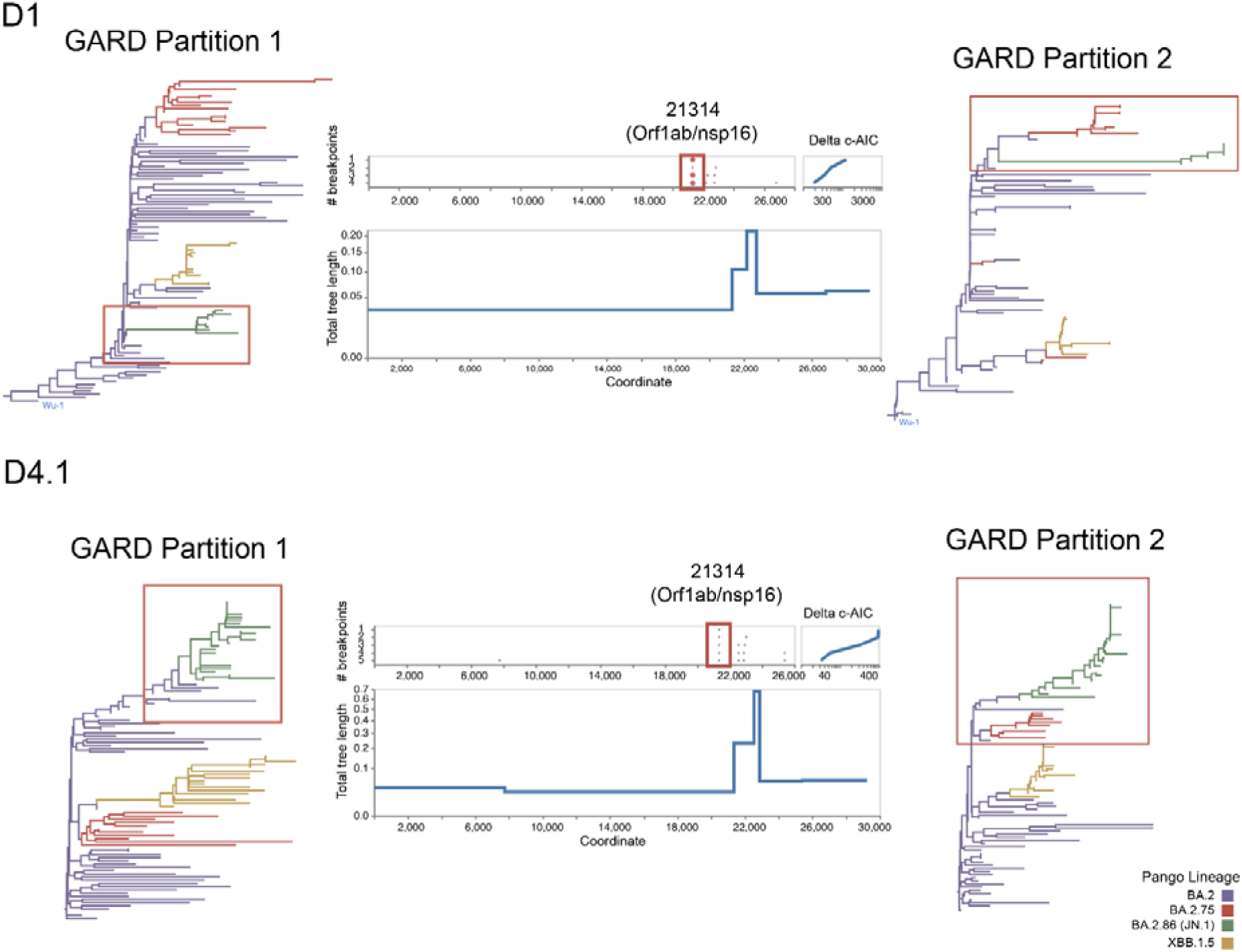
Evidence of recombination in the emergence of BA.2.86. Recombination breakpoints inferred from alignment-wide testing associated with topological incongruence (GARD), applied to both the D4.1 (later sampled) and D1 (earlier sampled) reduced datasets. A consistent topological incongruency results in two partition trees (GARD Partition tree 1 and GARD Partition tree 2), shown with branches coloured according to ‘Pango’ lineage. Consistent with the reference phylogeny, in the GARD Partition 1 tree, BA.2.86* (in green) directly descends from the parental BA.2 (in purple). In contrast, in the GARD Partition 2 trees, BA.2.86 is a sister lineage of BA.2.75 (in red), both descending from BA.2. A single best supported breakpoint across the alignments was identified at codon site 21314 in ORF1ab (highlighted with red boxes). Site 21314 corresponds to residue I6994 within ORF1ab, located in a loop connecting a beta-sheet and an alpha-helix in nsp16 (motif PKEQIDGY). This residue is positioned distally from known inhibitor binding sites, with recombination at this point unlikely to disrupt protein function.

Due to recurrent homoplasy and a high genetic similarity between closely related lineages, identifying recombination-associated mutation patterns has remained challenging in SARS-CoV-2^30^. In the case of BA.2.86, this is likely confounded by preceding recombination events amongst related parental lineages^31^. Thus, to find recombination-associated mutation patterns, for all lineages of interest (BA.2, BA.2.75, XBB.1.5, BA.2.86), we identified all ‘private mutations’ (defined as unique to a given lineage, but not observed in the nearest phylogenetically related clade^32^) and excluded those observed as homoplasy (shown in black in *Supplementary Table 2*). Then, to detect the subset of LDMs contributed by each parental lineage, we mapped full mutational trajectories onto deeper nodes of each partition tree (*Supplementary Figure 6*), cross-referencing these with the abovementioned list of ‘private mutations’ (*Supplementary Table 2*). We identify two LDMs as unique contributions by each parental lineage, namely A6183G in ORF1a (K1973R, NSP3:K1155R) contributed by BA.2 (as in the cluster of BA.2.86-like intermediates), and C22033A (S:F157L) in Spike contributed by BA.2.75. For the latter, given the degeneracy of the genetic code, we presume that the directional F→L→S at site 157 change would be the most parsimonious explanation for L157S to be fixed in BA.2.86. Supporting this observation, state F157 is present in >90% of all BA.2*-assigned genomes (excluding BA.2.75), and state L157 is present in >90% of BA.2.75 genomes (<u>Nextstrain 157</u>). Consistent with the inferred recombination breakpoint, the LDMs identified either locate to ORF1a or to the Spike ORF (*Supplementary Figure 6, Supplementary Table 2*).

## DISCUSSION

We tested the hypothesis that the highly divergent BA.2.86 variant emerged through a single event following accelerated evolution within an individual chronic infection case. Through our analysis, we identify a cluster of approximately 100 BA.2 genomes consistently positioned as directly ancestral to BA.2.86, and effectively shortening the phylogenetic branch leading to the clade. These isolates also show a variable presence pattern of multiple BA.2.86-specific LDMs, rendering these as putative evolutionary intermediates. Following a detailed investigation on the sequencing quality scores and mutation frequencies of a subset of these assemblies, both genetic and evolutionary evidence suggests that at least a fraction of these isolates represent genuine evolutionary intermediates. Together, our findings indicate that BA.2.86 is unlikely to have emerged from a single chronic infection. Instead, we find evidence supporting that the emergence of this variant was most probably the result of a complex evolutionary process, including a gradual accumulation of mutations, evolution across multiple transmission chains, and recombination derived from a standing genetic variation, without excluding the possible contribution of chronic infection cases.

We further propose a two-step evolutionary process to explain the emergence of LDMs that reached fixation in BA.2.86. First, we note that the earliest LDMs arose almost two years before the detection of BA.2.86, observed in SARS-CoV-2 lineages not directly related to BA.2.86 (hence homoplasic across the full evolutionary history of the virus). Subsequently, and in an independent manner, a primary constellation of LDMs emerged in the Omicron-descending BA.2* lineages over a year before the detection of BA.2.86. These LDMs reached fixation through gradual evolution, observed mostly within the cluster of BA.2.86-like intermediates. Following this, and approximately six months prior to the detection of BA.2.86, a secondary constellation of LDMs likely emerged through recombination between the parental BA.2 and BA.2.75 lineages, resulting in a (partial) burst of mutations later observed in BA.2.86. These observations underscore the complexity of the processes shaping the evolution of this highly divergent SARS-CoV-2 variant. Moreover, based on the evidence of recombination we find, we further suggest that BA.2.86 should be reclassified as an “X” recombinant lineage within the ‘Pango’ nomenclature^10^.

Evolutionary intermediates have been previously reported, albeit to a lesser extent, in the emergence of other highly divergent SARS-CoV-2 variants. Although controversy surrounds their detection — particularly in the context of the emergence of the Omicron variant^33^ — there is limited but credible evidence for true evolutionary intermediates preceding other VOCs. For instance, for the Alpha and Gamma variants, independent studies reveal a handful of intermediate genomes showing an early acquisition of constellations of mutations through multiple interhost transmission chains, with cryptic circulation of endemic lineages prior to VOC detection^34–36^. Thus, our findings largely align with and extend prior observations, providing additional support for the role of intermediate precursors in the evolution of some highly divergent SARS-CoV-2 lineages.

Disparities in sequencing capacity across regions can hinder the detection of critical stages in the evolution of highly divergent SARS-CoV-2 variants^8^. For example, Omicron was initially identified in South Africa and Botswana, areas displaying a robust genomic surveillance relative to other regions in Africa, and was suggested that this variant could have circulated undetected in other countries. Prior to our study, and in line with our results, an earlier report had already suggested that the closest ‘antigenically related’ isolates to BA.2.86, correspond to BA.2 strains sampled from South Africa circulating as early as 2022^37^. Nonetheless, identifying emerging SARS-CoV-2 variants before they reach epidemiological success becomes near-impossible solely based on genomic surveillance records. Novel virus lineages emerge constantly, and the spread of precursors may not stand out during early stages of evolution. Thus, monitoring variant emergence may be more effective as an early warning system if genomic surveillance is combined with viral fitness prediction tools based on the viral genetic make-up^23,38^.

Although our results do not support the emergence of BA.2.86 linked to a single long-term infection case, it remains highly likely that SARS-CoV-2 evolution in chronically infected patients has contributed to shaping the genetic makeup of emerging variants. Notably, within the cluster of BA.2.86-like intermediates, we identify two isolates from Spain that derive from the same patient, with individual samples taken weeks apart from a single chronic infection ^Footnote 1^ (*Supplementary Text 1*). Additionally, though not directly related to BA.2.86, we also identified four earlier BA.2 isolates from Mexico deriving from a single patient ^Footnote 2^. In both instances, assemblies are not clonal, and display a differential presence/absence pattern of LDMs, highlighting the role of chronic infections in enabling the rapid acquisition of mutations^5,6^. While chronic infections are expected to represent a small fraction of SARS-CoV-2 infections (0.1–0.5%)^39^, both chronic and non-chronic infections likely contribute to a standing genetic variation at an intra-host level, setting the stage for subsequent fixation at a population level^23^. Nonetheless, with an underestimated prevalence and a limited reporting of these cases within our data (and in GISAID in general), we are unable to draw further conclusions on how chronic infection may have contributed to the evolution of BA.2.86.

Our results are incompatible with a scenario in which the BA.2.86 variant emerged through a single event linked to an individual long-term infection. Instead, our analyses reconstruct a complex evolutionary process, with a gradual accumulation of mutations likely involving multiple transmission chains, a cryptic circulation of precursor genomes, and recombination derived from an extant standing variation. Whether the evolutionary dynamics and patterns we describe here can be extrapolated to the emergence of other highly divergent SARS-CoV-2 variants remains an open question. Though, by elucidating these dynamics, we provide a valuable framework for investigating the origins and trajectories of other divergent viral lineages.

## METHODS

Capturing the emergence and diversification of BA.2.86 up to May 2024, we collated four independent datasets (named here D1, D2, D3, D4) comprising high-quality, complete SARS-CoV-2 genomes (low-coverage excluded, Ns ≤ 5%) from GISAID^18–21^. Datasets were retrieved at different time points and/or sequentially subsampled using a phylogenetically-informed approach. Subsequently, we applied a suite of evolutionary analyses, including high-resolution phylogenetic inference, ancestral state reconstructions and node dating under time-scaled analyses, as well as testing for recombination.

### Dataset curation

D1 (n=100, compiled as of August 21^st^ 2023) corresponds to the original dataset derived from the initial characterisation of BA.2.86 made available through the Nextstrain platform^12,40^. D1 includes the eight first detected BA.2.86 genomes, and a set of 92 genomes assigned to the parental/most closely related virus lineages: BA.2, BA.2.75, and XBB.1.5 (*Supplementary Data 1-Supplementary Table 3; Supplementary Figure 1a*). BA.2.86 genomes correspond to those first identified in Denmark, UK, USA, and Israel as of August 14^th^ 2023 ^12^: EPI_ISL_18097345, EPI_ISL_18125259, EPI_ISL_18125249, EPI_ISL_18097315, EPI_ISL_18096761, EPI_ISL_18110065, EPI_ISL_18114953, EPI_ISL_18111770. D1 was used to confirm the phylogenetic placement of BA.2.86 relative to its parental/most closely related virus lineages, and to confirm our observations on recombination derived from analysis of D4 (see Methods section *‘Detection of Recombination’*).

D2 (n=9134, including Wu-1 NC_045512 for rooting purposes) was compiled as of January 9^th^ 2024, and considers an expanding diversity of the BA.2.86*/JN.1* lineages (labelled as ‘Foreground’ or FG) sampled within six months following the detection of BA.2.86 (*Supplementary Data 1-Supplementary Table 4; Supplementary Figure 1b*). To generate D2, 35,902 genomes assigned as BA.2.86* were retrieved. Drawing from a preliminary neighbour-joining tree, genomes were subsampled to 6847 to include a representative diversity of the BA.2.86* clade. D2 also includes an extended sampling (n=2282) of other BA.2, BA.2.75, and XBB.1.5 genomes (labelled here ‘Background’ or BG). D2 was used to infer a preliminary high-resolution phylogeny capturing the evolution of the BA.2.86* clade.

D3, or *“full dataset”* (n=7760) was compiled as of March 15, 2024. D3 was generated based on the initial identification of a cluster of BA.2 genomes directly ancestral to the BA.2.86* clade (BA.2.86-like intermediates) in D2. As placement could be biased by the over-representation of the BA.2.86*/JN.1* and XBB.1.5* lineages, we sought to generate an updated dataset achieving an extensive and balanced taxonomic representation (*Supplementary Data 1-Supplementary Table 5; Supplementary Figure 1c*). To further capture a potentially undersampled virus diversity of BA.2 genomes directly ancestral to BA.2.86, we mined the GISAID database to retrieve all BA.2*-assigned genomes displaying each of the BA.2.86-specific LDMs at the nucleotide level^13^ (n=15,607, labelled as ‘Mutated’ or MT). Genomes within D3 were re-assigned to their corresponding virus lineage prior to subsequent analyses using Pangolin v4.3 (PUSHER-v1.26, Scorpio v0.3.19, and Constellation v0.1.12)^41^ (*Supplementary Data 1-Supplementary Table 8*). After correcting for ‘Pango’ lineage misassignment, only a small fraction of genomes (<1.5%, 110/7760) displayed assignment conflict, with only seven of these corresponding to the evolutionary intermediates identified (*Supplementary Data 5*). Following this step, a new set of foreground (n= 2706) and background genomes (n= 5053) was subsampled in equal numbers (∼750 genomes per lineage). To confirm an overall phylogenetic placement, we further resampled a replica of D3 to reconstruct a second phylogenetic tree, retaining only those genomes showing a consistent phylogenetic placement in both datasets (*Supplementary Data 6*).

D4 or “*time-scaled dataset”* (n=947) was compiled as of May 22^nd^ 2024. D4 comprises an updated and quality-controlled down-sampled version of D3, filtering out sequences with incomplete collection dates and with a genome ambiguity content >0.1 (defined as the proportion of content that is *not A, T, C, G, or -* relative to genome completeness) (*Supplementary Data 1-Supplementary Table 6; Supplementary Figure 1d*). Additionally, potentially misdated genomes identified as outliers within the root-to-tip divergence plots generated using TempEst v1.5.3 ^42^ were further removed. This resulted in a streamlined dataset that enabled subsequent computationally costly analyses (*Supplementary Data 7*).

D4.1 or “*GARD dataset”* (n=100) comprises an *in-frame* and reduced version of D4 generated to enable a genome-wide recombination testing based on phylogenetic incongruency (GARD) (see Methods section *‘Detection of Recombination’*). To generate D4.1, we used an *in-house* script to partition D4 into sub-alignments comprising the *in-frame* open reading frames of all 11-known SARS-CoV-2 genes, verifying ORF identify using BLAST^43,44^. Individual ORF alignments were then concatenated to generate an “all ORF” *in-frame* alignment, further subsampled to 100, retaining a representative diversity relative to D1 (*Supplementary Data 1-Supplementary Table 7; Supplementary Figure 1e; Supplementary Data 8 and 9*).

### Alignment construction and iterative subsampling and phylogenetic inference

Individual datasets were initially mapped against the SARS-CoV-2 reference genome (Wu-1, NC_045512) using minimap2-2.28^45^ under default settings. Following UTR trimming, datasets were then re-aligned using MAFFT v7.520^46^ (mafft --auto --thread 8). The resulting alignments were subsequently inputted for phylogenetic inference under a Maximum Likelihood (ML) framework using IQ-TREE multicore version 2.2.6 ^47^ (iqtree2 -nt 8 -m GTR+I+G -B 1000 -keep-ident). Following this, applied to D3-D4.1, we used a phylogenetically-informed subsampling approach, carried out as an iterative process using Treemmer v0.3^48^. Phylogenetic congruency (*i.e.,* a consistent placement for all virus lineages of interest) was verified for all output trees through comparison to the global SARS-CoV-2 phylogeny available from Nextstrain ^2,3^, updated between August 2023 and August 2024.

### Reconstruction of Ancestral States

The definition of LDMs used in this study includes those mutations listed as ‘specific’ to BA.2.86, displaying a prevalence of >75% only within this lineage (defined as of August 2023 ^13^ in <u>outbreak.info</u>, with the site accessed as of August 2023). Only non-synonymous mutations (*i.e.,* resulting in amino acid changes) were considered, as these are expected to contribute to evolutionary processes linked to adaptation. To investigate the occurrence pattern of BA.2.86 specific-LDMs ^13^, corresponding mutations/sites were identified and extracted at an amino acid level from the alignments, coded as discrete traits, and mapped onto all nodes of the D2-D4 trees. The definition of deeper nodes considers those occurring at the backbone of the phylogeny, including the most recent common ancestor (MRCAs) of major clades (*e.g.,* virus lineages of interest). Optimizing computational costs relative to dataset size, we opted for a ML framework. For reconstructing site-specific mutational trajectories, we used the *mugration* model (*treetime mugration --aln X.fasta --tree X.nwk --states LDM.csv --attribute X –confidence)* in TreeTime v0.11.3^49^. Within D3, we further annotated all genomes with early LDM constellations, defined as those displaying a co-occurrence of at least three LDMs, verified using the original metadata sheets. Genomes meeting this criterion were annotated within the consensus D3 phylogeny and visualized using <u>FigTree</u> V1.4.4. Applied to D4, we then reconstructed full mutational trajectories for ORFs where LDMs are known to occur. This was done using the *ancestral* model *(treetime ancestral --tree X.nwk --aln X_aa.fasta --aa)* in TreeTime v0.11.3^49^, inputting the individual ORF1ab, Spike, M and N codon alignments translated to amino acids. Finally, the D3 and D4 trees were time-calibrated informed by tip dates using TreeTime^49^ *(treetime --aln X.fasta --tree X.nwk -- dates X.csv --keep-root --stochastic-resolve --clock-rate 8e-4 --clock-filter 0)*.

### Time-scaled analysis

The D4 ML time-calibrated phylogeny was used as a starting tree for a time-scaled Bayesian analysis using the BEAST software package v1.10.4^50^. We used a customized set of tree operators to maintain a fixed topology while re-estimating the root and internal node heights. To ensure robust age estimates, potential tip misdating observed within the cluster of BA.2.86-like intermediates (*i.e.,* samples Malaysia with collection date of 2022-03-11: EPI_ISL_18821487, EPI_ISL_18821492, EPI_ISL_18821480, EPI_ISL_18821490, EPI_ISL_18821494, EPI_ISL_18821486, EPI_ISL_18821491, EPI_ISL_18821483, EPI_ISL_18821484, EPI_ISL_18821485) (*Supplementary Text 1)* was accounted for by adding a sampling uncertainty of ± 2 years, as informed by collection and submission dates available from GISAID metadata. Analysis was performed under the HKY substitution model, a strict clock model, and an exponential growth tree model. Drawing from existing literature^26,27^, we used an informative clock prior under a log-normal distribution centred on a mean of 8.0E-4, and with a standard deviation of 1.0E-4. All other priors were left to default settings. We ran three independent MCMC chains with length of 10E9, sampling every 1E5 states. For all chain, convergence was assessed using Tracer v.1.7^51^, evaluating an effective sample size (ESS)D>D200 for key parameters. Outputs were combined into a single chain of 2.073E8 using LogCombiner v1.10, removing the initial 1% as burn-in. Finally, a ‘Maximum clade credibility’ tree was summarized using TreeAnnotator v.1.10.4.

### Detection of Recombination

To identify genome-wide recombination breakpoints associated with phylogenetic incongruency, we employed the Genetic Algorithm Approach for Detection of Recombination (GARD)^52,53^ (*Supplementary Data 8 and 9*). Due to its high computational demand, GARD is recommended for datasets of 100 sequences or fewer, and thus was applied to the reduced D1 and D4 datasets. Implemented as part of the HyPhy package v2.5.48^52–54^, we ran GARD using the following command line: hyphy gard --alignment X.fasta --rv Gamma --rate-classes 4 --max-breakpoints 10 - -mode Faster ENV=TOLERATE_NUMERICAL_ERRORS=). Additionally, to detect individual recombinant sequence, we employed RDP4 (Recombination Detection Program) under default settings^55^, retrieving negative results. Derived from the analysis of D4.1, and following the same approach described above, the full mutational trajectory at an amino acid level corresponding to each partition alignment was mapped onto deeper nodes of each of the partition tree (*Supplementary Figure 6*).

### Assembly quality control

Following the initial release of this study as a preprint on July 2024^56^,concerns were raised that some of the evolutionary intermediates identified derived from low-quality sequencing libraries, or represented assemblies generated from contamiation/mixed infections (*Supplementary Text 1)*. This prompted us to run extensive quality checks on a fraction of these assemblies based on their raw short-read sequencing data. For this purpose, we downloaded data available for 16 SRA assemblies corresponding to the cluster of BA.2.86-like intermediates, identified by linking individual GISAID identifiers to a matching BioSample record. In a further attempt to obtain more sequencing libraries, we contacted over 50 originating labs that had submitted these assemblies. We obtained response from 9 of these, from which 7 shared short-read data directly, and 2 provided ARTIC sequencing reports (see Acknowledgements section and *Supplementary Text 1*) (*Supplementary Data 1-Supplementary Table 1,* assemblies with available short-read data are highlighted in yellow, assemblies with available ARTIC sequencing reports are highlighted in red). This resulted in a total of 23 sequencing libraries that could be assessed. For comparative purposes, we set to establish a background distribution of ‘high-quality’ raw data corresponding to a random subsample of BA.2.86-assigned assemblies displaying a consistent phylogenetic placement. For this, we retrieved 901 publicly available SRA datasets, following the inclusion criteria of: only complete consensus genomes (excluding proviral, laboratory, vaccine, and environmental strains), BioSample records linked to a single genome, and libraries generated only using either Illumina paired-end or Oxford Nanopore sequencing technologies. Data was further curated based on a random selection of one library per country, collection date, and sequencing platform, resulting in a total of 599 libraries generated through Illumina, and 302 generated through Nanopore sequencing technologies, representing assemblies from Germany, Japan, Saudi Arabia, Senegal, Slovakia, USA, and the UK, with collection dates ranging from June 30^th^ 2023, to August 24^th^ 2024.

All data, whether shared by originating labs or retrieved directly from NCBI, was processed using the same pipeline. SRA libraries were retrieved using the NCBI’s *entrez-direct* command-line API v21.6. For both the BA.2 and background sequencing libraries, short-read data was converted to FASTQ format using the *prefetch* and *fastq-dump* commands from NCBI’s sra-tools API v3.11 to be further processed. First, sequencing adapters and low-quality read ends were removed, and read pairs with a mean base quality of <Q20 were filtered out using the *BBDuk.sh* utility from the BBMap v39.06. package (https://sourceforge.net/projects/bbmap/). Quality-filtered reads were then mapped against the human CHM13 reference genome (accession: GCF_009914755.1) using Bowtie2 v2.5.1 ^57^, discarding all mapped pairs. Retained pairs were subsequently mapped against the Wu-1 reference genome, keeping those for which both reads mapped to the reference. Duplicate reads were further removed using the *markdup* utility from Samtools v1.20^58^. Major/minor allele (MAF) and Single Amino acid Variant (SAV) frequencies were estimated from the de-duplicated, high-quality data using the *call codonvar* utility in Quasitools v0.7.0 (https://phac-nml.github.io/quasitools/), specifying a sequencing error rate corresponding to Q20 (‘*--error_rate 0.01*’). We plotted mean quality scores (base and mapping quality, mean sequencing depth) and SAV counts for the BA.2 sequencing libraries relative to the background distribution. Under the assumption of a positive correlation between genome sequence quality and mutation counts, we further plotted normalised mutation counts relative to ambiguous content per genome.

### Dataframes, Plots and Statistical Analyses

All metadata was extracted from the GISAID metadata sheets (see DATA AVAILABILITY, GISAID EPI_SET IDs). Plots and logistic regression analyses generated using the “stats” (https://rdrr.io/r/stats/stats-package.html) and “ggplot2” (https://cran.r-project.org/web/packages/ggplot2/index.html) R packages. Tables were created in Microsoft Excel 2018.

## DATA AVAILABILITY

EPI_SET IDs for D1-D4 are as follow:

D1 EPI_SET ID: EPI_SET_240717hn doi: 10.55876/gis8.240717hn D2 EPI_SET ID: EPI_SET_240717ps doi: 10.55876/gis8.240717ps

D3 EPI_SET ID: EPI_SET_240717xg doi: 10.55876/gis8.240717xg

D4 EPI_SET ID: EPI_SET_240717xr doi: 10.55876/gis8.240717xr

Unique identifiers are used for both the acknowledgment of data contributors and to permit registered users to retrieve the dataset of all the records encompassed in the EPI_SET ID. The resulting GARD json outputs (*Supplementary Data 8 and 9*) can be visualized using http://vision.hyphy.org/.

## ACKNOWLEDGMENTS

M.E.Z, F.B, L.vD and this project are part of the grant HORIZON-HLTH-2021-CORONA-01-02 agreement no. 101046314, Funded by the European Union. Views and opinions expressed are however those of the author(s) only and do not necessarily reflect those of the European Union or the European Health and Digital Executive Agency. Neither the European Union nor the granting authority can be held responsible for them. M.E.Z is funded by a UCL Rosetrees Excellence Fellowship UCL2024\2. L.vD is further funded by a Future Leaders Fellowship MR/X034828/1. C.C.S.T. is funded by the National Science Scholarship from the Agency for Science, Technology and Research (A*STAR), Singapore. We would like to acknowledge Pooja Swali and Alexei Yavlinsky for their support in dataframe construction and visualization. Additionally, we would like to thank all originating labs who kindly shared their raw short read data and case-specific insights with us. Particularly, we acknowledge representatives of the:

1. Monterey County Public Health Laboratory, USA
2. Lab voor klinische biologie, Belgium
3. Department of Clinical Microbiology, GIGA Medical Genomics, Belgium
4. Hospital of Southern Norway - Kristiansand, Department of Medical Microbiology. Norwegian Institute of Public Health, Department of Virology, Norawy
5. Virginia Tech. Virginia Division of Consolidated Laboratory Services, USA
6. Servicio de Microbiologia Hospital Ramon y Cajal, Spain
7. Servicio de Microbiologia, Hospital Universitario Virgen del Rocio, Sevilla, Spain. Fundación Progreso y Salud Computational Medicine Platform, Spain
8. Waikato DHB Laboratory Institute of Environmental Science and Research (ESR), New_Zealand
9. Max von Pettenkofer Institute, Virology, National Reference Center for Retroviruses, LMU Munich, Germany

## Footnotes

1 Information retrieved from the originating lab. EPI_ISL_17630096, EPI_ISL_17797704. Listed in *Supplementary Data 1-Supplementary Table 1*.

1 Information retrieved from the originating lab. EPI_ISL_18415854, EPI_ISL_18798234, EPI_ISL_18798204 and EPI_ISL_1841583. Listed in *Supplementary Data 1-Supplementary Table 2*.

