## Supplementary Information for "Early evolution of BA.2.86 sheds light on the origins of highly divergent SARS-CoV-2 lineages"

#### Supplementary Text 1. Extended Discussion

Supplementary Table 1. LDMs presence/absence in 11 genomes from Malaysia

#### Supplementary Figures

Supplementary Figure 1. A consistent phylogenetic placement with a BA.2 cluster directly ancestral to BA.2.86
Supplementary Figure 2. Quality scores and SAV counts for BA.2 assemblies fall within the typical range of other sequencing libraries
Supplementary Figure 3. The mutational trajectory of LDMs showing gradual evolution in the BA.2 cluster directly ancestral to BA.2.86
Supplementary Figure 4. Occurrence of LDM across non-BA.2-86* genomes
Supplementary Figure 5. Normalized mutation counts against ambiguous content per genome
Supplementary Figure 6. Mutational trajectory inferred on P1 and P2 recombination trees

#### Supplementary Tables

Supplementary Table 2. LDMs acquired through recombination

#### Supplementary Files

Supplementary Data 1- Supplementary Tables

- Supplementary Table 1. BA.2 genomes representing evolutionary intermediates leading to the BA.2.86* clade
- Supplementary Table 2. Early BA.2* genomes with ≥ 3 LDMs
- Supplementary Table 3. Metadata and GISAID EPI_ISL for genomes in D1
- Supplementary Table 4. Metadata and GISAID EPI_ISL for genomes in D2
- Supplementary Table 5. Metadata and GISAID EPI_ISL for genomes in D3
- Supplementary Table 6. Metadata and GISAID EPI_ISL for genomes in D4
- Supplementary Table 7. Metadata and GISAID EPI_ISL for genomes in D4.1
- Supplementary Table 8. Assemblies displaying conflict during Pango reassignation

Supplementary_Data_2_LDMs-RASML.pdf
Supplementary_Data_3_Major_AlleleFrequencies.pdf
Supplementary_Data_4_Minor_AlleleFrequencies.pdf
Supplementary_Data_5_Pango_Lineage_Reassignation.xlsx
Supplementary_Data_6_D3.tree
Supplementary_Data_7_D4.tree
Supplementary_Data_8_D1_GARD.json
Supplementary_Data_9_D4_1_GARD.json

#### Supplementary Text 1. Extended Discussion

##### Notes on the 'Malaysia' genomes

Within the cluster of BA.2 identified as BA.2.86-like intermediates, we identified 11 genomes from Malaysia with collection dates preceding the detection of BA.2.86 (*Supplementary Data 1-Supplementary Table 1).* These 'Malaysia' genomes were all collected on 2022-03-11, sequenced by the same originating lab. We noted that, despite temporal and regional clonality, these assemblies are not genetically identical, displaying a variation in branch length within the phylogenetic trees, and a differential presence/ absence pattern of BA.2.86-specifc LDMs (*Supplementary Table 1*). Nonetheless, their collection date poses a two-year difference compared to their submission date, flagging these as putative contaminants. Further investigations shared with us through personal communication show that, in addition to displaying multiple LDMs, these 'Malaysia' genomes also feature a deletion characteristic of BA.2.86* genomes sampled around November 2023 (>95%, Δ23008-230210, resulting in the loss of residue 484 in the Spike protein) (*Supplementary Table 1*). Two of the 'Malaysia' genomes (EPI_ISL_18821484 and EPI_ISL_18821485) also share mutations G19677T and G22578A, associated with a 'Malaysia-endemic' lineage (BA.2.40, with >88% mutation frequency), circulating within the region during March 2022. Within a broader phylogenetic context, the 'Malaysia' cluster is also closely related to other Malaysian isolates collected as of 2022. As short-read data for these assemblies was unavailable to validate quality scores and mutation counts, we cannot rule out possible cross-contamination. Although these 'Malaysia' genomes were not identified as outliers in the root-to-tip divergence plots (**Fig. 2b, main text**), uncertainty in their sampling date was accounted for (±2 years) within our time-scaled analysis, and we do not consider these assemblies in our results.

##### Notes on other genomes with earlier collection dates

We also identify three other genomes with confirmed collection dates predating the detection of BA.2.86 (*Supplementary Data 1-Supplementary Table1*). For isolate EPI_ISL_16731648, sampled in the USA on 2023-01-15, short-read data was retrieved, with our MAF analyses revealing no autocorrelation in allele frequencies counts. Moreover, all quality scores and mutation counts fall within the typical range of other high-quality sequencing libraries (**Fig. 2c, main text**), (*Supplementary Figure 2, Supplementary Data 4 and 5*). This, together with confirmed metadata accuracy (as information retrieved through direct communication with the originating lab), we can rule out contamination contributing to the phylogenetic placement and mutation patterns observed for this isolate (see Results section *'Quality Control'*). Additionally, we were able to recover the ARTIC sequencing reports for two other earlier genomes sampled from Spain (EPI_ISL_17630096, EPI_ISL_17797704, with collection dates of 2023-04-25 and 2023-05-26, respectively), for which short read data was no longer available from the originating lab (*Supplementary Data 1-Supplementary Table 1, highlighted in red*). Again, these assemblies show high-quality scores, sequenced with an average depth of 331x, and with 91% coverage at >100x. Both reports include a single warning regarding an increased number of 'private mutations' (defined as mutations unique to these samples, yet not observed in their nearest neighbour within the Nextclade reference tree) [1]. Nonetheless, no 'labelled mutations' were identified (shared mutations with those common to specific clades), often regarded as a sign of contamination, coinfection or recombination[1]. Again, these assemblies display variations in branch lengths and a differential presence/ absence pattern of LDMs. Moreover, we know that these isolates represent samples from the same patient in the context of a chronic infection (see **Discussion, main text**).

2 GitHub. *cov-lineages/pango-designation #2183*, 2024).

#### Supplementary Table 1. LDMs presence/absence in genomes from Malaysia

| | A211D | V1056L | N2526S | A2710T | V3593F | S50L | V127F | R158G | L216F | H245N | A264D | I332V | D339H | K356T | R403K | V445H | G446S | N450D | L452W | N460K | N481K | A484K | F486P | R493Q | E554K | A570V | P621S | H681R | D3H | T30A | A104V | Q229K |
|---|---|---|---|---|---|---|---|---|---|---|---|---|---|---|---|---|---|---|---|---|---|---|---|---|---|---|---|---|---|---|---|---|
| Consensus BA.2.86* | D | L | S | T | F | L | F | G | F | N | D | V | H | T | K | H | S | D | W | K | K | K | P | Q | K | V | S | R | H | A | V | K |
| Consensus early BA.2 | A | V | N | A | V | S | V | R | L | H | A | I | G | K | R | V | S | N | L | N | N | E | F | Q | E | A | P | H | D | T | A | Q |
| EPI_ISL_18821487 | A | V | S | A | V | S | V | R | L | H | A | I | D | T | K | H | S | D | W | K | K | Δ | P | Q | K | V | S | R | D | T | A | Q |
| EPI_ISL_18821492 | A | V | S | A | V | S | V | R | L | H | A | I | D | T | K | H | S | D | W | K | K | Δ | P | Q | K | V | S | R | D | T | A | Q |
| EPI_ISL_18821480 | A | V | S | A | V | S | V | R | L | H | A | I | D | T | K | H | S | D | W | K | K | Δ | P | Q | K | V | S | R | D | T | A | Q |
| EPI_ISL_18821490 | A | V | S | A | V | S | V | R | L | H | A | I | D | T | K | H | S | D | W | K | K | Δ | P | Q | K | V | S | R | D | T | A | Q |
| EPI_ISL_18821494\| | A | V | S | A | V | S | V | R | L | H | A | I | D | T | K | H | S | D | W | K | K | Δ | P | Q | K | V | S | R | D | T | A | Q |
| EPI_ISL_18821486 | A | V | N | A | V | S | V | R | L | H | A | I | D | T | K | H | S | D | W | K | K | Δ | P | Q | K | V | S | R | D | T | A | Q |
| EPI_ISL_18821491 | A | V | N | A | V | S | V | R | L | H | A | I | D | T | K | H | S | D | W | K | K | Δ | P | Q | K | V | S | R | D | T | A | Q |
| EPI_ISL_18821483 | A | V | S | A | V | S | V | R | L | H | A | I | D | T | K | H | S | D | W | K | K | Δ | P | Q | K | V | S | R | D | T | A | Q |
| EPI_ISL_18821484 | A | V | N | A | V | S | V | R | L | H | A | I | D | T | K | H | S | D | W | K | K | Δ | P | Q | K | V | S | R | H | A | A | Q |
| EPI_ISL_18821485 | A | V | S | A | V | S | V | R | L | H | A | I | D | T | K | H | S | D | W | K | K | Δ | P | Q | K | V | S | R | H | A | A | Q |

**Supplementary Figure 1. A consistent phylogenetic placement with a BA.2 cluster directly ancestral to BA.2.86**

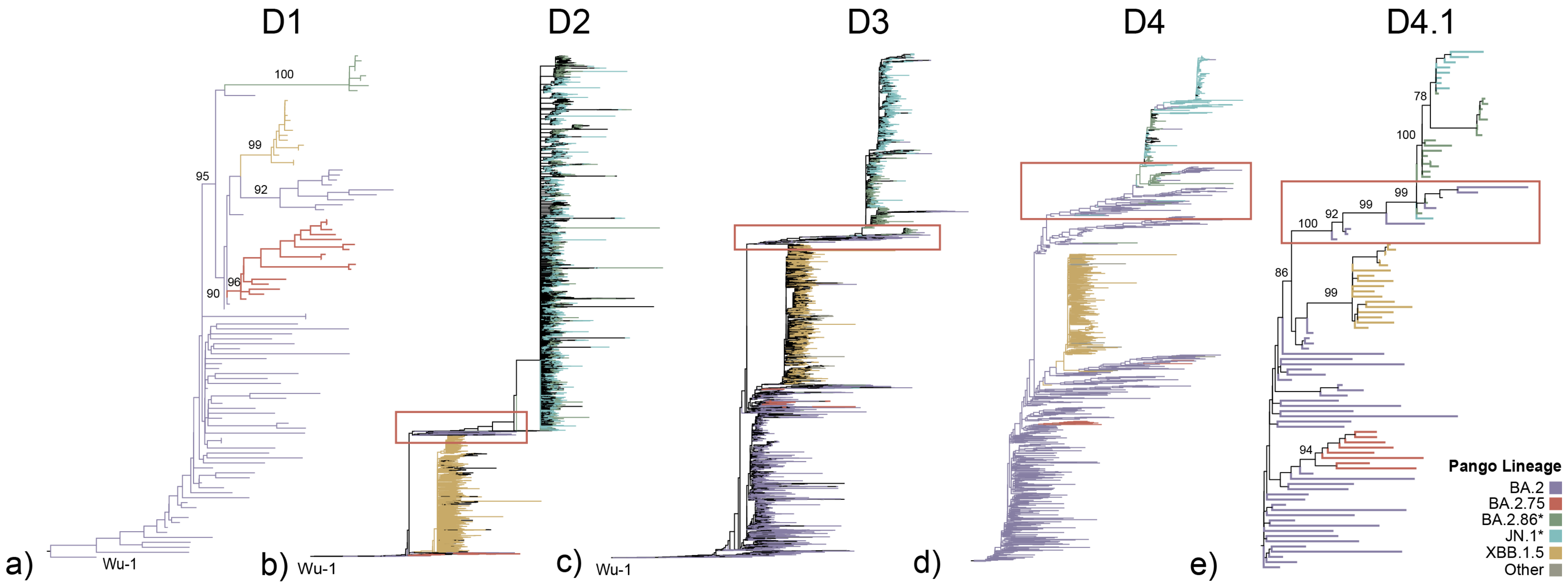


Maximum-likelihood (ML) trees derived from the D1-D4.1 datasets with branches coloured according to Pango lineage. All trees show a consistent phylogenetic placement for all virus lineages of interest, with a distal positioning of lineages BA.2 (in purple), BA.2.75 (in red), and XBB.1.5 (in yellow), and with the BA.2.86* clade (in green/teal) diverging from the parental BA.2. In the D1 tree, consistent with the initial characterization of this VOI [2], the BA.2.86 is separated by a long branch. However, within the D2-D4 trees, we observe a cluster of approximately 100 BA.2 genomes positioned as directly ancestral and effectively shortening the branch leading to the BA.2.86* clade (highlighted with red boxes). With D1 and D4.1 tree as an illustrative example, bootstrap values are indicated for branches of interest. The D1-D3 trees were rooted using Wu-1, while D4 and D4.1 were rooted to maintain polarity according to D1.

**Supplementary Figure 2. Quality scores and SAV counts for BA.2 assemblies fall within the typical range of other sequencing libraries**

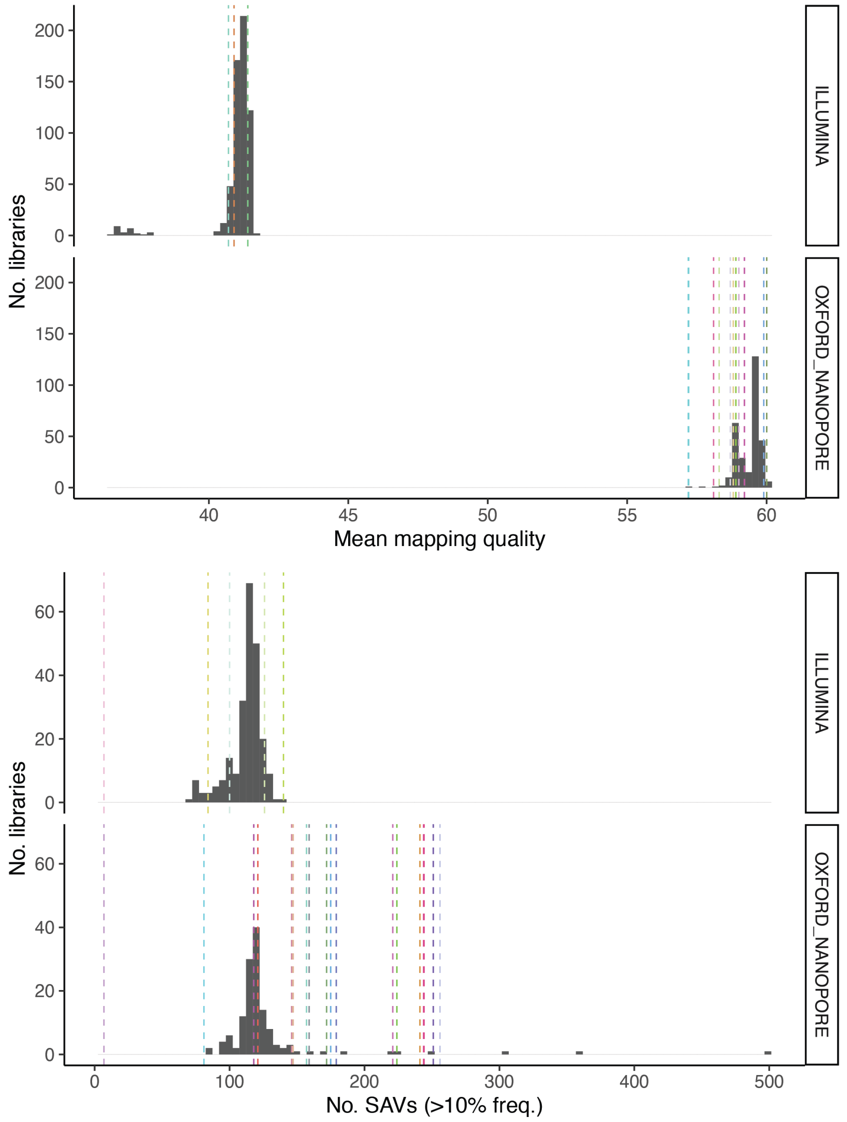


Summary plots showing the distribution of mean quality score metrics for mapping quality and for the number of single amino acid variants (SAVs) with frequency >10%. Grey areas represent background distributions derived from 901 publicly accessible BA.2.86 sequencing libraries, comprising 599 generated through Illumina and 302 generated through Nanopore sequencing technologies. Coloured dashed lines indicate the corresponding mean values for each of the 23 assemblies analysed belonging to the BA.2 cluster directly ancestral to the BA.2.86* clade.

**Supplementary Figure 3. The mutational trajectory of LDMs showing gradual evolution in the BA.2 cluster directly ancestral to BA.2.86**

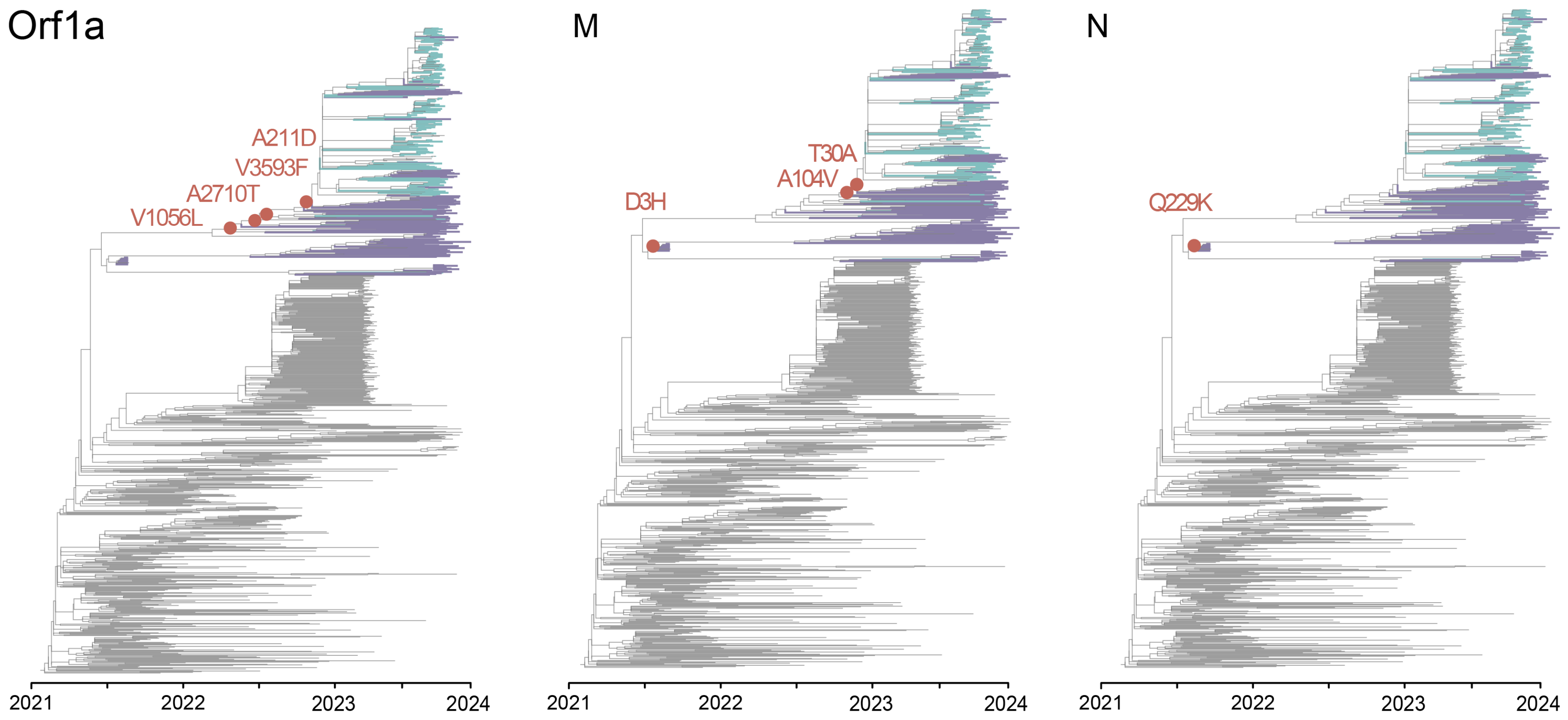


The Orf1a, M, and N full mutational trajectories mapped onto nodes of the time-scaled ML tree derived from our “time-scaled dataset”. Branches are coloured according to Pango lineage, highlighting only the BA.2 cluster enoting evolutionary intermediates (in purple) and the BA.2.86* clade (in teal). A proportion of LDMs (4 in Orf1a, 3 in M, and 1 in N) locate onto deeper nodes of the tree (shown in red circles), where it is evident that some LDMs were sequentially fixed in the BA.2 cluster directly ancestral to the BA.2.86* clade.

**Supplementary Figure 4. Occurrence of LDMs across non-BA.2-86* genomes**

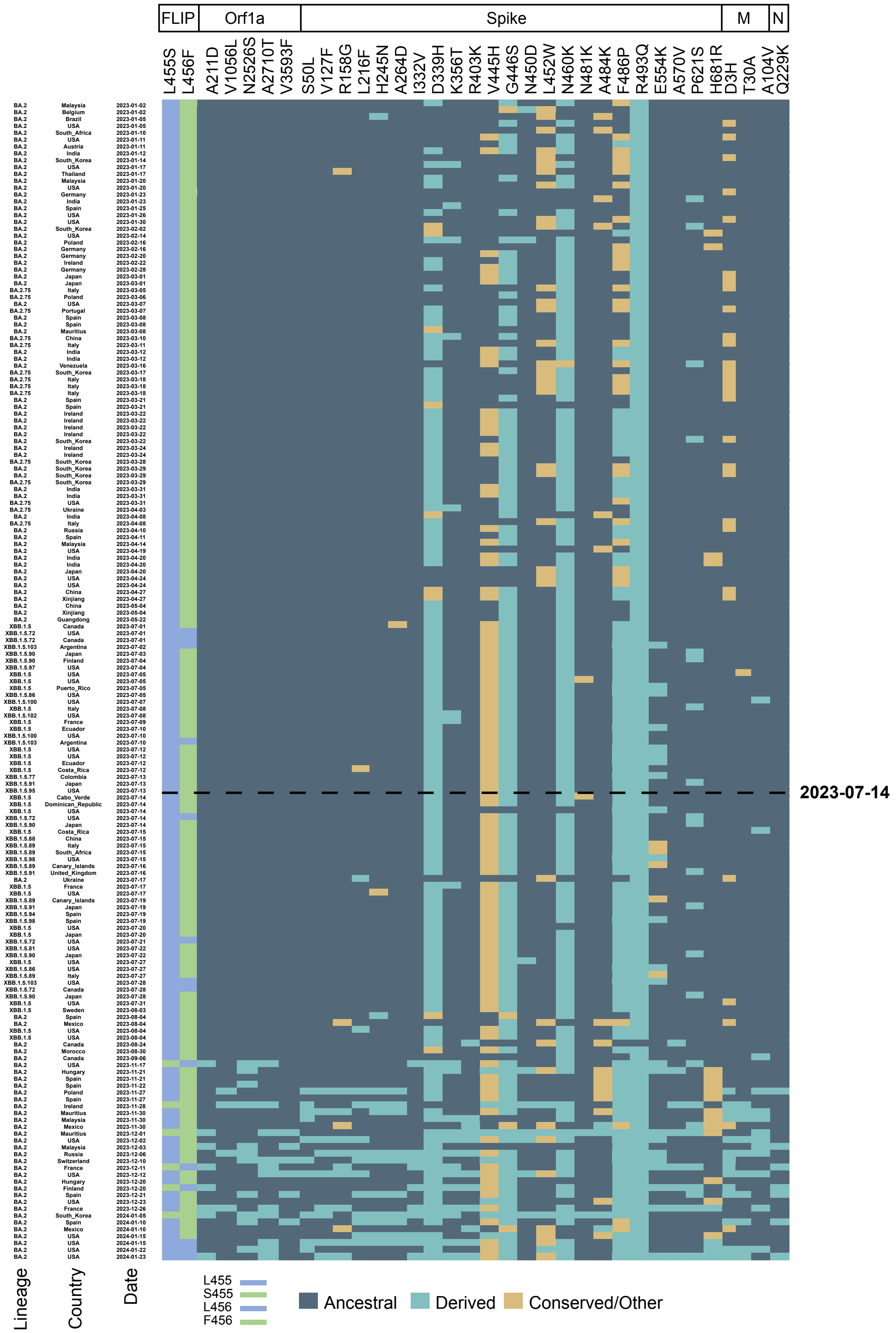


Heatmap for a subset of 170 BA.2, BA.2.75 and XBB.1.5-assigned genomes homogeneously subsampled according to collection date (from January 2023 to January 20249, displaying a variable LDM presence/absence pattern. “BA.2.86-specific”-LMDs are shown as 'derived' states in teal, while 'ancestral' states (*i.e.,* those present in other closely related BA.2* lineages but not in BA.2.86*) are shown in dark blue. Conserved or other states are shown in ochre. The dashed line marks the detection date of BA.2.86

**Supplementary Figure 5. Normalized mutation counts against ambiguous content per genome**

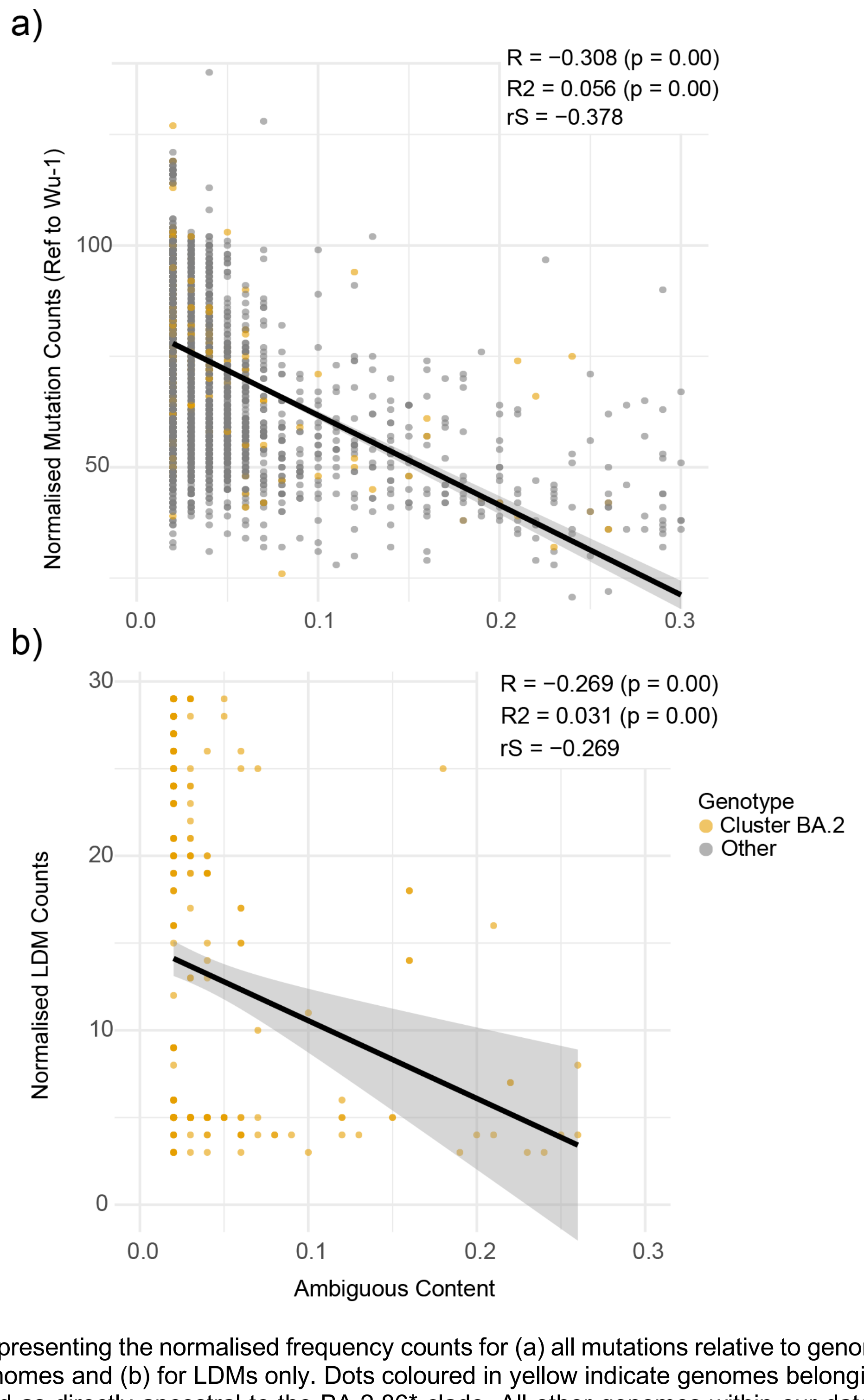


Scatter plots representing the normalised frequency counts for (a) all mutations relative to genome ambiguous content per genomes and (b) for LDMs only. Dots coloured in yellow indicate genomes belonging to the BA.2 cluster identified as directly ancestral to the BA.2.86* clade. All other genomes within our data are shown in grey. Linear regression (R, R2) and Spearman Correlation Coefficient (rS) tests reveal a negative correlation between variables, indicating that lower sequence quality can be generally associated with a reduction of SAVs, rather than with an increase. Genomes belonging to the BA.2 cluster are interspersed within the data.

**Supplementary Figure 6. Mutational trajectory inferred on P1 and P2 recombination trees**

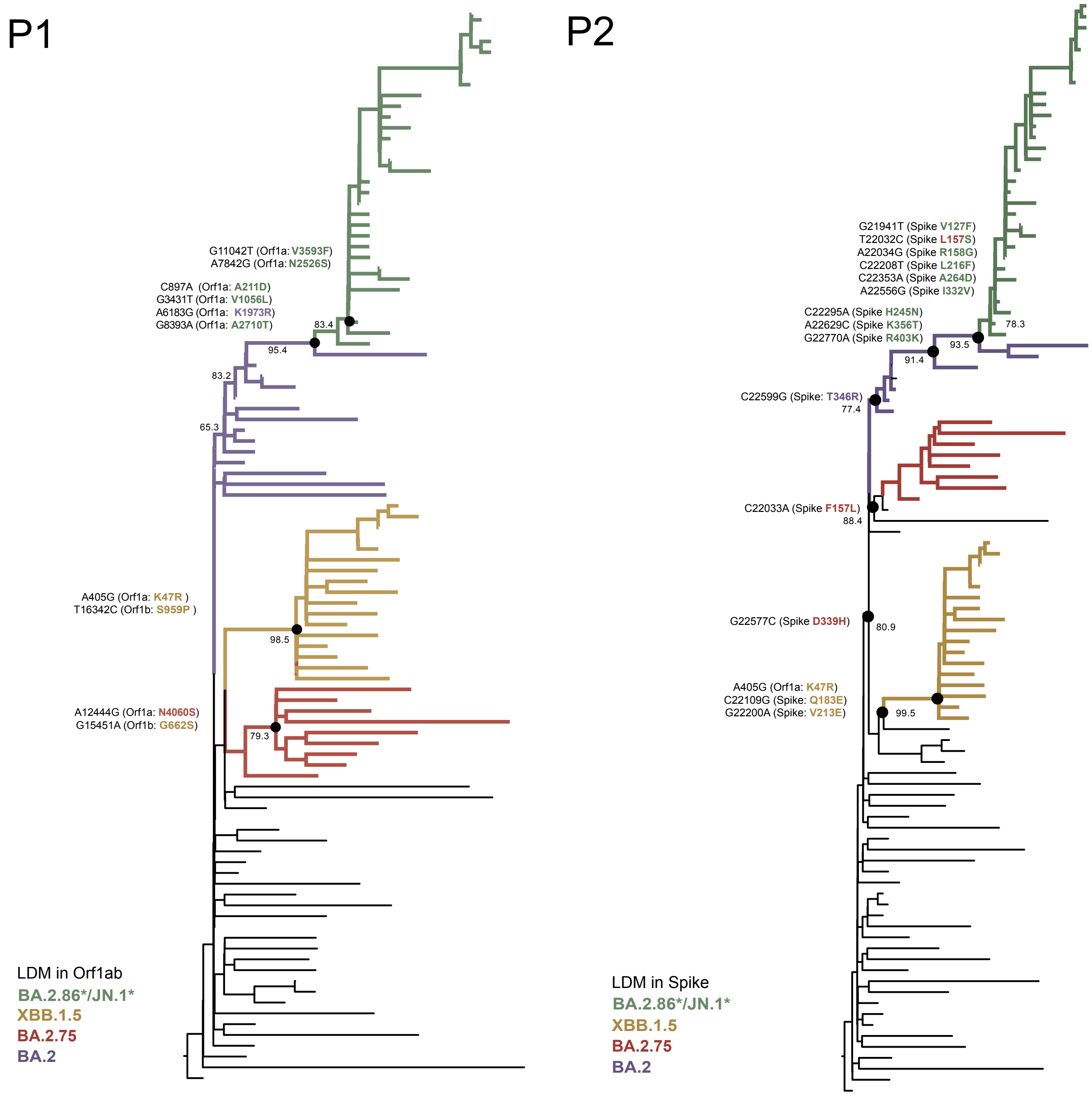


The full mutational trajectories at an amino acid level mapped onto deeper nodes of each of the ML partition trees (P1, P2) derived from the GARD analysis of D4.1. LDMs are coloured according to Pango lineage, highlighting lineage-specific 'private' LDMs for BA.2 (in purple), BA.2.75 (in red), XBB.1.5 (in yellow), and for the BA.2.86* lineages (in green) (see *Supplementary Table 2*, where homoplasic sites/mutations shared across two or more lineages are shown in black). Bootstrap support values are shown for the branches of interest. In the P1 tree, we identify one mutation in Orf1ab (K1973R) as directly contributed through the parental BA.2 lineage. In the P2 tree, mutation F157L in Spike, was directly contributed by BA.2.75 following recombination.

#### Supplementary Table 2. LDMs acquired through recombination

| BA.2.86* | | | XBB | | | BA.2.75 | | | BA.2 (distal) | |
|---|---|---|---|---|---|---|---|---|---|---|
| ORF1ab (5) | S (23) | | ORF1ab (15) | S (38) | | ORF1ab (16) | S (30) | | ORF1ab (13) | S (28) |
| A211D | S50L | | K47R | T19I | | S135R | T19I | | S135R | T19I |
| V1056L | V127F | | S135R | L24S | | T842I | L24S | | T842I | L24S |
| N2526S | L157S | | T842I | V83A | | S1221L | F157L | | G1307S | G142D |
| A2710T | R158G | | G1307S | G142D | | G1307S | I210V | | L3027F | V213G |
| V3593F | L216F | | L3027F | H146Q | | P1640S | V213G | | T3090I | G339D |
| K1973R | H245N | | T3090I | Q183E | | L3027F | G257S | | L3201F | S371F |
| | A264D | | L3201F | V213E | | T3090I | D339H | | T3255I | S373P |
| | I332V | | T3255I | G252V | | L3201F | S371F | | P3395H | S375F |
| | D339H | | P3395H | G339H | | T3255I | S373P | | P314L | T376A |
| | K356T | | P314L | R346T | | P3395H | S375F | | R1315C | D405N |
| | R403K | | G662S | L368I | | N4060S | T376A | | I1566V | R408S |
| | V445H | | S959P | S371F | | P314L | D405N | | T2163I | K417N |
| | G446S/G446S | | R1315C | S373P | | G662S | R408S | | K1973R | N440K |
| | N450D | | I1566V | S375F | | R1315C | K417N | | | S477N |
| | L452W | | T2163I | T376A | | I1566V | N440K | | | T478K |
| | N460K/N460K | | | D405N | | T2163I | G446S | | | E484A |
| | N481K | | | R408S | | | N460K | | | Q493R |
| | A484K | | | K417N | | | S477N | | | Q498R |
| | F486P | | | N440K | | | T478K | | | N501Y |
| | R493Q | | | V445P | | | E484A | | | Y505H |
| | E554K | | | G446S | | | Q498R | | | D614G |
| | A570V | | | N460K | | | N501Y | | | H655Y |
| | P621S | | | S477N | | | Y505H | | | N679K |
| | H681R | | | T478K | | | D614G | | | P681H |
| | | | | E484A | | | H655Y | | | N764K |
| | | | | F486P | | | N679K | | | D796Y |
| | | | | F490S | | | P681H | | | Q954H |
| | | | | Q498R | | | N764K | | | N969K |
| | | | | N501Y | | | D796Y | | | |
| | | | | Y505H | | | Q954H | | | |
| | | | | D614G | | | N969K | | | |
| | | | | H655Y | | | | | | |
| | | | | N679K | | | | | | |
| | | | | P681H | | | | | | |
| | | | | N764K | | | | | | |
| | | | | D796Y | | | | | | |
| | | | | Q954H | | | | | | |
| | | | | N969K | | | | | | |

### As defined in (as of July 2024):
https://outbreak.info/situation-reports?xmin=2023-09-25&xmax=2024-03-25&pango=BA.2.86
https://outbreak.info/situation-reports?xmin=2024-02-22&xmax=2024-08-22&pango=XBB.1.5
https://outbreak.info/situation-reports?xmin=2024-02-22&xmax=2024-08-22&pango=BA.2.75
https://outbreak.info/situation-reports?xmin=2024-02-22&xmax=2024-08-22&pango=BA.2
