## Supplementary Data 2 for "Early evolution of BA.2.86 sheds light on the origins of highly divergent SARS-CoV-2 lineages"

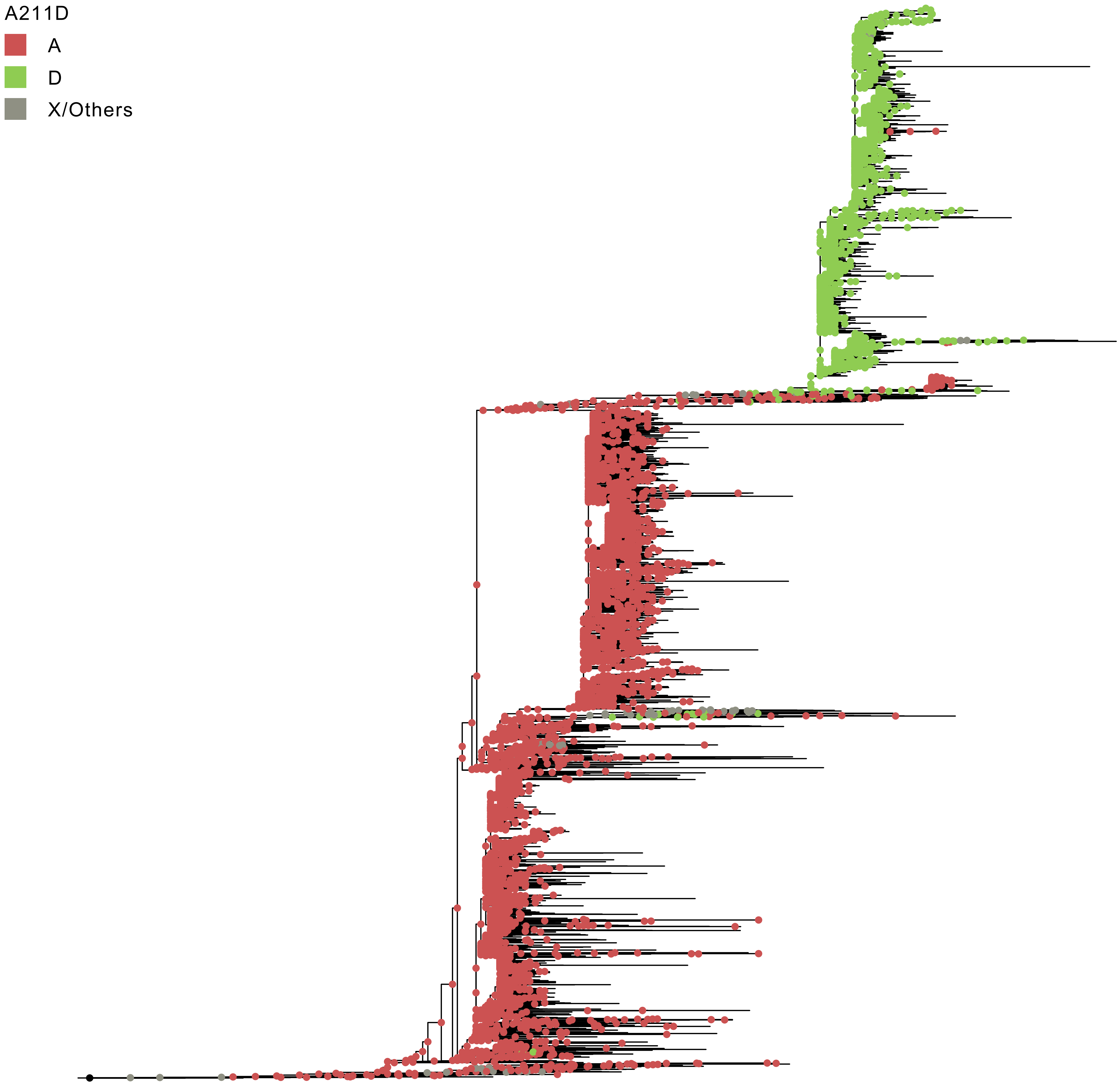
A211D
A
D
X/Others

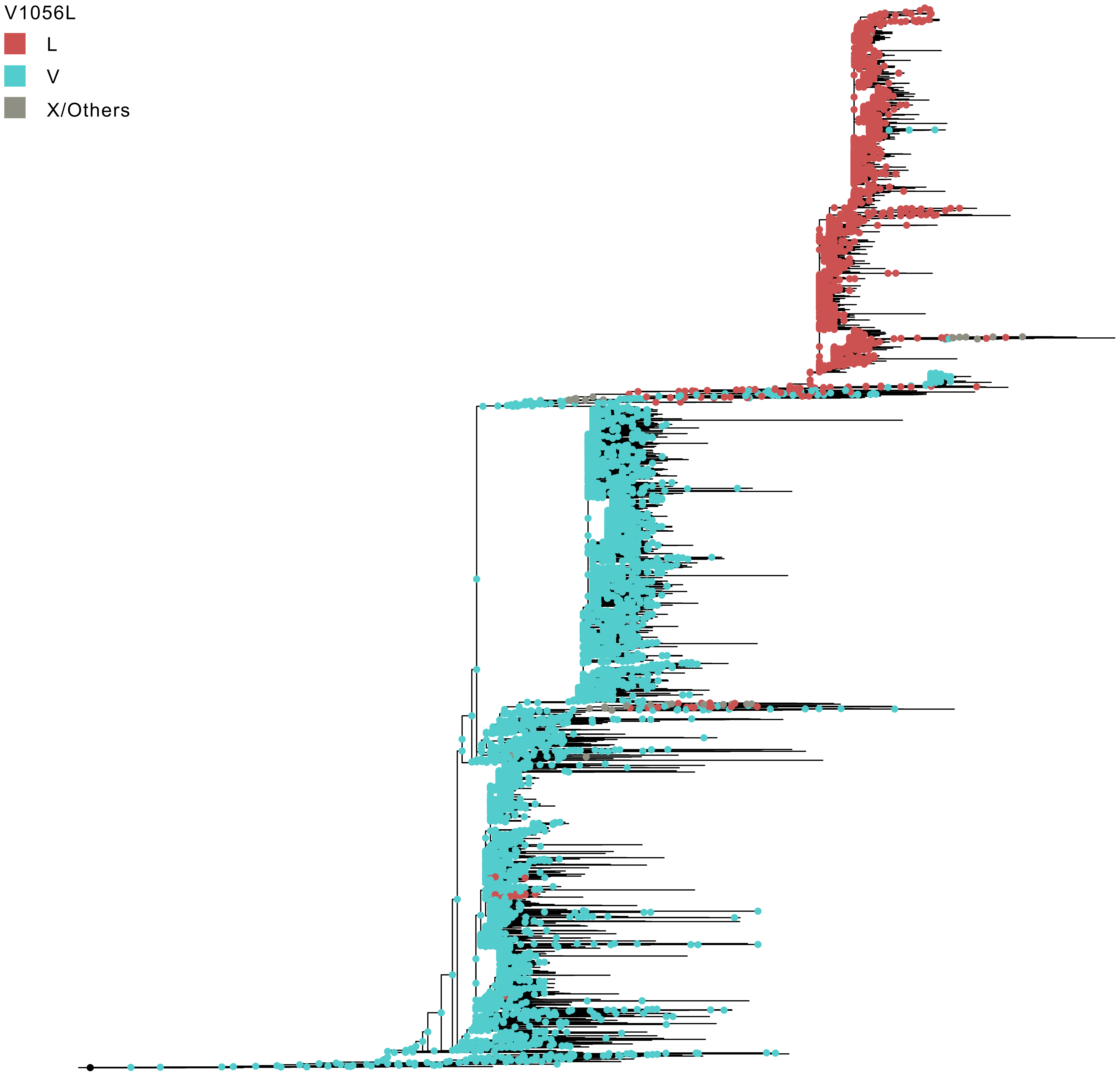
V1056L
L
V
X/Others

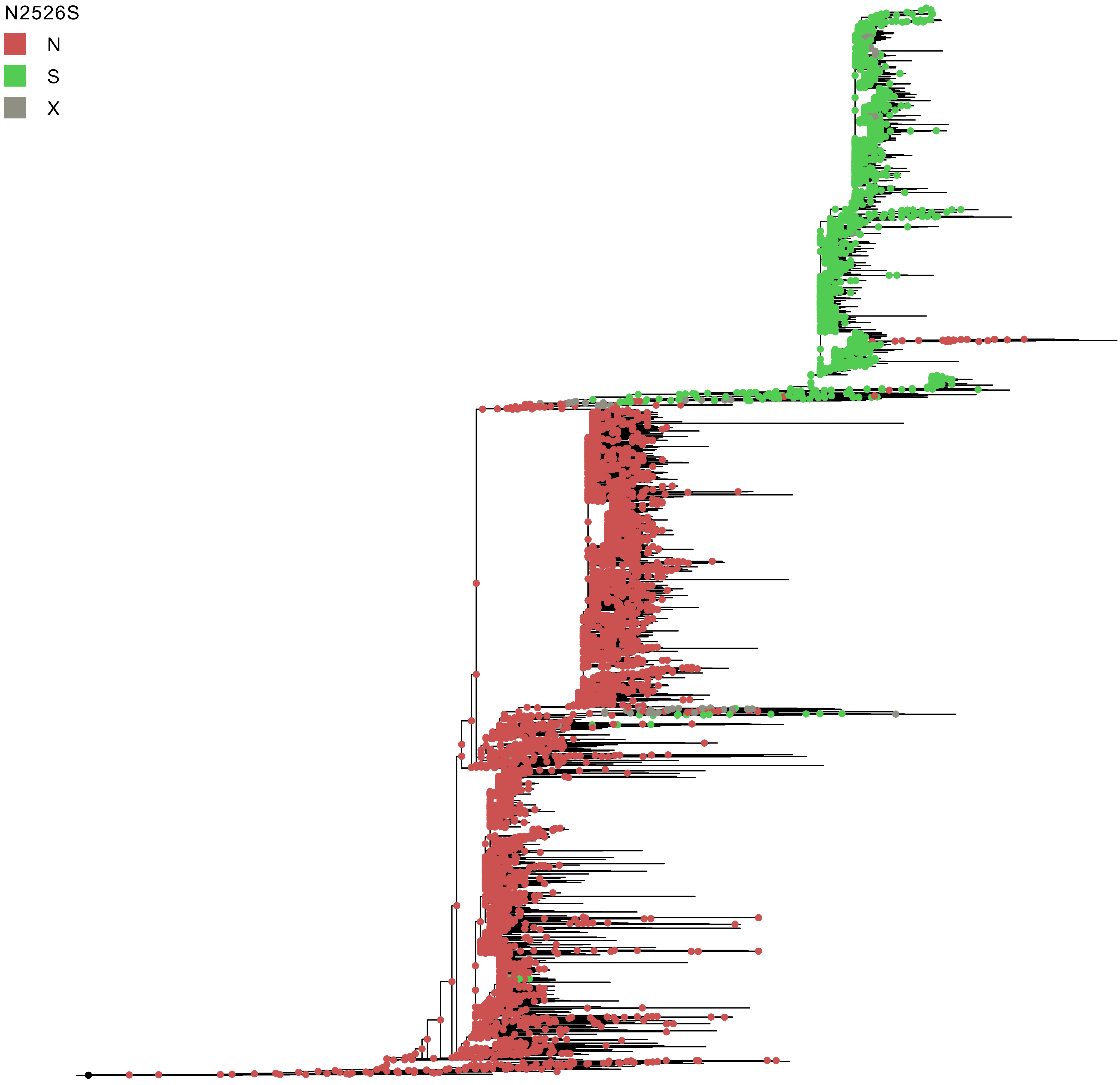
N2526S
N
S
X

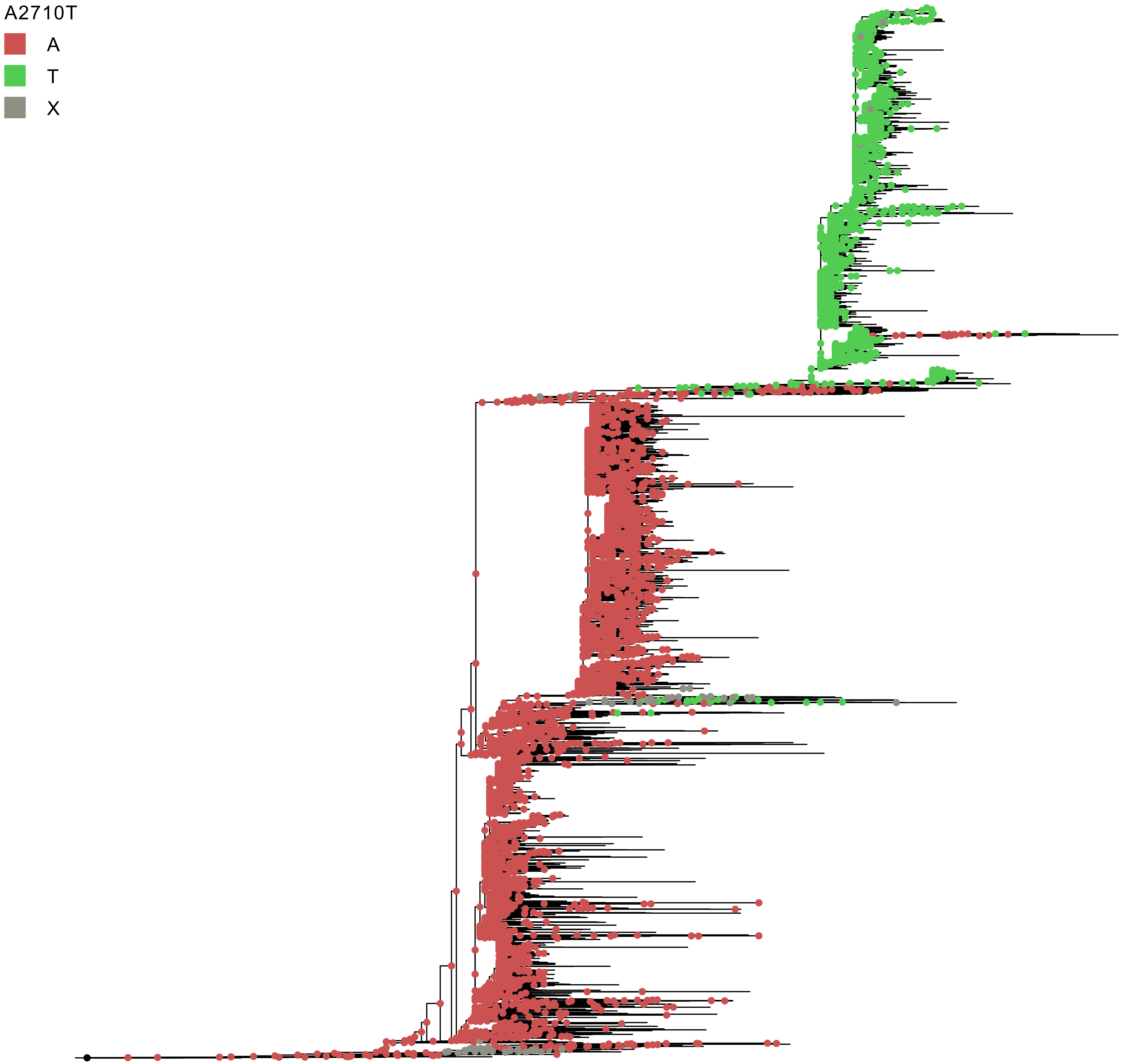
A2710T
A
T
X

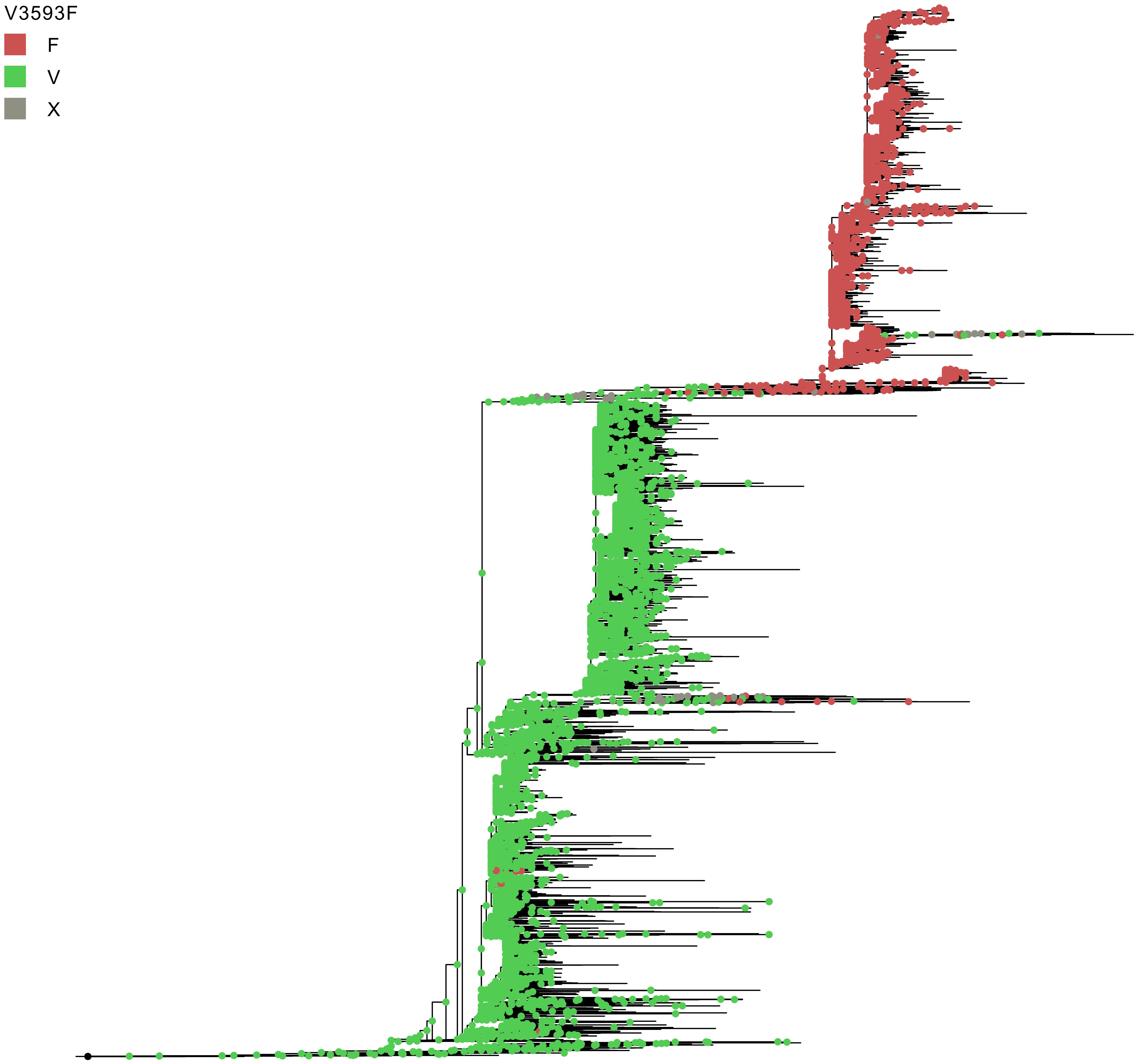
V3593F
F
V
X

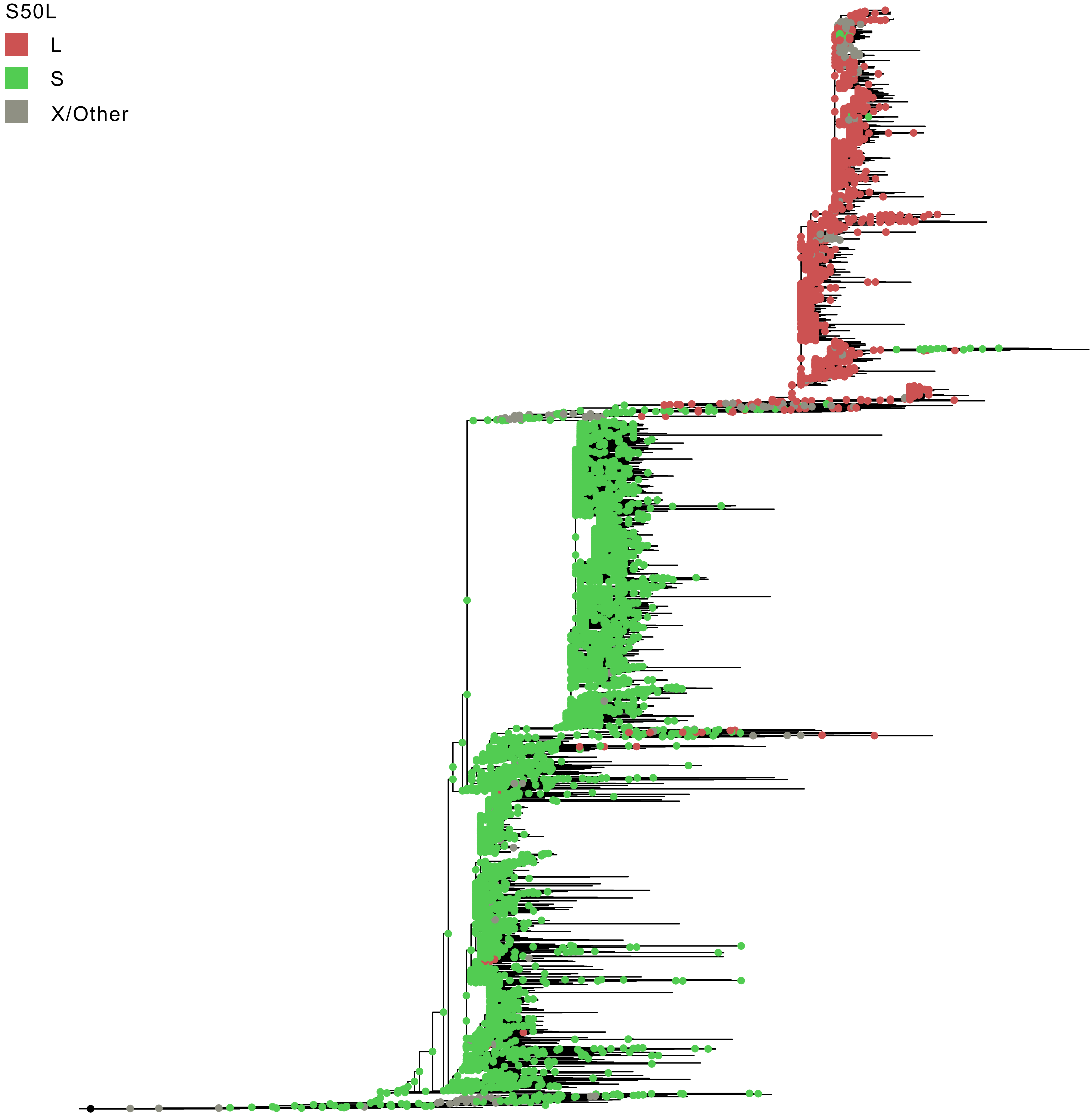
S50L
L
S
X/Other

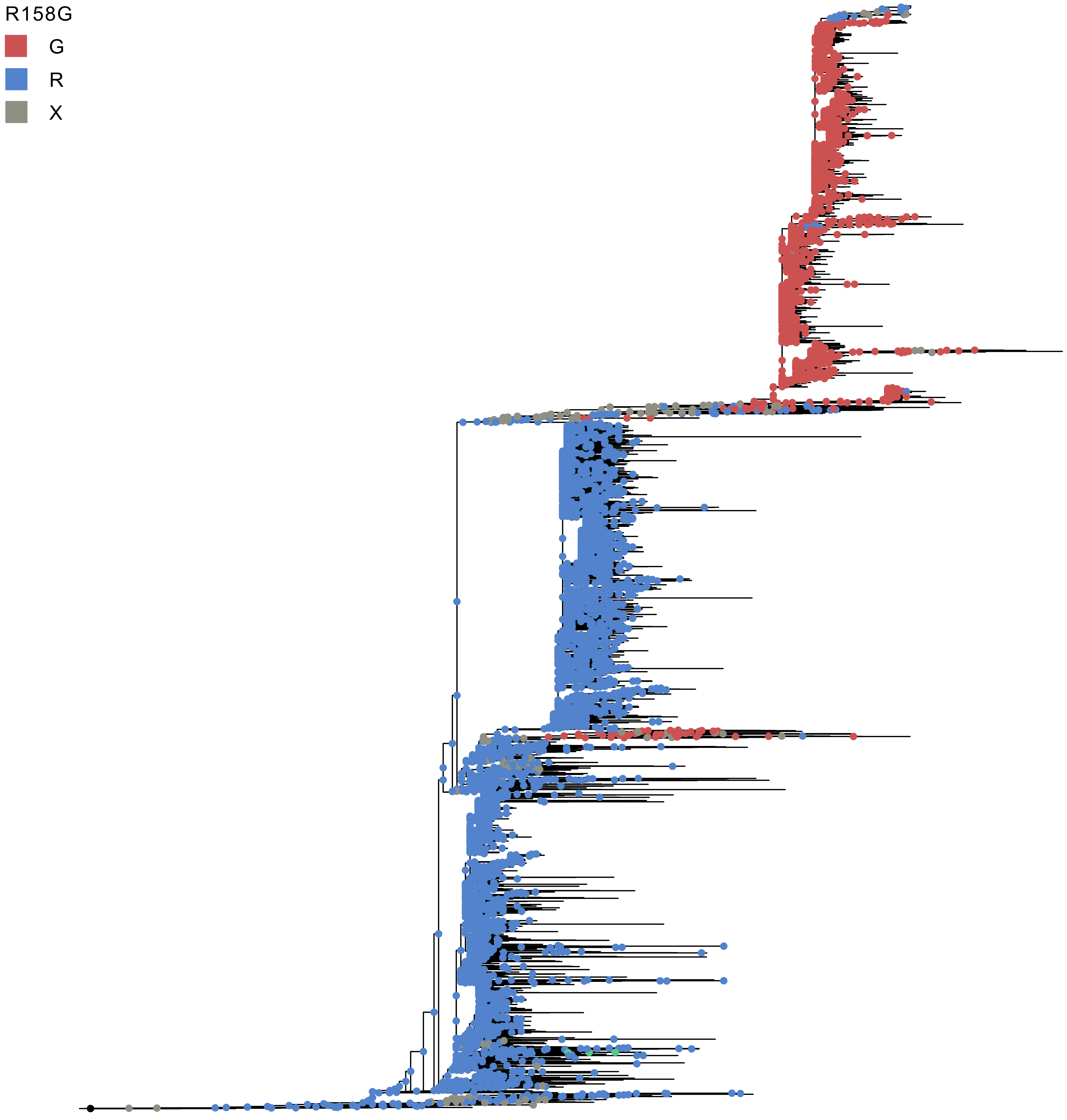
R158G
G
R
X

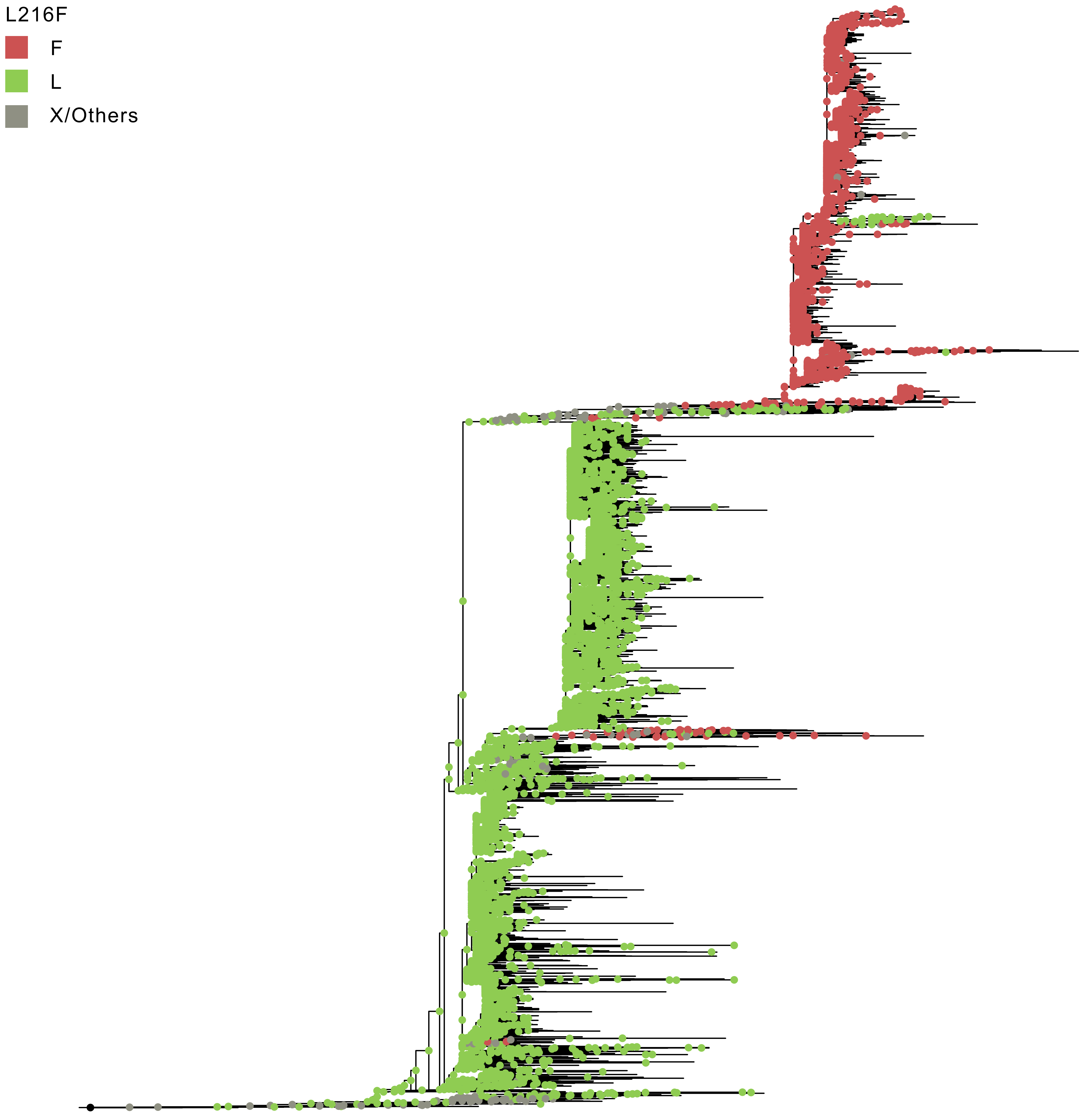
L216F
F
L
X/Others

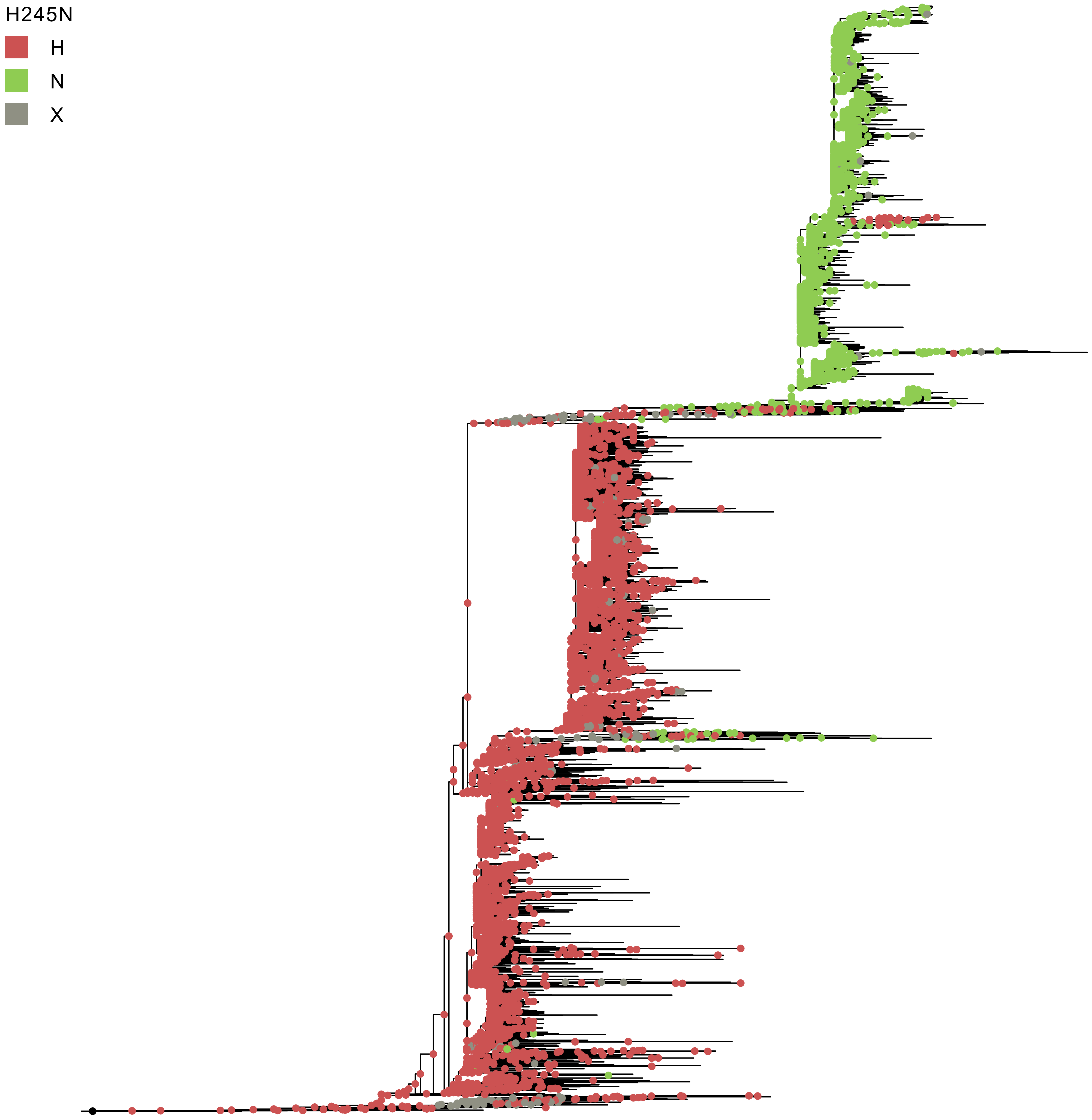
H245N
H
N
X

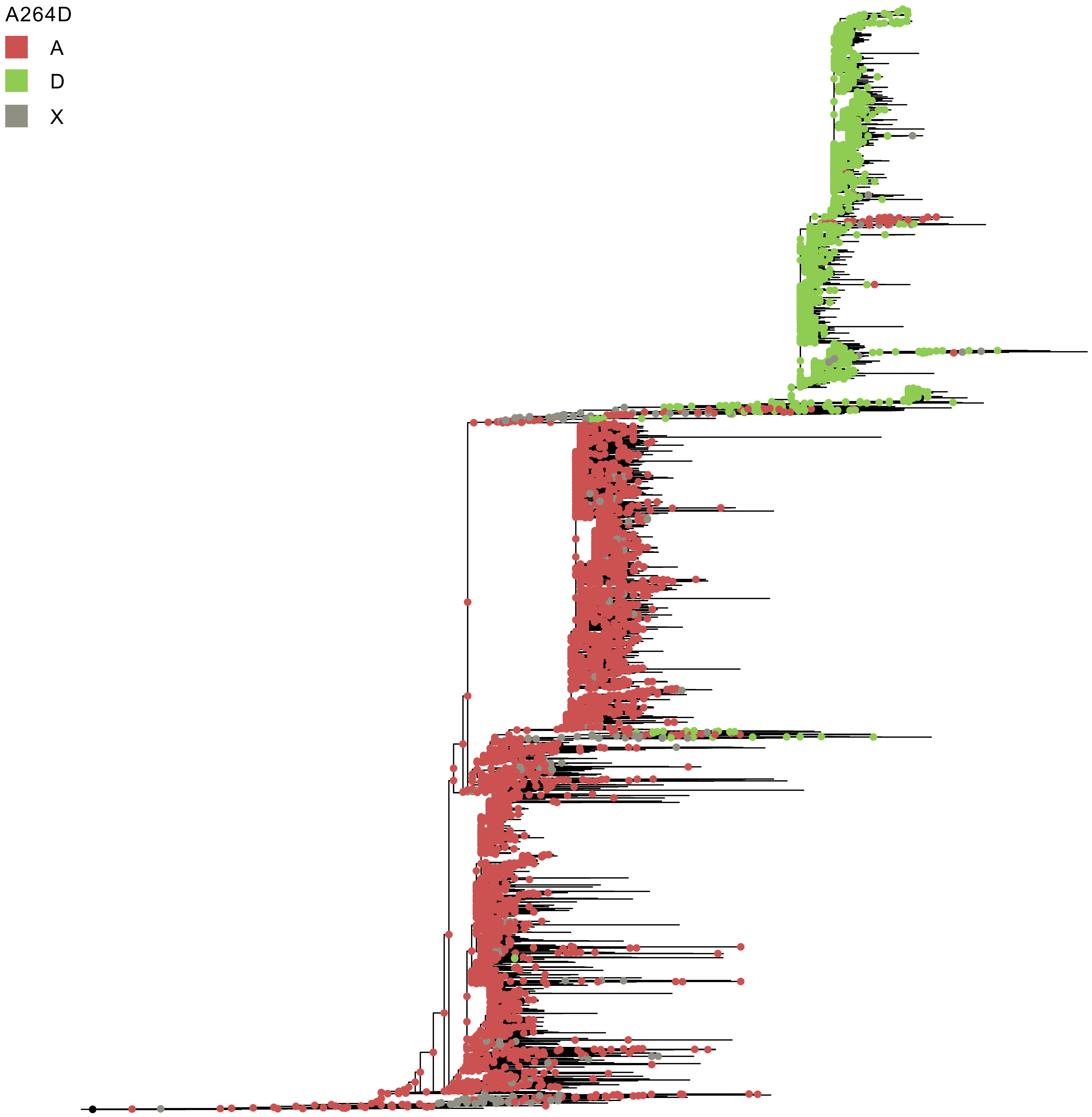

A264D
A
D
X

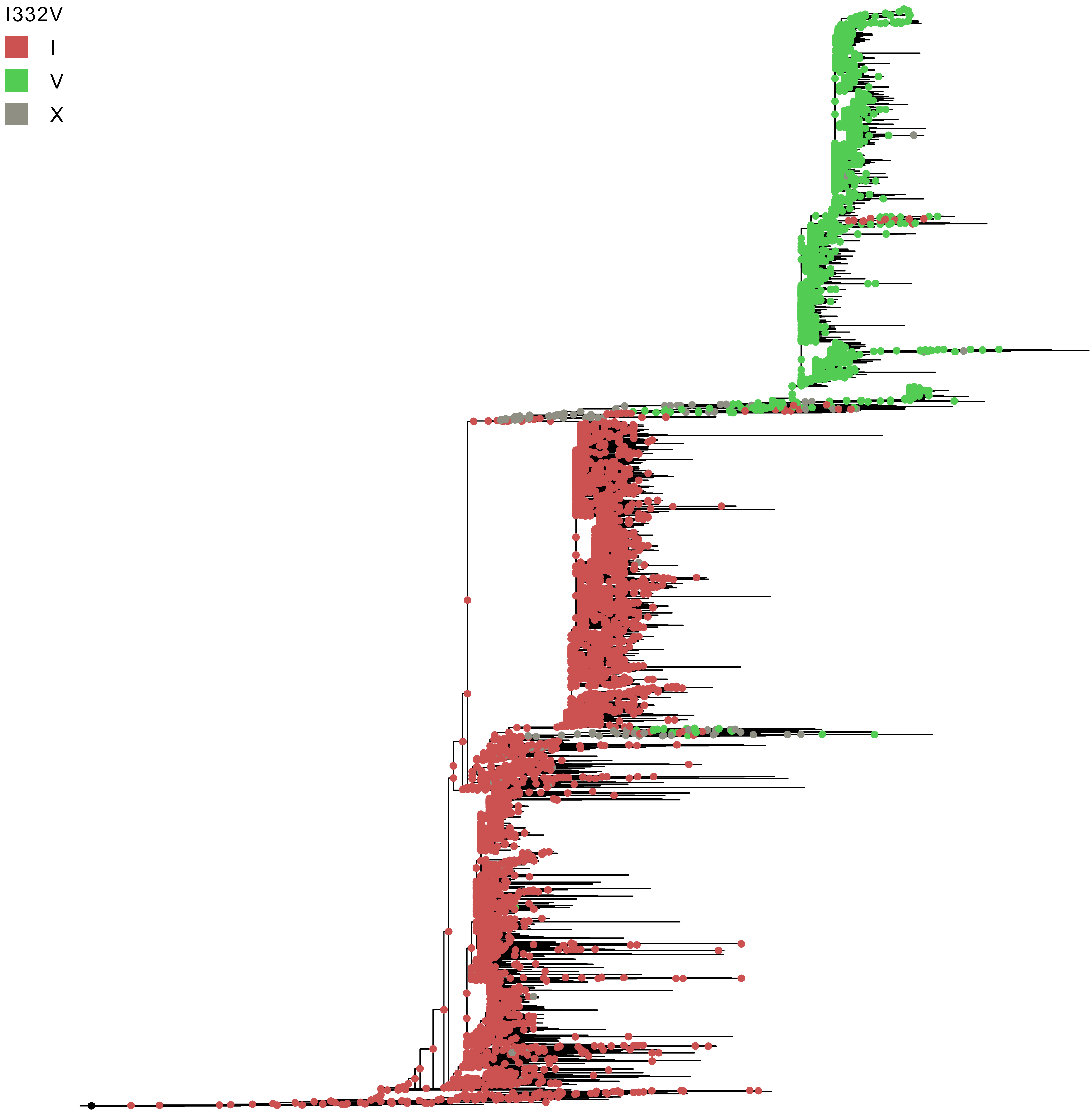

I332V
I
V
X

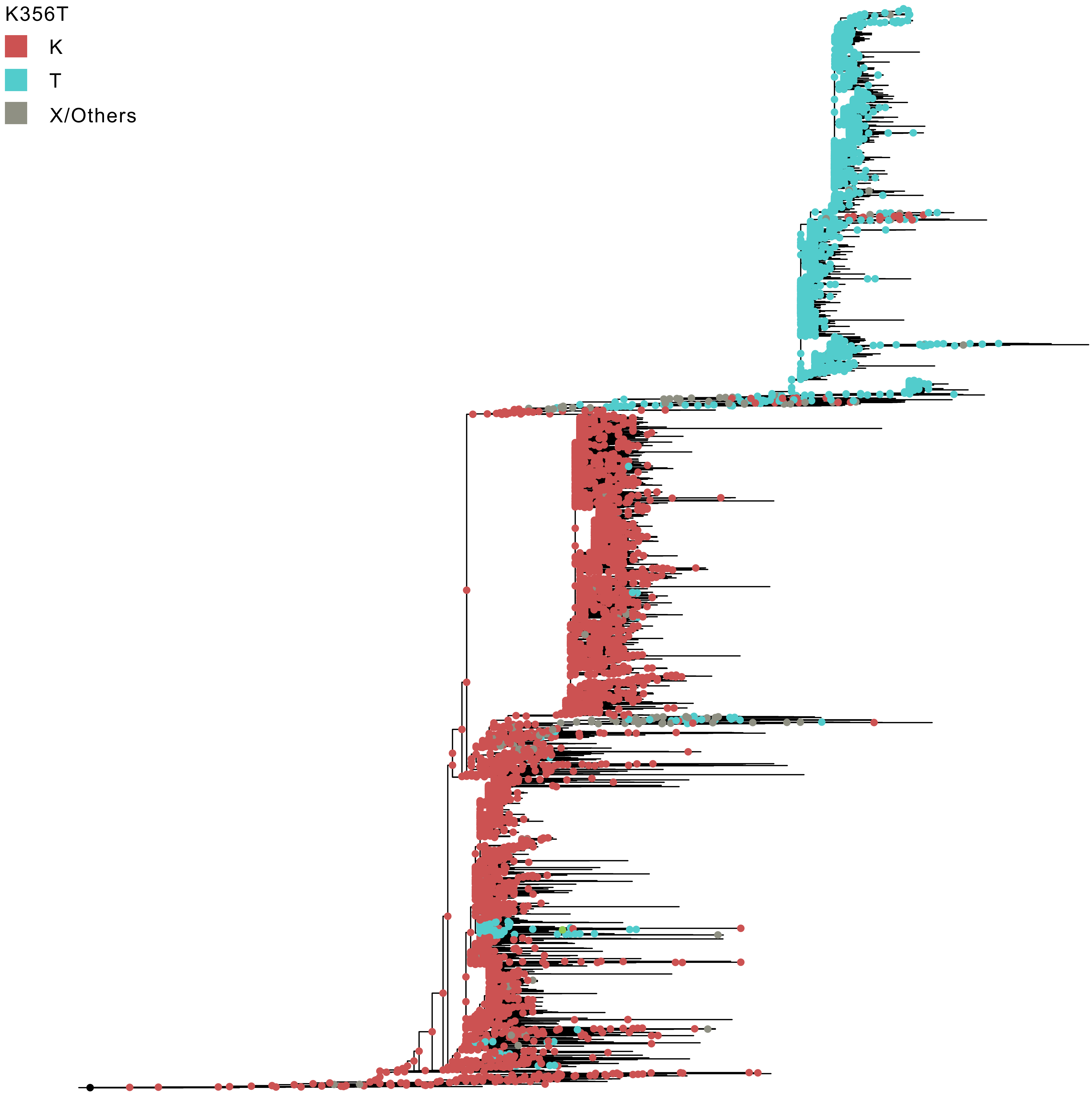
K356T
K
T
X/Others

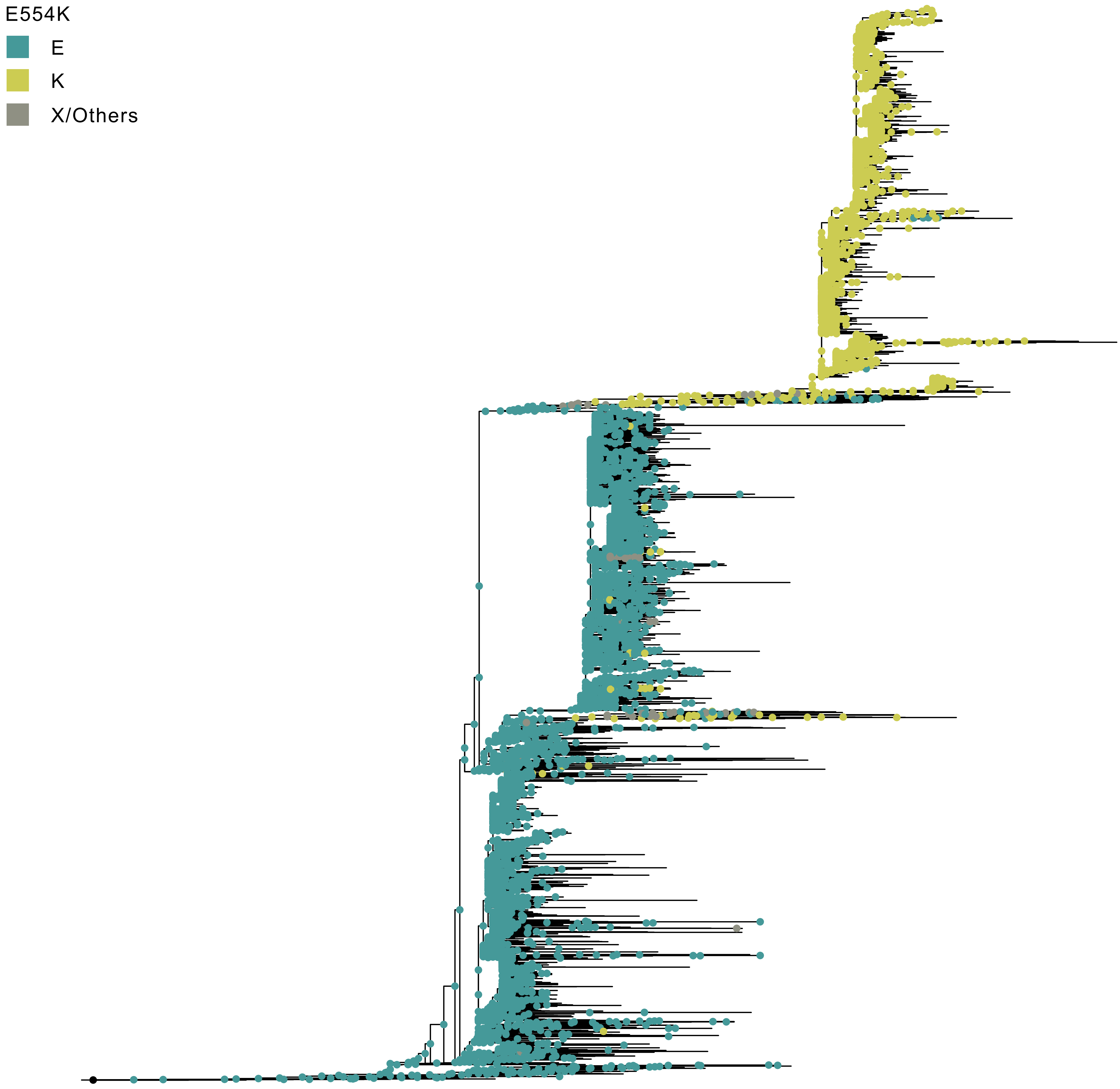
E554K
E
K
X/Others

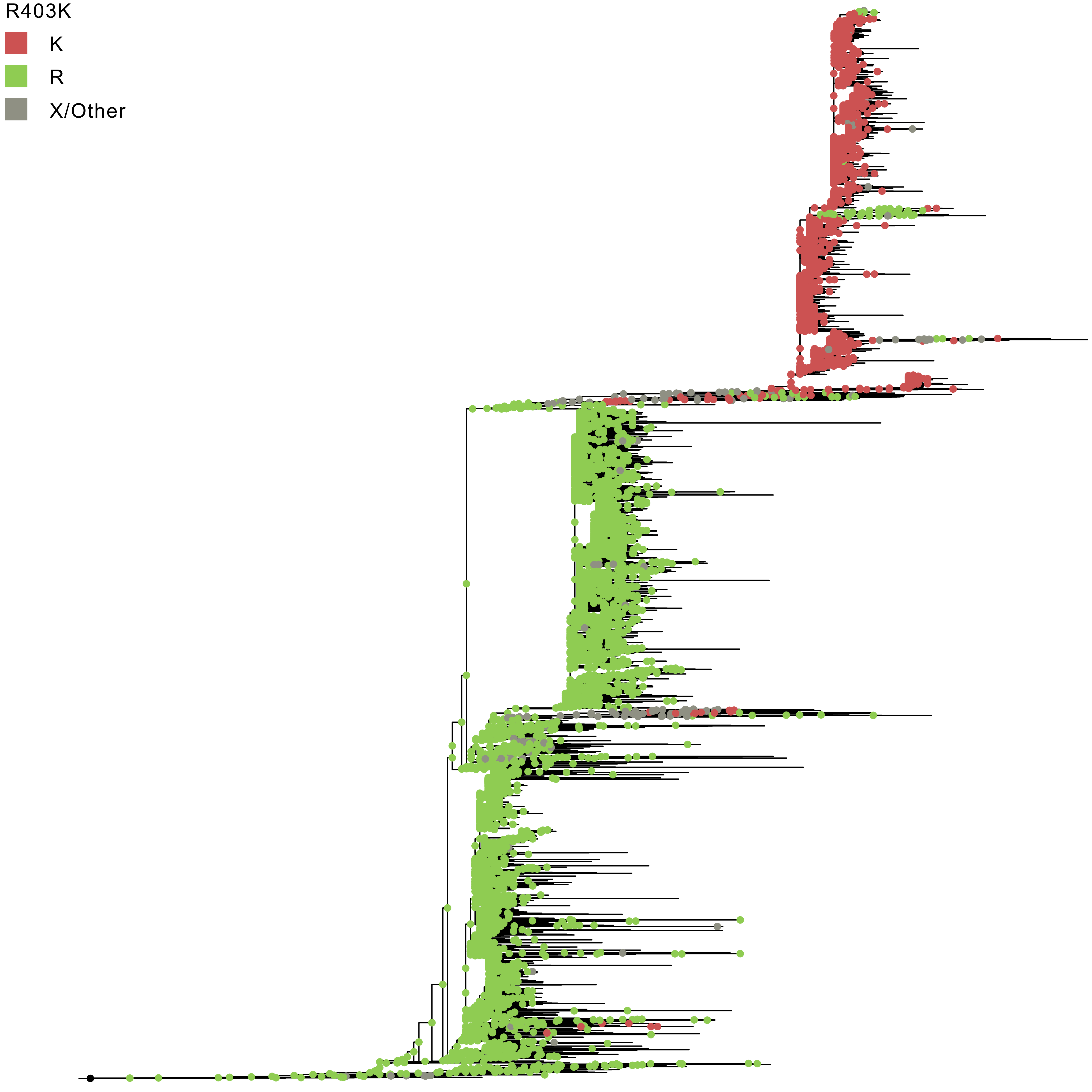
R403K
K
R
X/Other

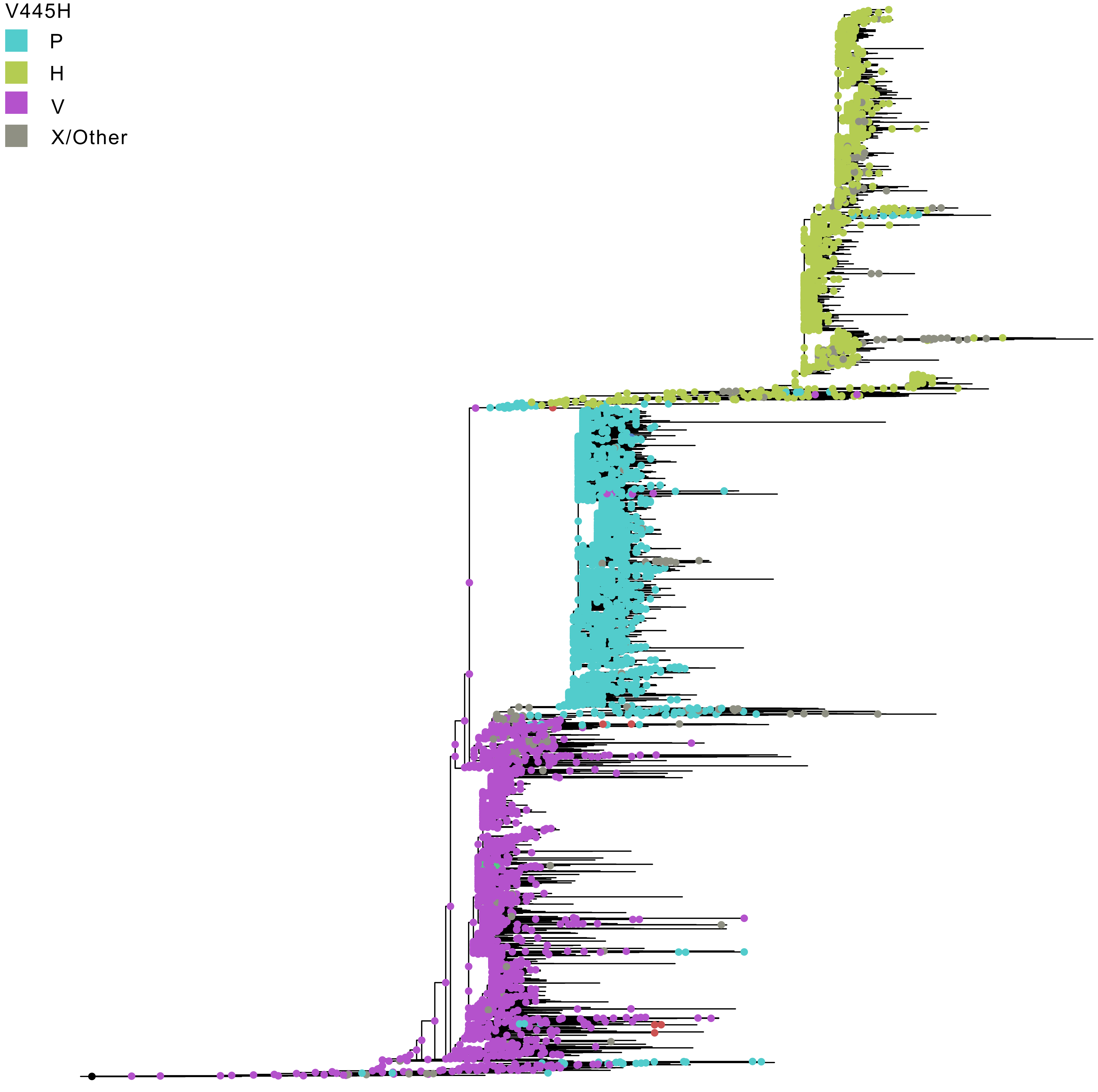
V445H
P
H
V
X/Other

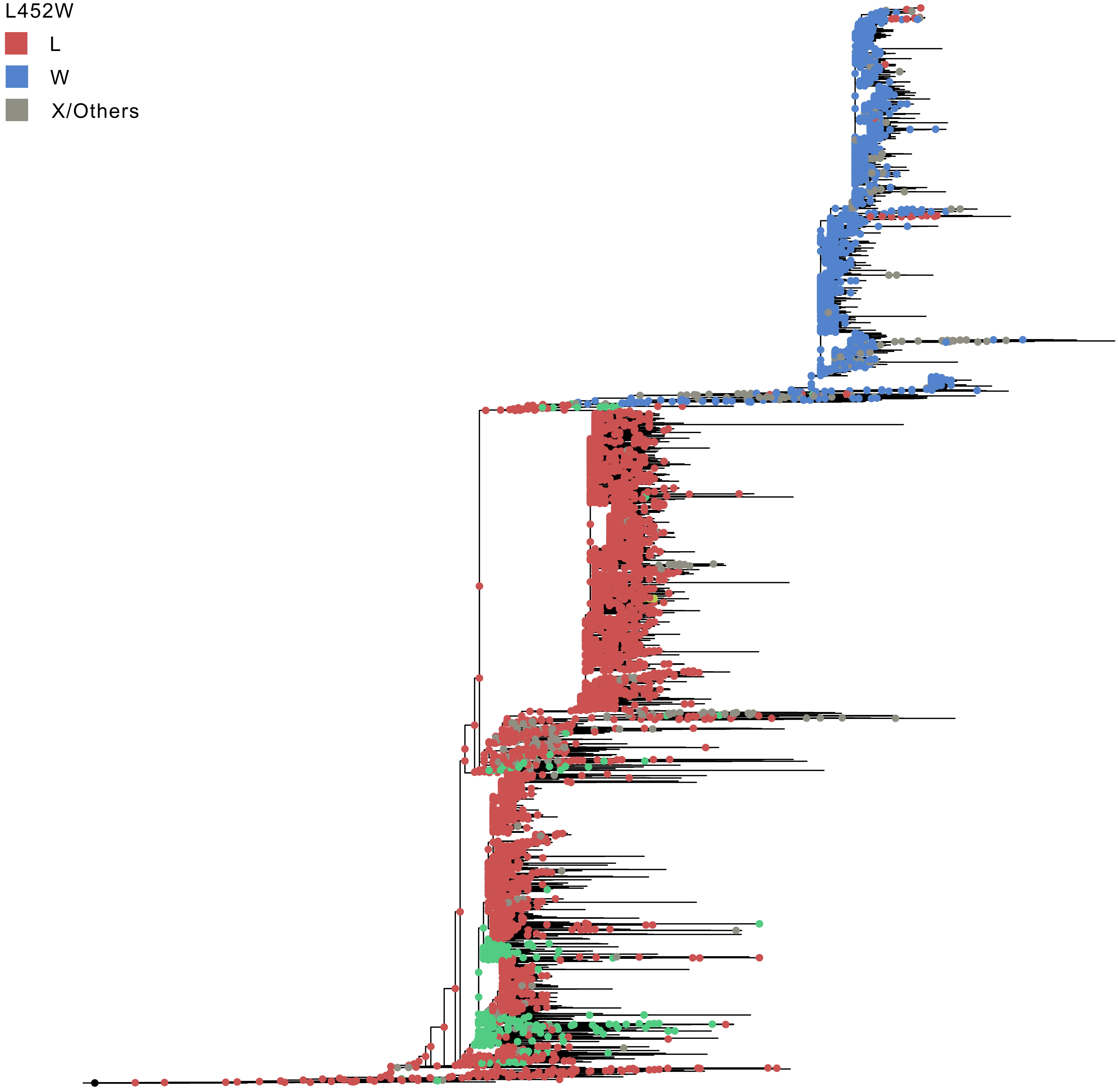
L452W
L
W
X/Others

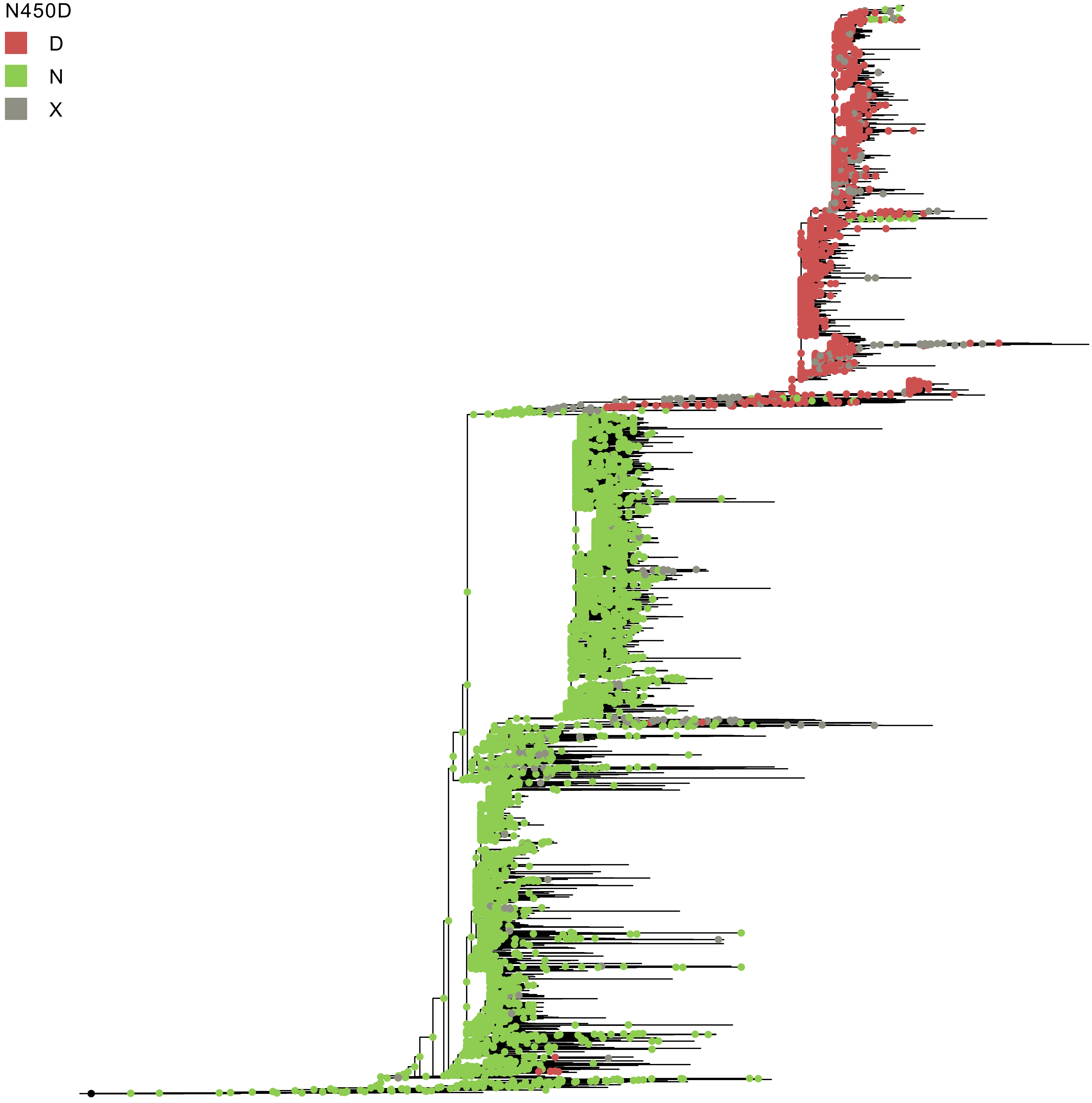
N450D
D
N
X

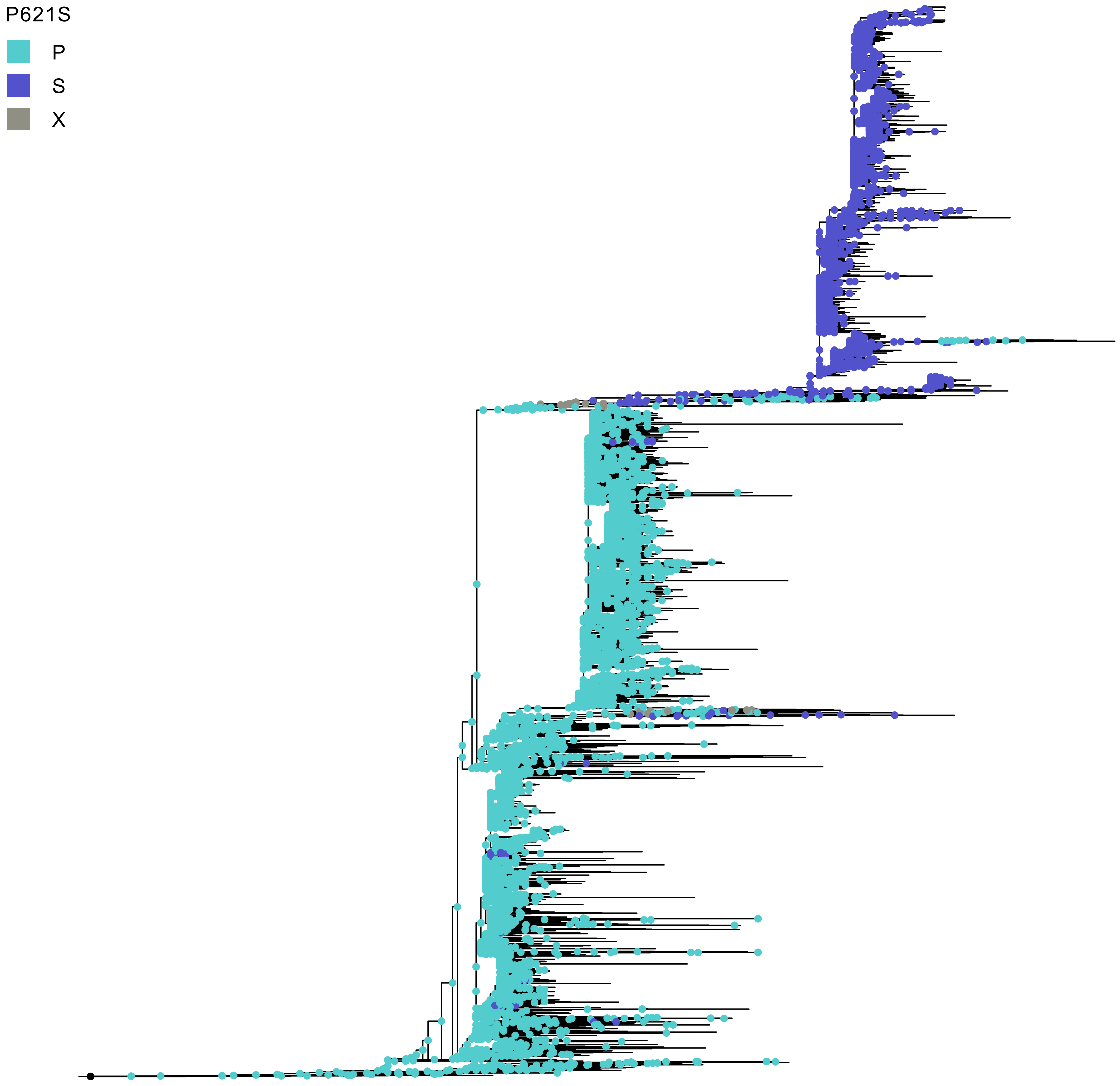
P621S
P
S
X

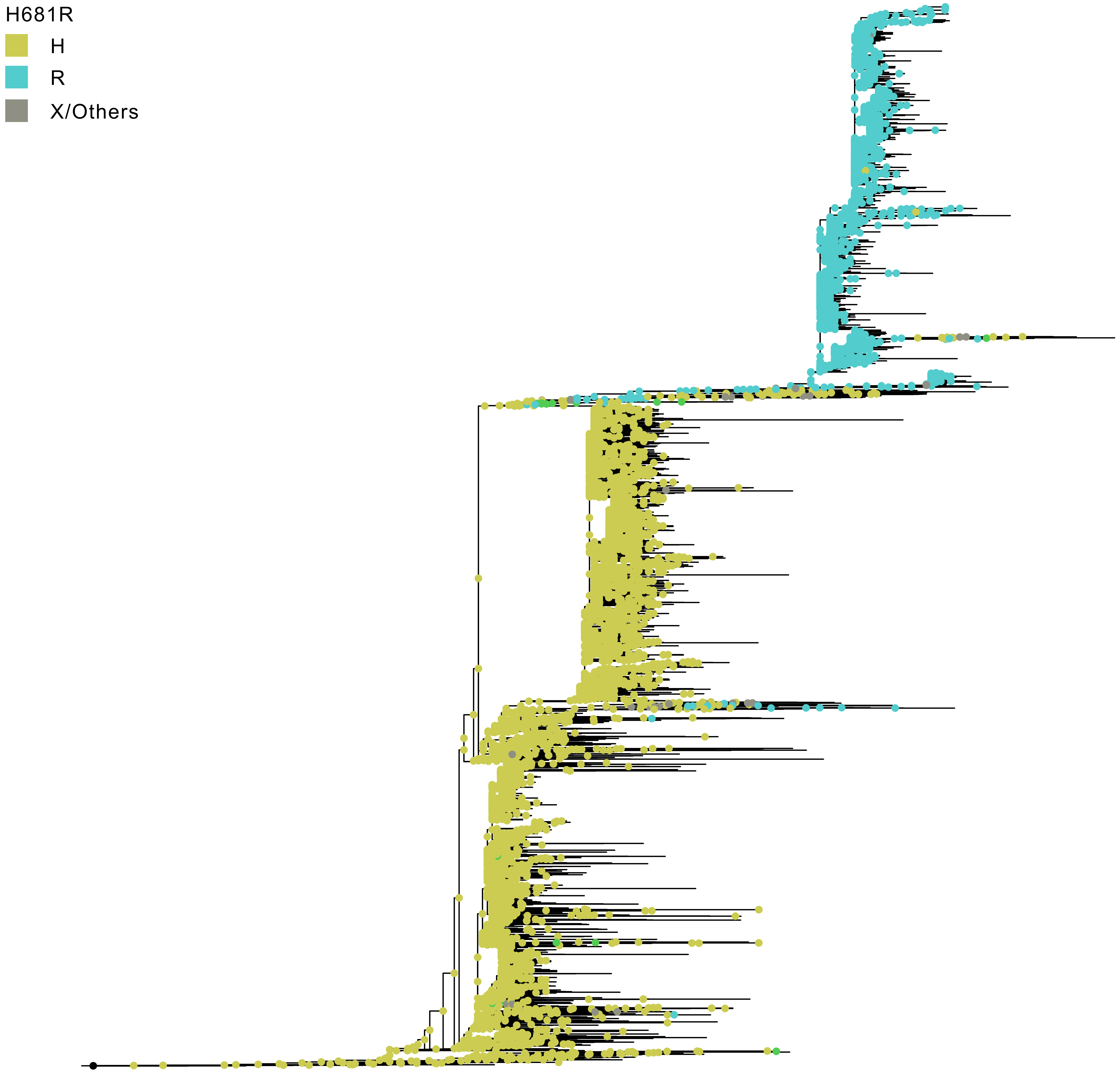
H681R
H
R
X/Others

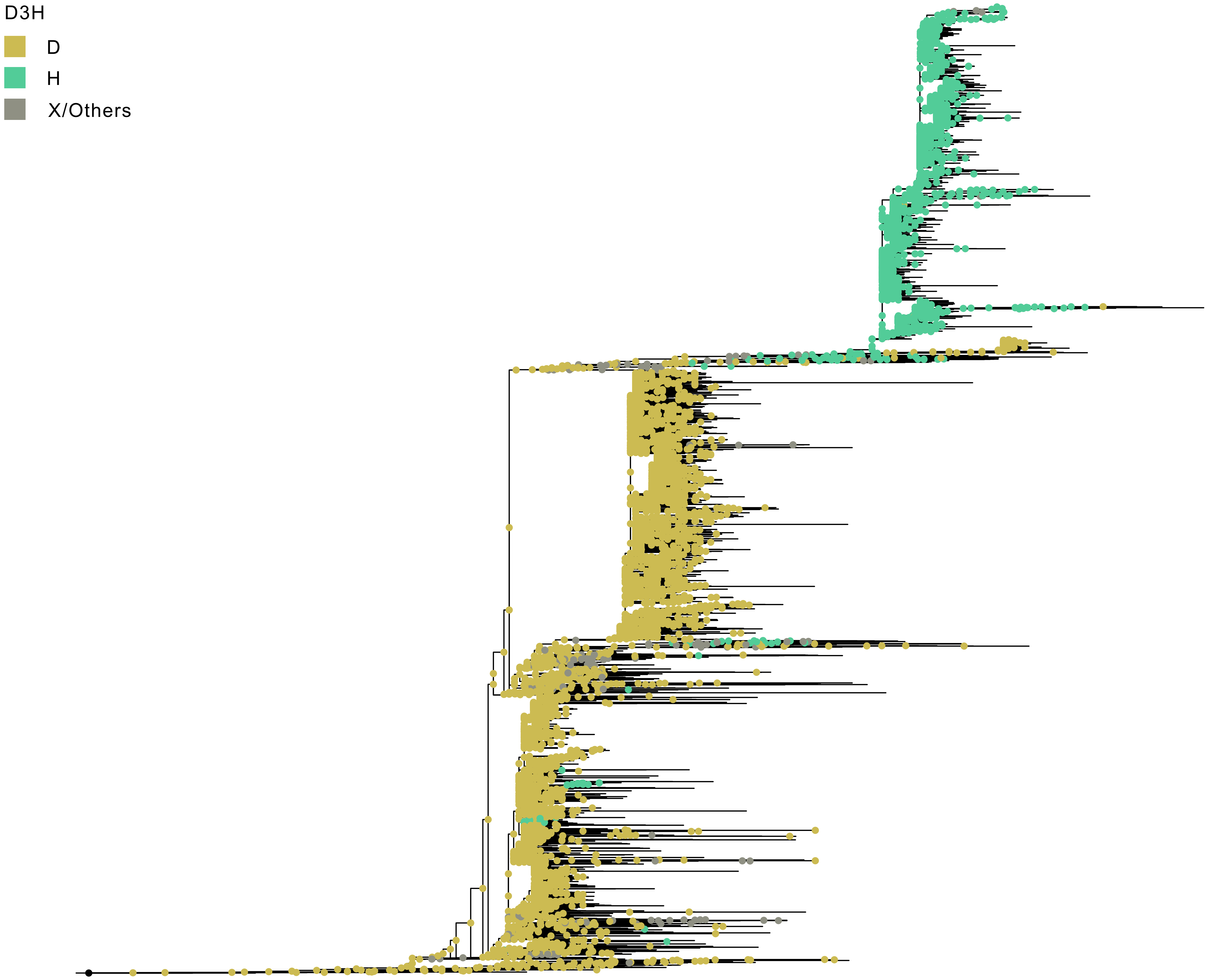
D3H
D
H
X/Others

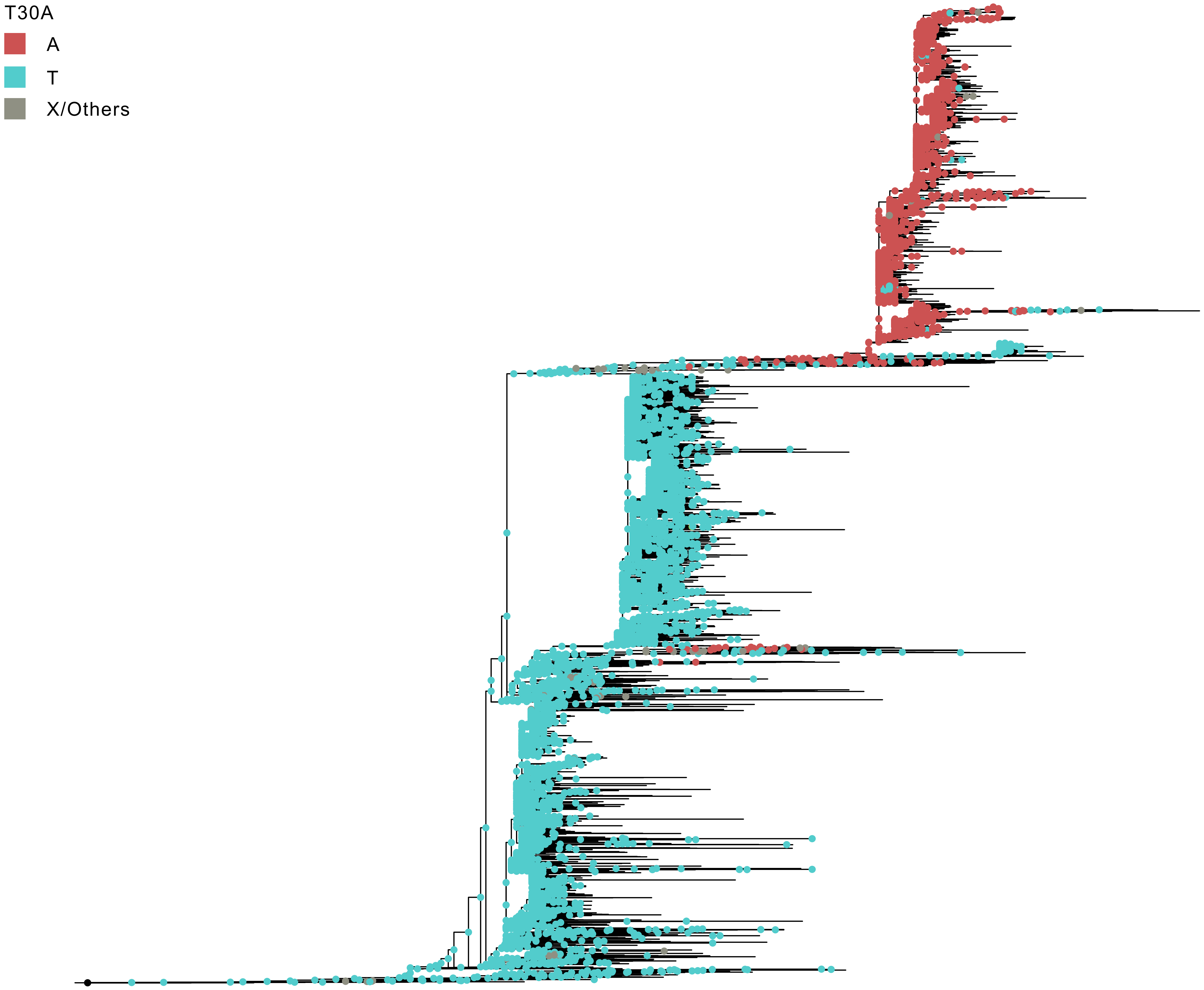
T30A
A
T
X/Others

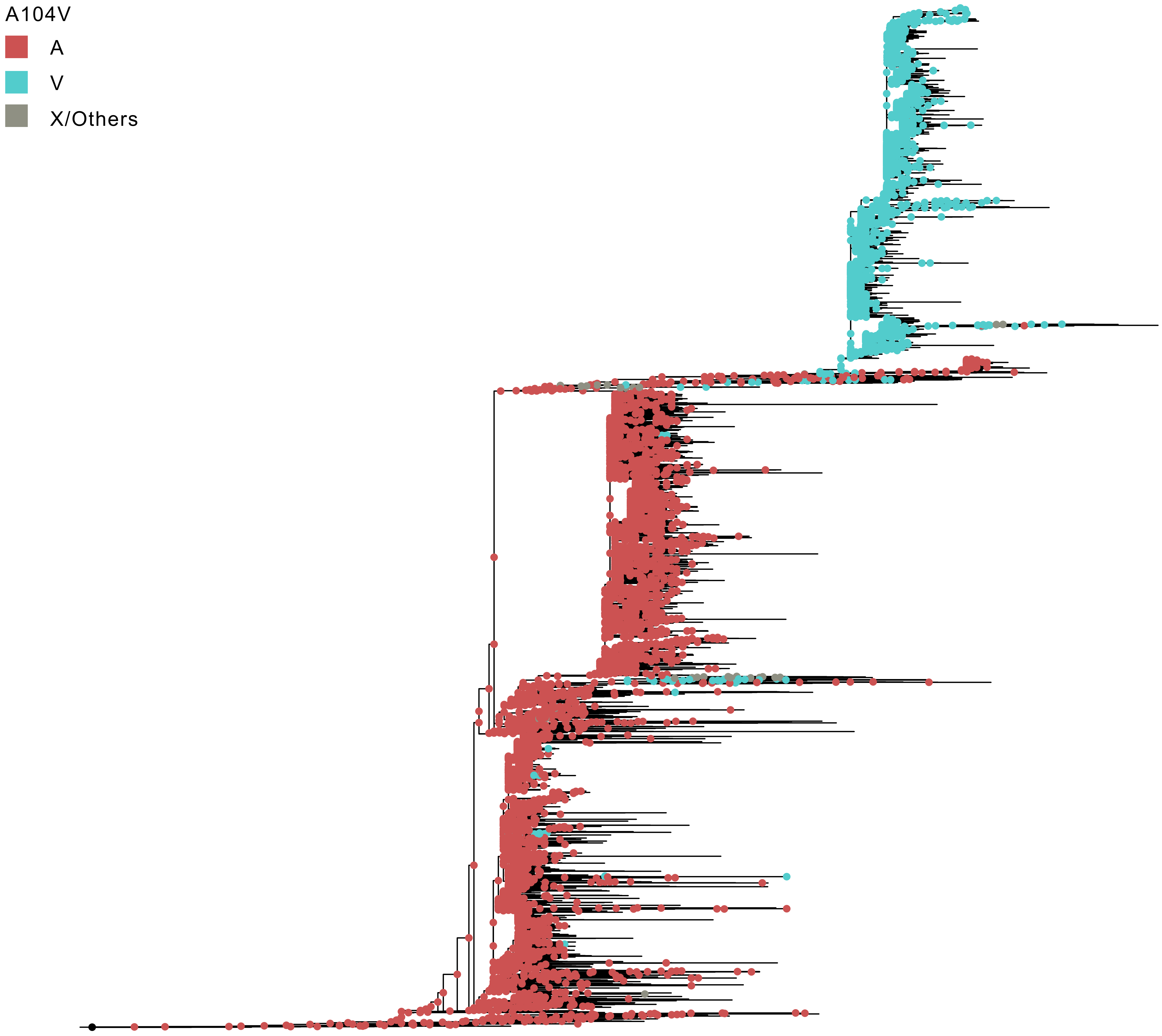
A104V
A
V
X/Others

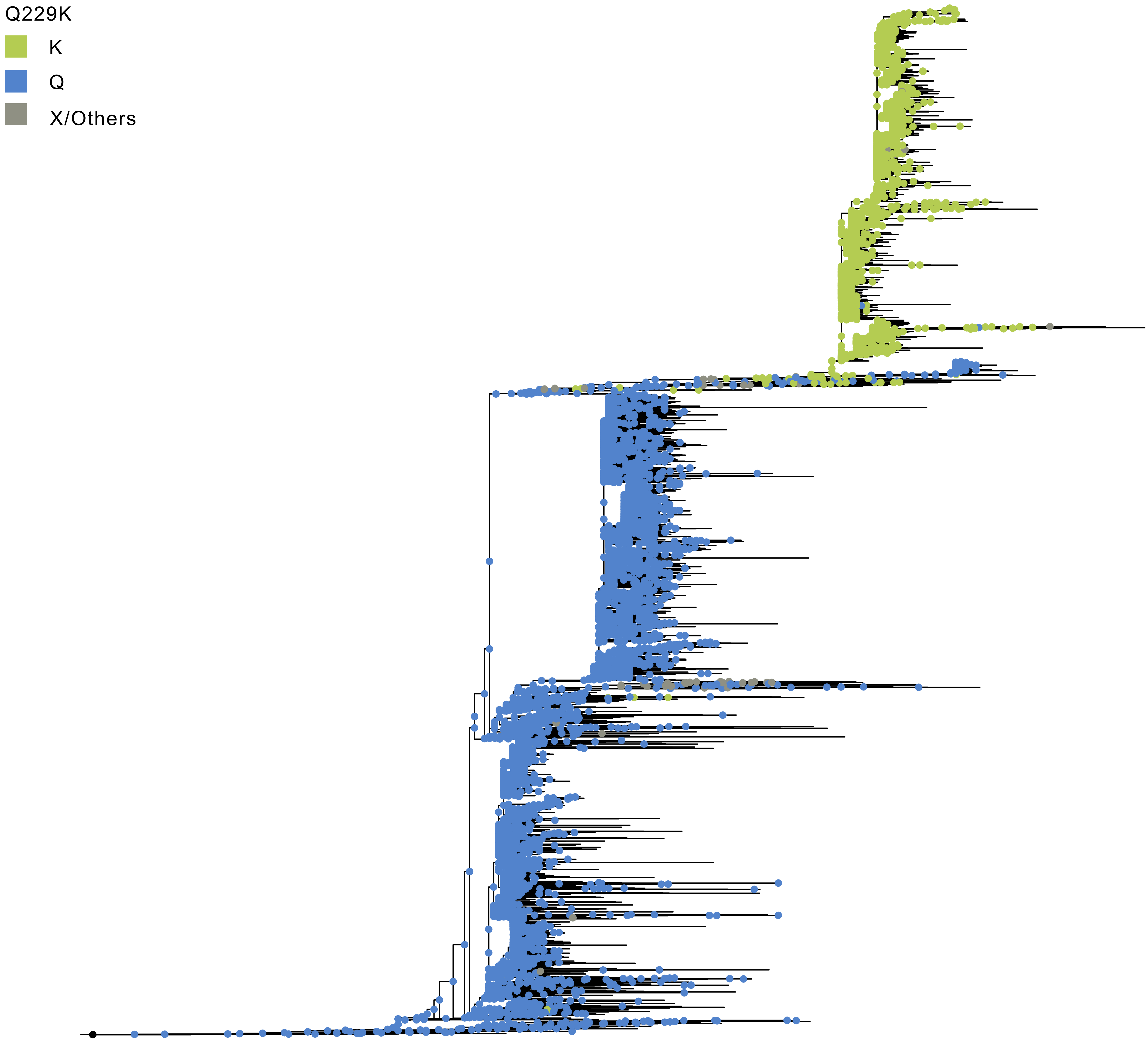
Q229K
K
Q
X/Others
