## Supplementary Data 3 for "Early evolution of BA.2.86 sheds light on the origins of highly divergent SARS-CoV-2 lineages"

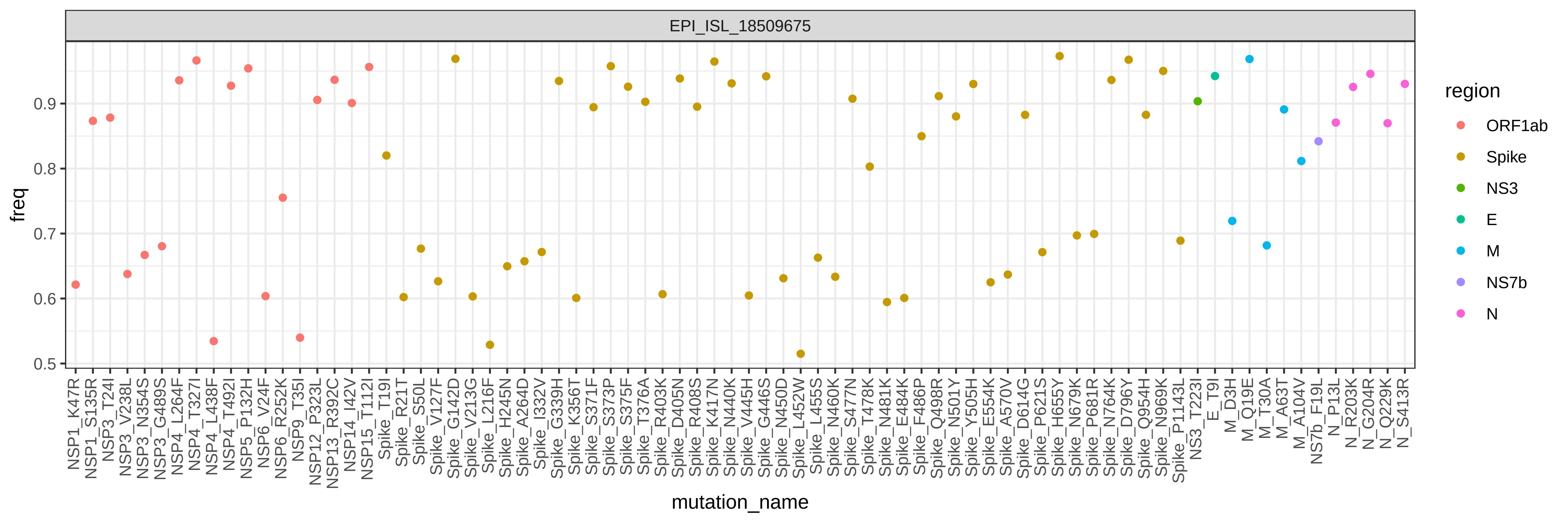

EPI_ISL_18509675
freq
mutation_name
region
ORF1ab
Spike
NS3
E
M
NS7b
N

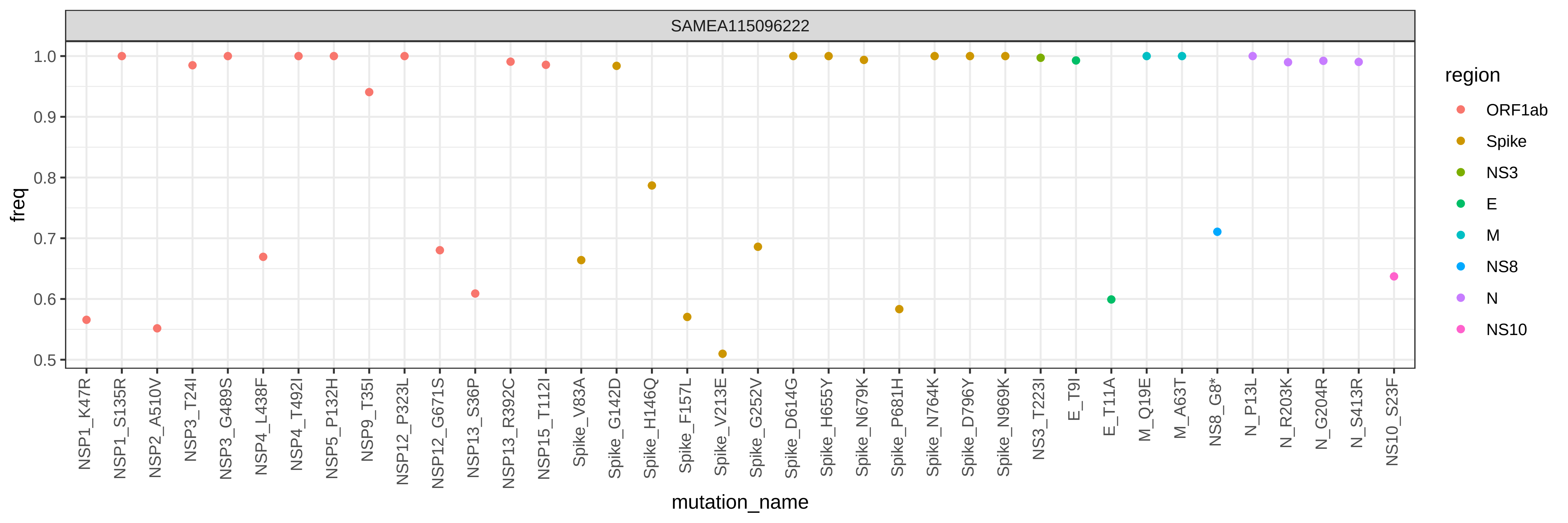

SAMEA115096222
freq
1.0
0.9
0.8
0.7
0.6
0.5
mutation_name
NSP1_K47R
NSP1_S135R
NSP2_A510V
NSP3_T24I
NSP3_G489S
NSP4_L438F
NSP4_T492I
NSP5_P132H
NSP9_T35I
NSP12_P323L
NSP12_G671S
NSP13_S36P
NSP13_R392C
NSP15_T112I
Spike_V83A
Spike_G142D
Spike_H146Q
Spike_F157L
Spike_V213E
Spike_G252V
Spike_D614G
Spike_H655Y
Spike_N679K
Spike_P681H
Spike_N764K
Spike_D796Y
Spike_N969K
NS3_T223I
E_T9I
E_T11A
M_Q19E
M_A63T
NS8_G8*
N_P13L
N_R203K
N_G204R
N_S413R
NS10_S23F
region
ORF1ab
Spike
NS3
E
M
NS8
N
NS10

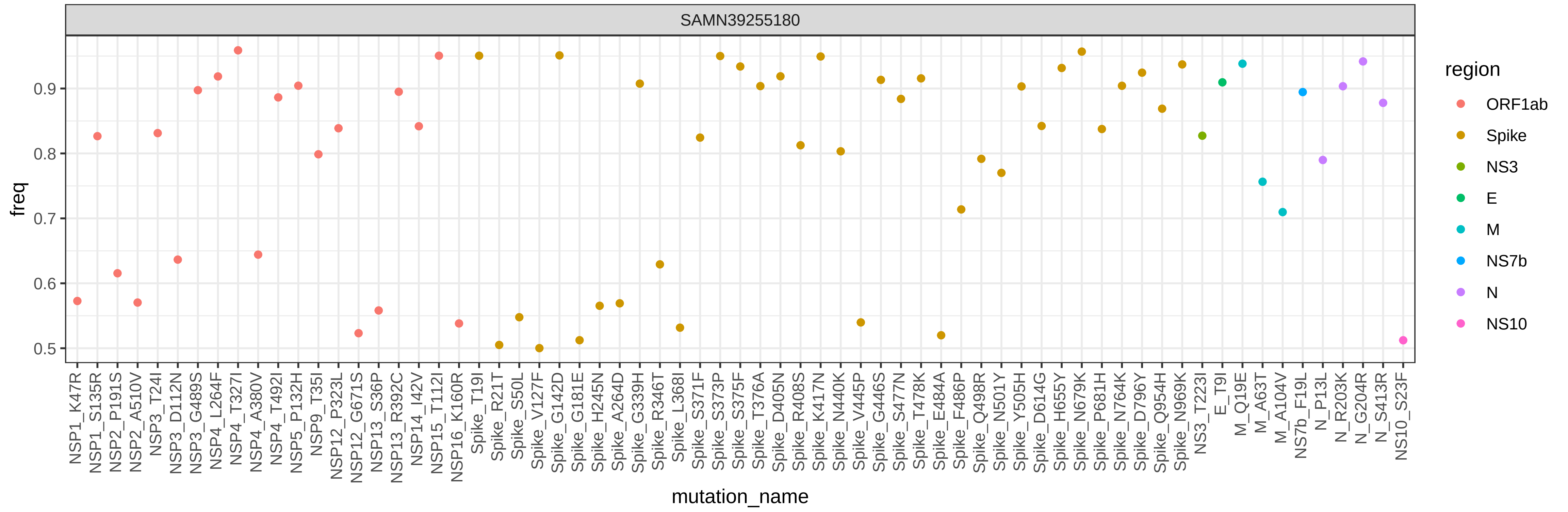
SAMN39255180
freq
0.5
0.6
0.7
0.8
0.9
mutation_name
NSP1_K47R
NSP1_S135R
NSP2_P191S
NSP2_A510V
NSP3_T24I
NSP3_D112N
NSP3_G489S
NSP4_L264F
NSP4_T327I
NSP4_A380V
NSP4_T492I
NSP5_P132H
NSP9_T35I
NSP12_P323L
NSP12_G671S
NSP13_S36P
NSP13_R392C
NSP14_I42V
NSP15_T112I
NSP16_K160R
Spike_T19I
Spike_R21T
Spike_S50L
Spike_V127F
Spike_G142D
Spike_G181E
Spike_H245N
Spike_A264D
Spike_G339H
Spike_R346T
Spike_L368I
Spike_S371F
Spike_S373P
Spike_S375F
Spike_T376A
Spike_D405N
Spike_R408S
Spike_K417N
Spike_N440K
Spike_V445P
Spike_G446S
Spike_S477N
Spike_T478K
Spike_E484A
Spike_F486P
Spike_Q498R
Spike_N501Y
Spike_Y505H
Spike_D614G
Spike_H655Y
Spike_N679K
Spike_P681H
Spike_N764K
Spike_D796Y
Spike_Q954H
Spike_N969K
NS3_T223I
E_T9I
M_Q19E
M_A63T
M_A104V
NS7b_F19L
N_P13L
N_R203K
N_G204R
N_S413R
NS10_S23F
region
ORF1ab
Spike
NS3
E
M
NS7b
N
NS10

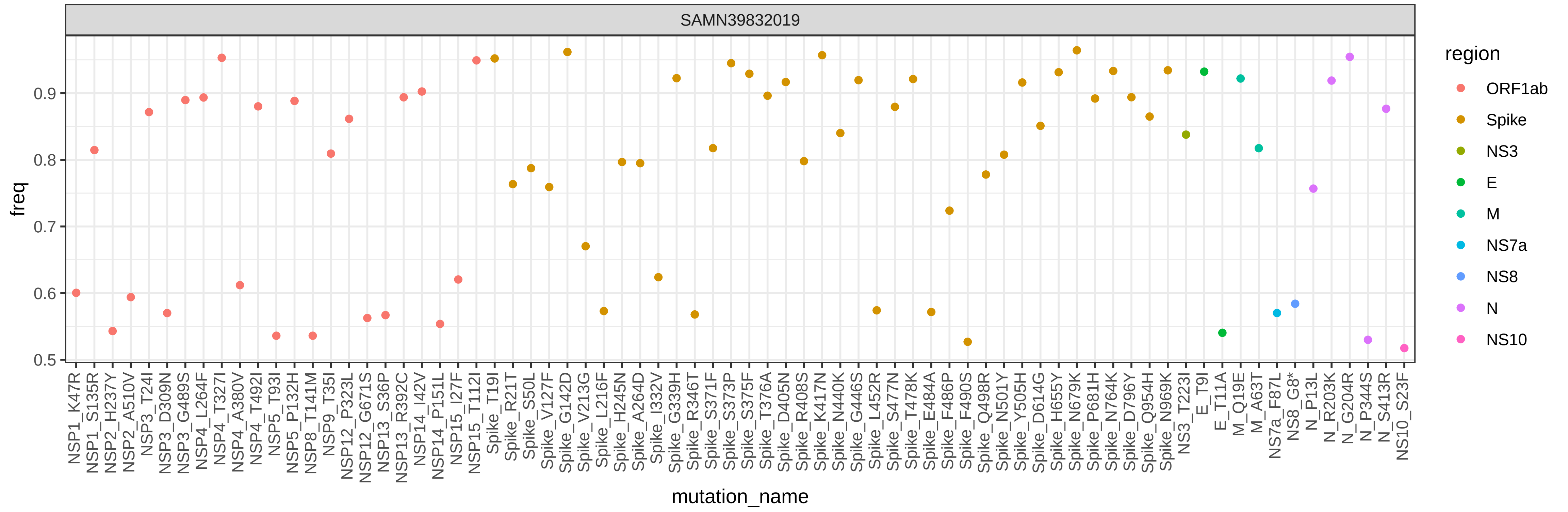

SAMN39832019
freq
0.5
0.6
0.7
0.8
0.9
mutation_name
NSP1_K47R
NSP1_S135R
NSP2_H237Y
NSP2_A510V
NSP3_T24I
NSP3_D309N
NSP3_G489S
NSP4_L264F
NSP4_T327I
NSP4_A380V
NSP4_T492I
NSP5_T93I
NSP5_P132H
NSP8_T141M
NSP9_T35I
NSP12_P323L
NSP12_G671S
NSP13_S36P
NSP13_R392C
NSP14_I42V
NSP14_P151L
NSP15_I27F
NSP15_T112I
Spike_T19I
Spike_R21T
Spike_S50L
Spike_V127F
Spike_G142D
Spike_V213G
Spike_L216F
Spike_H245N
Spike_A264D
Spike_I332V
Spike_G339H
Spike_R346T
Spike_S371F
Spike_S373P
Spike_S375F
Spike_T376A
Spike_D405N
Spike_R408S
Spike_K417N
Spike_N440K
Spike_G446S
Spike_L452R
Spike_S477N
Spike_T478K
Spike_E484A
Spike_F486P
Spike_F490S
Spike_Q498R
Spike_N501Y
Spike_Y505H
Spike_D614G
Spike_H655Y
Spike_N679K
Spike_P681H
Spike_N764K
Spike_D796Y
Spike_Q954H
Spike_N969K
NS3_T223I
E_T9I
E_T11A
M_Q19E
M_A63T
NS7a_F87L
NS8_G8*
N_P13L
N_R203K
N_G204R
N_P344S
N_S413R
NS10_S23F
region
ORF1ab
Spike
NS3
E
M
NS7a
NS8
N
NS10

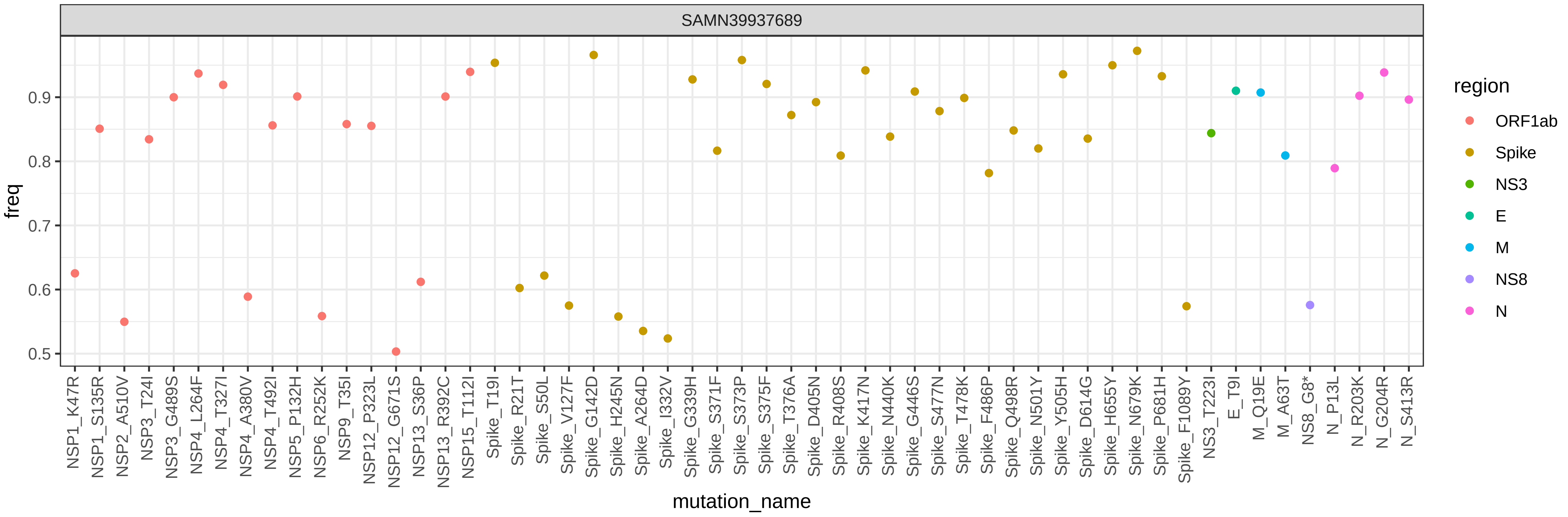

SAMN39937689
freq
0.5
0.6
0.7
0.8
0.9
NSP1_K47R
NSP1_S135R
NSP2_A510V
NSP3_T24I
NSP3_G489S
NSP4_L264F
NSP4_T327I
NSP4_A380V
NSP4_T492I
NSP5_P132H
NSP6_R252K
NSP9_T35I
NSP12_P323L
NSP12_G671S
NSP13_S36P
NSP13_R392C
NSP15_T112I
Spike_T19I
Spike_R21T
Spike_S50L
Spike_V127F
Spike_G142D
Spike_H245N
Spike_A264D
Spike_I332V
Spike_G339H
Spike_S371F
Spike_S373P
Spike_S375F
Spike_T376A
Spike_D405N
Spike_R408S
Spike_K417N
Spike_N440K
Spike_G446S
Spike_S477N
Spike_T478K
Spike_F486P
Spike_Q498R
Spike_N501Y
Spike_Y505H
Spike_D614G
Spike_H655Y
Spike_N679K
Spike_P681H
Spike_F1089Y
NS3_T223I
E_T9I
M_Q19E
M_A63T
NS8_G8*
N_P13L
N_R203K
N_G204R
N_S413R
mutation_name
region
ORF1ab
Spike
NS3
E
M
NS8
N

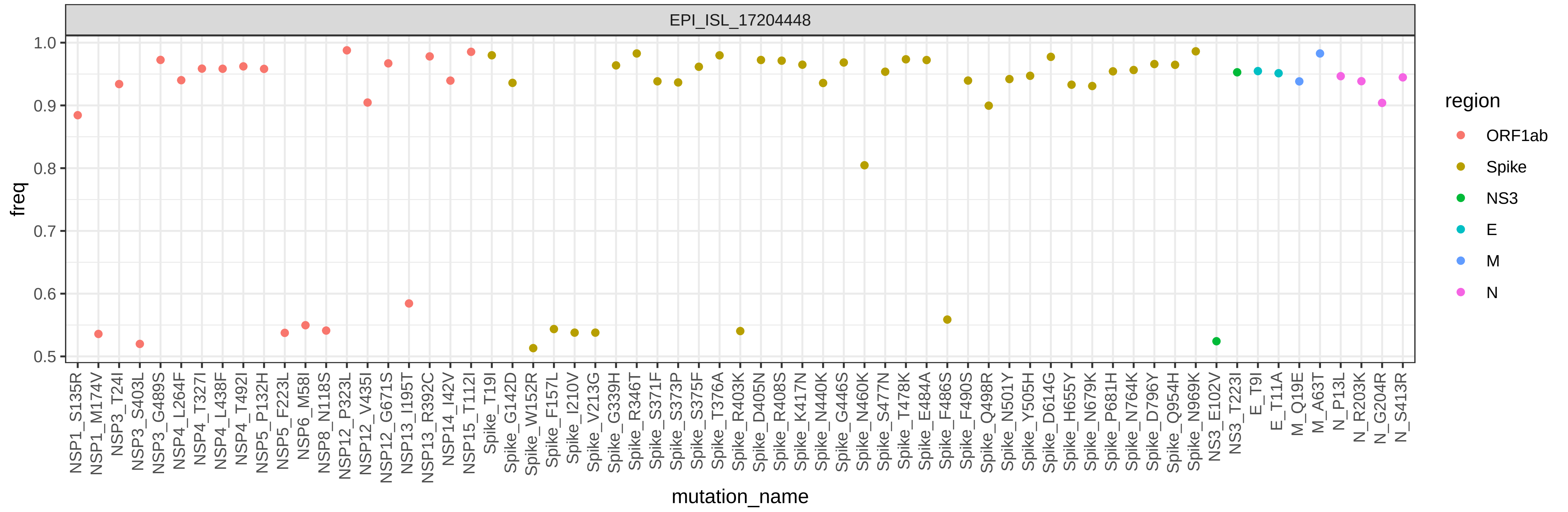

EPI_ISL_17204448
freq
1.0
0.9
0.8
0.7
0.6
0.5
mutation_name
NSP1_S135R
NSP1_M174V
NSP3_T24I
NSP3_S403L
NSP3_G489S
NSP4_L264F
NSP4_T327I
NSP4_L438F
NSP4_T492I
NSP5_P132H
NSP5_F223L
NSP6_M58I
NSP8_N118S
NSP12_P323L
NSP12_V435I
NSP12_G671S
NSP13_I195T
NSP13_R392C
NSP14_I42V
NSP15_T112I
Spike_T19I
Spike_G142D
Spike_W152R
Spike_F157L
Spike_I210V
Spike_V213G
Spike_G339H
Spike_R346T
Spike_S371F
Spike_S373P
Spike_S375F
Spike_T376A
Spike_R403K
Spike_D405N
Spike_R408S
Spike_K417N
Spike_N440K
Spike_G446S
Spike_N460K
Spike_S477N
Spike_T478K
Spike_E484A
Spike_F486S
Spike_F490S
Spike_Q498R
Spike_N501Y
Spike_Y505H
Spike_D614G
Spike_H655Y
Spike_N679K
Spike_P681H
Spike_N764K
Spike_D796Y
Spike_Q954H
Spike_N969K
NS3_E102V
NS3_T223I
E_T9I
E_T11A
M_Q19E
M_A63T
N_P13L
N_R203K
N_G204R
N_S413R
region
ORF1ab
Spike
NS3
E
M
N

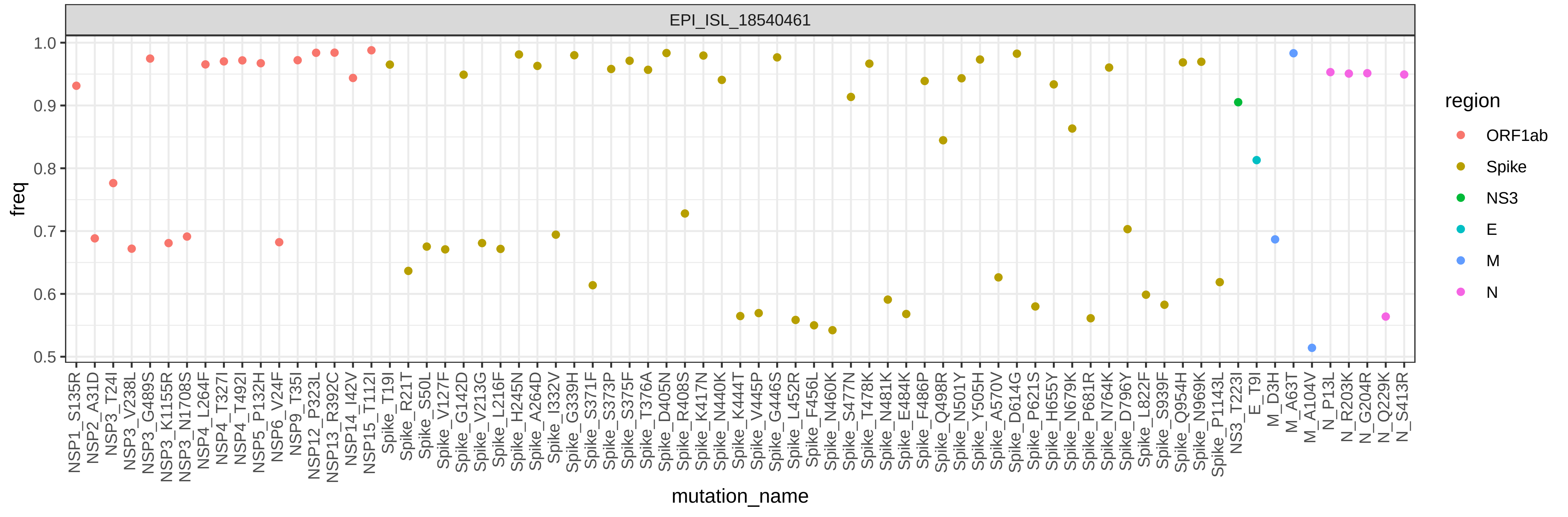

EPI_ISL_18540461
freq
1.0
0.9
0.8
0.7
0.6
0.5
mutation_name
NSP1_S135R
NSP2_A31D
NSP3_T24I
NSP3_V238L
NSP3_G489S
NSP3_K1155R
NSP3_N1708S
NSP4_L264F
NSP4_T327I
NSP4_T492I
NSP5_P132H
NSP6_V24F
NSP9_T35I
NSP12_P323L
NSP13_R392C
NSP14_I42V
NSP15_T112I
Spike_T19I
Spike_R21T
Spike_S50L
Spike_V127F
Spike_G142D
Spike_V213G
Spike_L216F
Spike_H245N
Spike_A264D
Spike_I332V
Spike_G339H
Spike_S371F
Spike_S373P
Spike_S375F
Spike_T376A
Spike_D405N
Spike_R408S
Spike_K417N
Spike_N440K
Spike_K444T
Spike_V445P
Spike_G446S
Spike_L452R
Spike_F456L
Spike_N460K
Spike_S477N
Spike_T478K
Spike_N481K
Spike_E484K
Spike_F486P
Spike_Q498R
Spike_N501Y
Spike_Y505H
Spike_A570V
Spike_D614G
Spike_P621S
Spike_H655Y
Spike_N679K
Spike_P681R
Spike_N764K
Spike_D796Y
Spike_L822F
Spike_S939F
Spike_Q954H
Spike_N969K
Spike_P1143L
NS3_T223I
E_T9I
M_D3H
M_A63T
M_A104V
N_P13L
N_R203K
N_G204R
N_Q229K
N_S413R
region
ORF1ab
Spike
NS3
E
M
N

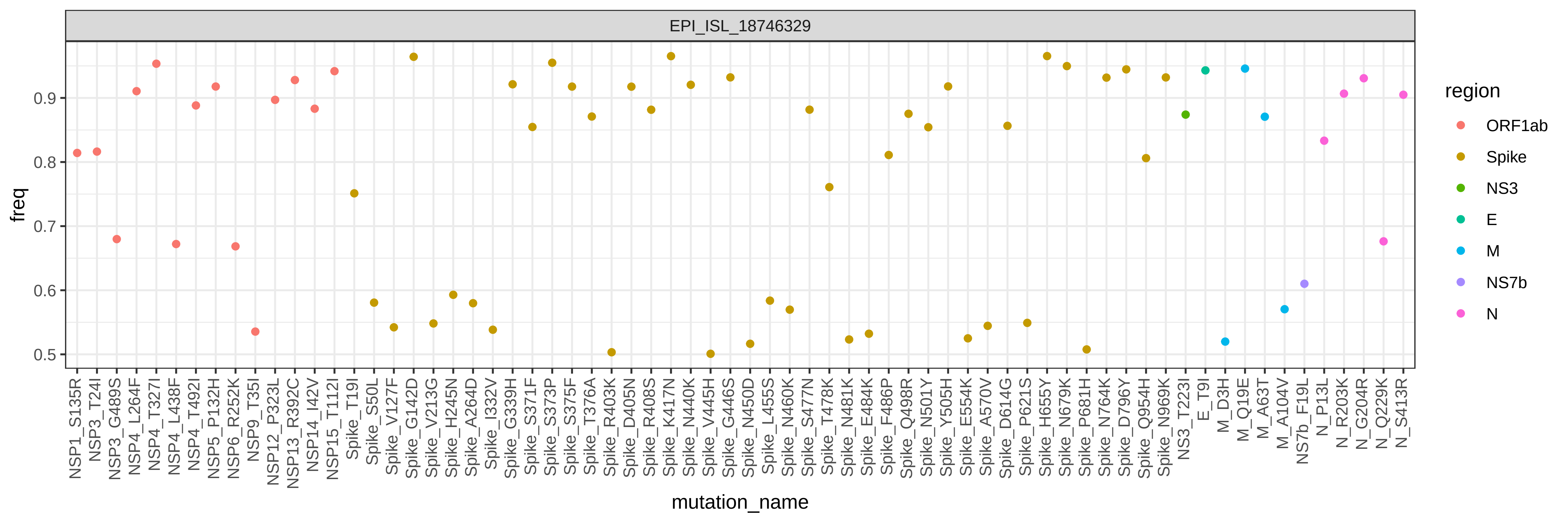

EPI_ISL_18746329
freq
0.5
0.6
0.7
0.8
0.9
mutation_name
region
ORF1ab
Spike
NS3
E
M
NS7b
N
NSP1_S135R
NSP3_T24I
NSP3_G489S
NSP4_L264F
NSP4_T327I
NSP4_L438F
NSP4_T492I
NSP5_P132H
NSP6_R252K
NSP9_T35I
NSP12_P323L
NSP13_R392C
NSP14_I42V
NSP15_T112I
Spike_T19I
Spike_S50L
Spike_V127F
Spike_G142D
Spike_V213G
Spike_H245N
Spike_A264D
Spike_I332V
Spike_G339H
Spike_S371F
Spike_S373P
Spike_S375F
Spike_T376A
Spike_R403K
Spike_D405N
Spike_R408S
Spike_K417N
Spike_N440K
Spike_V445H
Spike_G446S
Spike_N450D
Spike_L455S
Spike_N460K
Spike_S477N
Spike_T478K
Spike_N481K
Spike_E484K
Spike_F486P
Spike_Q498R
Spike_N501Y
Spike_Y505H
Spike_E554K
Spike_A570V
Spike_D614G
Spike_P621S
Spike_H655Y
Spike_N679K
Spike_P681H
Spike_N764K
Spike_D796Y
Spike_Q954H
Spike_N969K
NS3_T223I
E_T9I
M_D3H
M_Q19E
M_A63T
M_A104V
NS7b_F19L
N_P13L
N_R203K
N_G204R
N_Q229K
N_S413R

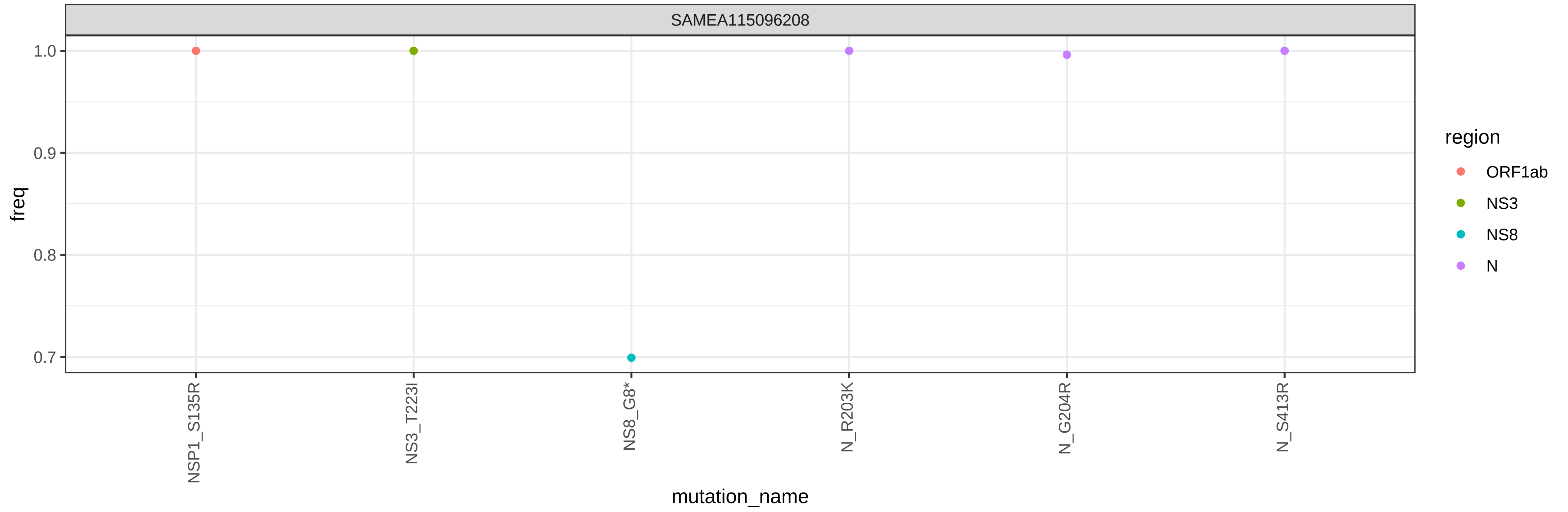
SAMEA115096208
freq
1.0
0.9
0.8
0.7
NSP1_S135R
NS3_T223I
NS8_G8*
N_R203K
N_G204R
N_S413R
mutation_name
region
ORF1ab
NS3
NS8
N

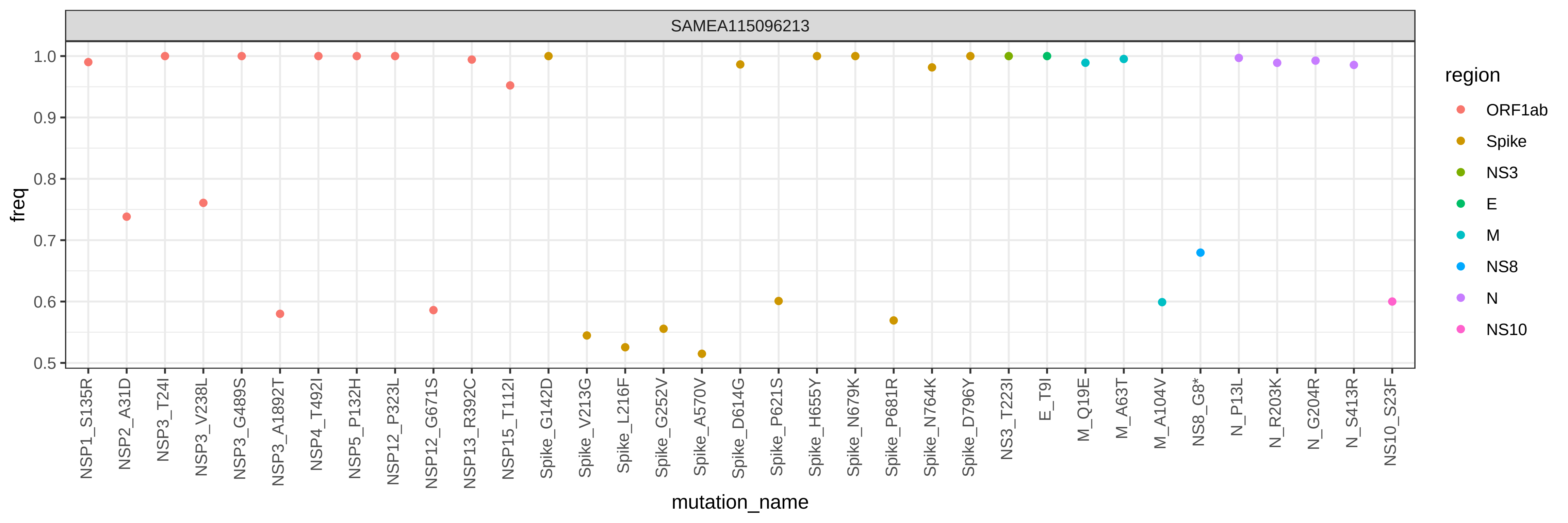

SAMEA115096213
freq
1.0
0.9
0.8
0.7
0.6
0.5
NSP1_S135R
NSP2_A31D
NSP3_T24I
NSP3_V238L
NSP3_G489S
NSP3_A1892T
NSP4_T492I
NSP5_P132H
NSP12_P323L
NSP12_G671S
NSP13_R392C
NSP15_T112I
Spike_G142D
Spike_V213G
Spike_L216F
Spike_G252V
Spike_A570V
Spike_D614G
Spike_P621S
Spike_H655Y
Spike_N679K
Spike_P681R
Spike_N764K
Spike_D796Y
NS3_T223I
E_T9I
M_Q19E
M_A63T
M_A104V
NS8_G8*
N_P13L
N_R203K
N_G204R
N_S413R
NS10_S23F
mutation_name
region
ORF1ab
Spike
NS3
E
M
NS8
N
NS10

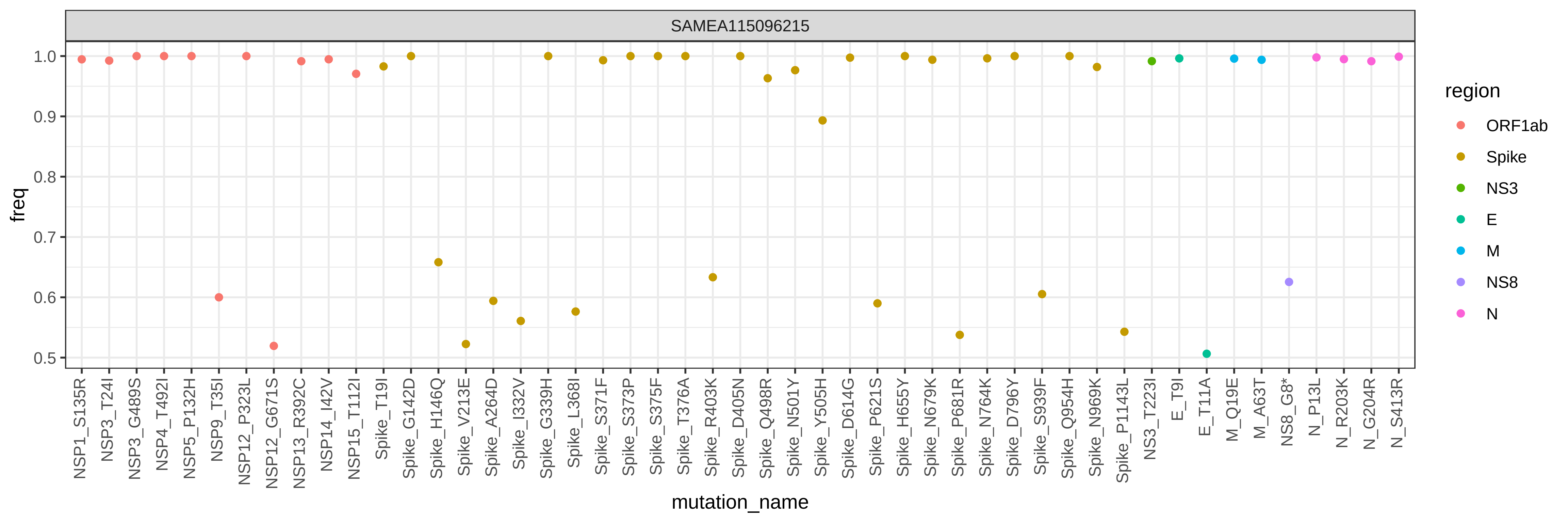
SAMEA115096215
freq
1.0
0.9
0.8
0.7
0.6
0.5
mutation_name
NSP1_S135R
NSP3_T24I
NSP3_G489S
NSP4_T492I
NSP5_P132H
NSP9_T35I
NSP12_P323L
NSP12_G671S
NSP13_R392C
NSP14_I42V
NSP15_T112I
Spike_T19I
Spike_G142D
Spike_H146Q
Spike_V213E
Spike_A264D
Spike_I332V
Spike_G339H
Spike_L368I
Spike_S371F
Spike_S373P
Spike_S375F
Spike_T376A
Spike_R403K
Spike_D405N
Spike_Q498R
Spike_N501Y
Spike_Y505H
Spike_D614G
Spike_P621S
Spike_H655Y
Spike_N679K
Spike_P681R
Spike_N764K
Spike_D796Y
Spike_S939F
Spike_Q954H
Spike_N969K
Spike_P1143L
NS3_T223I
E_T9I
E_T11A
M_Q19E
M_A63T
NS8_G8*
N_P13L
N_R203K
N_G204R
N_S413R
region
ORF1ab
Spike
NS3
E
M
NS8
N

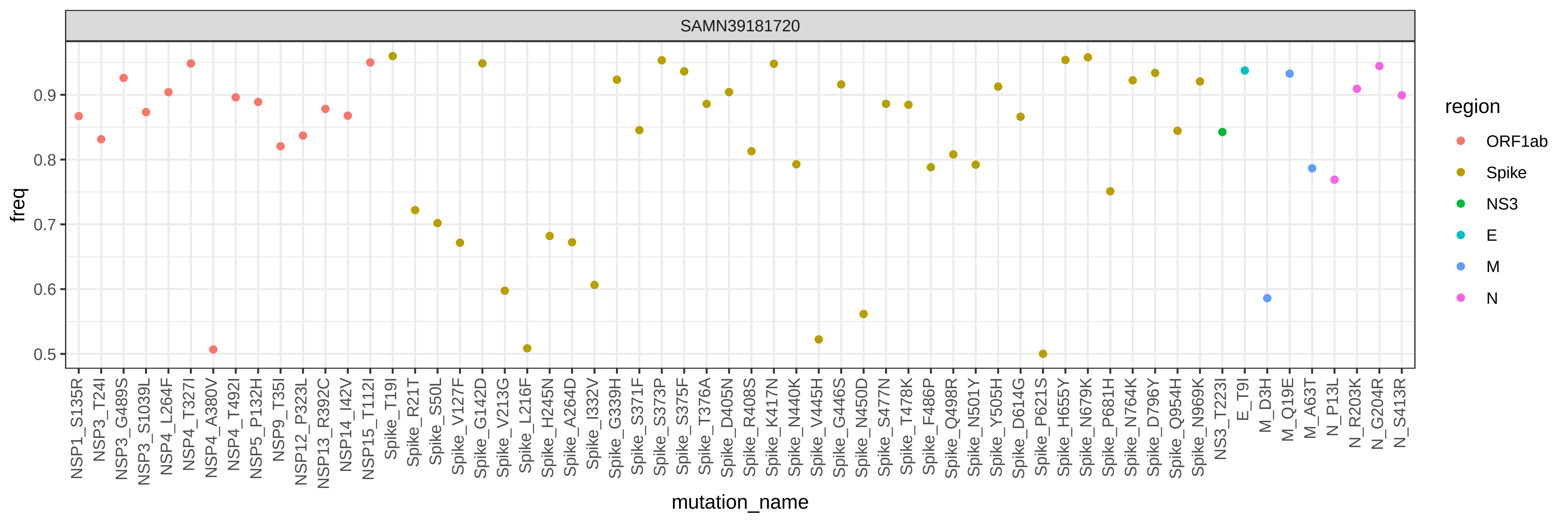

SAMN39181720
freq
0.5
0.6
0.7
0.8
0.9
mutation_name
NSP1_S135R
NSP3_T24I
NSP3_G489S
NSP3_S1039L
NSP4_L264F
NSP4_T327I
NSP4_A380V
NSP4_T492I
NSP5_P132H
NSP9_T35I
NSP12_P323L
NSP13_R392C
NSP14_I42V
NSP15_T112I
Spike_T19I
Spike_R21T
Spike_S50L
Spike_V127F
Spike_G142D
Spike_V213G
Spike_L216F
Spike_H245N
Spike_A264D
Spike_I332V
Spike_G339H
Spike_S371F
Spike_S373P
Spike_S375F
Spike_T376A
Spike_D405N
Spike_R408S
Spike_K417N
Spike_N440K
Spike_V445H
Spike_G446S
Spike_N450D
Spike_S477N
Spike_T478K
Spike_F486P
Spike_Q498R
Spike_N501Y
Spike_Y505H
Spike_D614G
Spike_P621S
Spike_H655Y
Spike_N679K
Spike_P681H
Spike_N764K
Spike_D796Y
Spike_Q954H
Spike_N969K
NS3_T223I
E_T9I
M_D3H
M_Q19E
M_A63T
N_P13L
N_R203K
N_G204R
N_S413R
region
ORF1ab
Spike
NS3
E
M
N

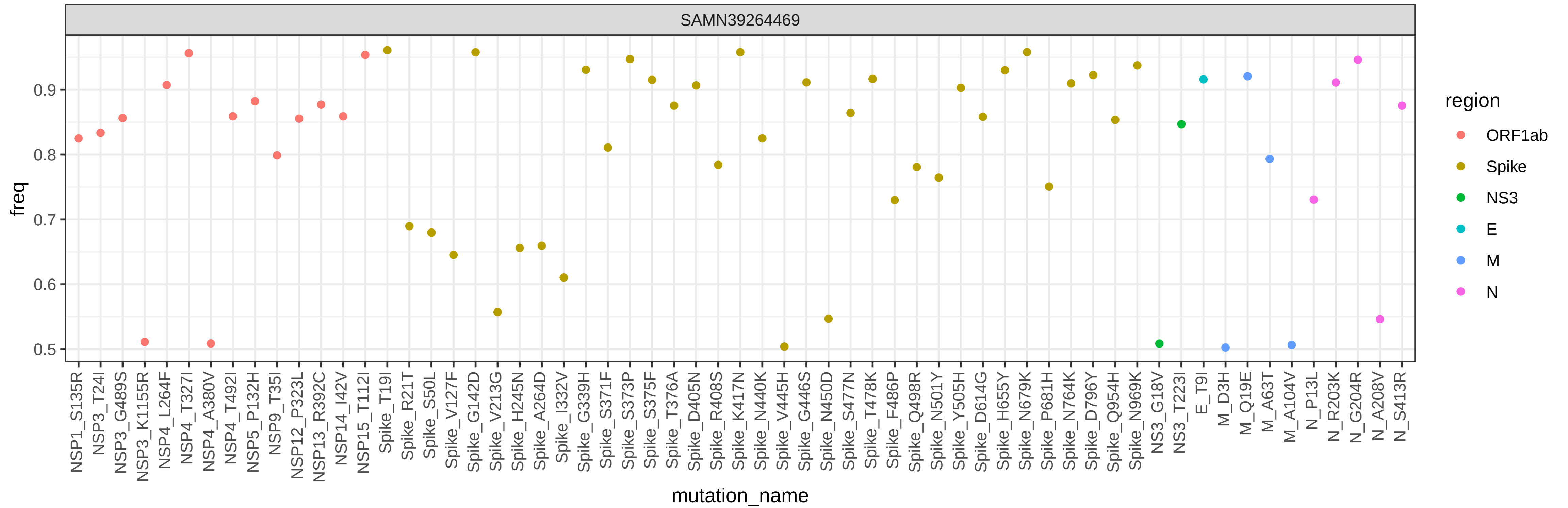

SAMN39264469
freq
0.5
0.6
0.7
0.8
0.9
mutation_name
NSP1_S135R
NSP3_T24I
NSP3_G489S
NSP3_K1155R
NSP4_L264F
NSP4_T327I
NSP4_A380V
NSP4_T492I
NSP5_P132H
NSP9_T35I
NSP12_P323L
NSP13_R392C
NSP14_I42V
NSP15_T112I
Spike_T19I
Spike_R21T
Spike_S50L
Spike_V127F
Spike_G142D
Spike_V213G
Spike_H245N
Spike_A264D
Spike_I332V
Spike_G339H
Spike_S371F
Spike_S373P
Spike_S375F
Spike_T376A
Spike_D405N
Spike_R408S
Spike_K417N
Spike_N440K
Spike_V445H
Spike_G446S
Spike_N450D
Spike_S477N
Spike_T478K
Spike_F486P
Spike_Q498R
Spike_N501Y
Spike_Y505H
Spike_D614G
Spike_H655Y
Spike_N679K
Spike_P681H
Spike_N764K
Spike_D796Y
Spike_Q954H
Spike_N969K
NS3_G18V
NS3_T223I
E_T9I
M_D3H
M_Q19E
M_A63T
M_A104V
N_P13L
N_R203K
N_G204R
N_A208V
N_S413R
region
ORF1ab
Spike
NS3
E
M
N

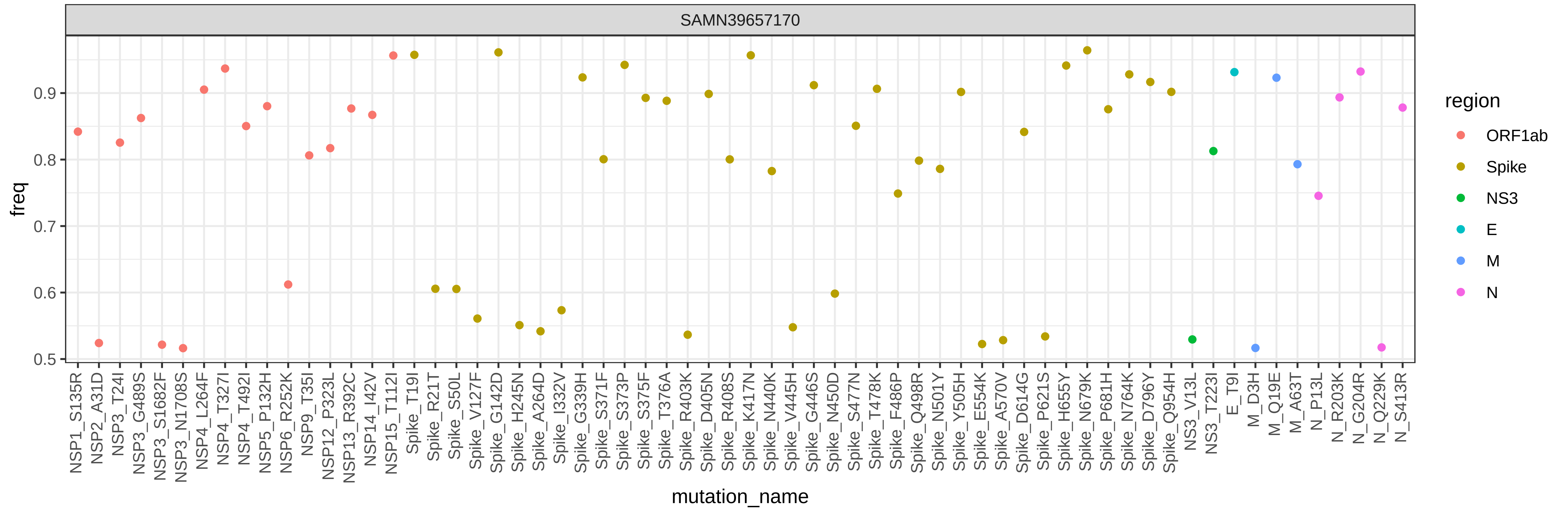

SAMN39657170
freq
0.5
0.6
0.7
0.8
0.9
mutation_name
NSP1_S135R
NSP2_A31D
NSP3_T24I
NSP3_G489S
NSP3_S1682F
NSP3_N1708S
NSP4_L264F
NSP4_T327I
NSP4_T492I
NSP5_P132H
NSP6_R252K
NSP9_T35I
NSP12_P323L
NSP13_R392C
NSP14_I42V
NSP15_T112I
Spike_T19I
Spike_R21T
Spike_S50L
Spike_V127F
Spike_G142D
Spike_H245N
Spike_A264D
Spike_I332V
Spike_G339H
Spike_S371F
Spike_S373P
Spike_S375F
Spike_T376A
Spike_R403K
Spike_D405N
Spike_R408S
Spike_K417N
Spike_N440K
Spike_V445H
Spike_G446S
Spike_N450D
Spike_S477N
Spike_T478K
Spike_F486P
Spike_Q498R
Spike_N501Y
Spike_Y505H
Spike_E554K
Spike_A570V
Spike_D614G
Spike_P621S
Spike_H655Y
Spike_N679K
Spike_P681H
Spike_N764K
Spike_D796Y
Spike_Q954H
NS3_V13L
NS3_T223I
E_T9I
M_D3H
M_Q19E
M_A63T
N_P13L
N_R203K
N_G204R
N_Q229K
N_S413R
region
ORF1ab
Spike
NS3
E
M
N

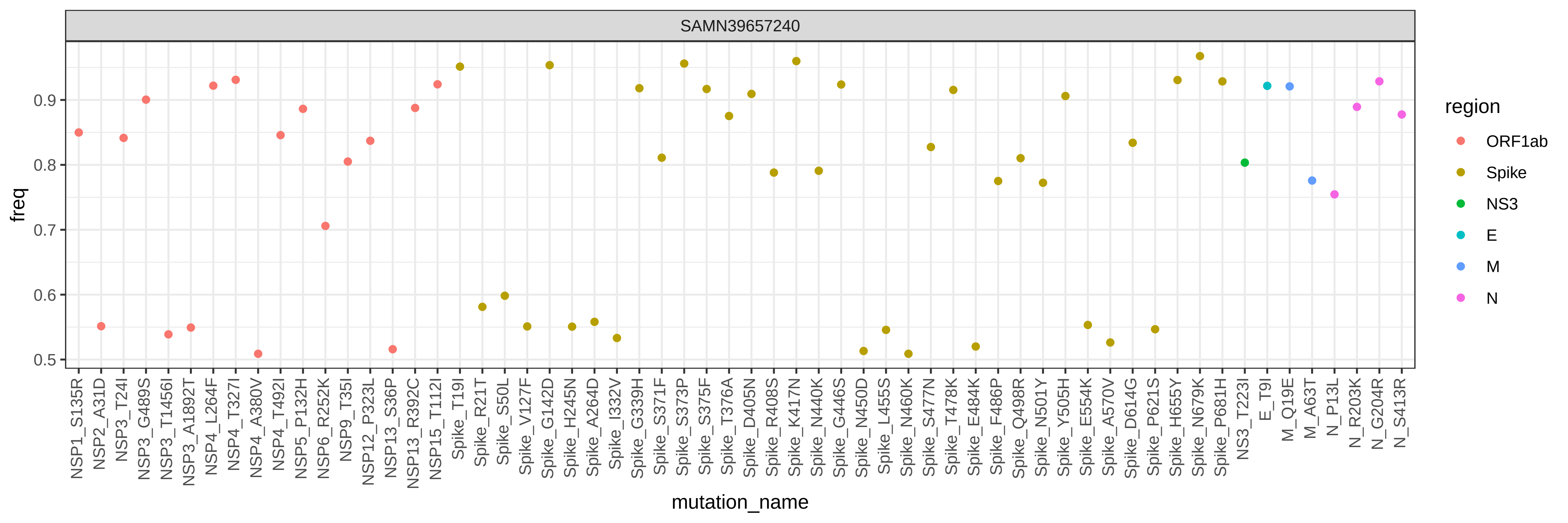

SAMN39657240
freq
0.5
0.6
0.7
0.8
0.9
mutation_name
NSP1_S135R
NSP2_A31D
NSP3_T24I
NSP3_G489S
NSP3_T1456I
NSP3_A1892T
NSP4_L264F
NSP4_T327I
NSP4_A380V
NSP4_T492I
NSP5_P132H
NSP6_R252K
NSP9_T35I
NSP12_P323L
NSP13_S36P
NSP13_R392C
NSP15_T112I
Spike_T19I
Spike_R21T
Spike_S50L
Spike_V127F
Spike_G142D
Spike_H245N
Spike_A264D
Spike_I332V
Spike_G339H
Spike_S371F
Spike_S373P
Spike_S375F
Spike_T376A
Spike_D405N
Spike_R408S
Spike_K417N
Spike_N440K
Spike_G446S
Spike_N450D
Spike_L455S
Spike_N460K
Spike_S477N
Spike_T478K
Spike_E484K
Spike_F486P
Spike_Q498R
Spike_N501Y
Spike_Y505H
Spike_E554K
Spike_A570V
Spike_D614G
Spike_P621S
Spike_H655Y
Spike_N679K
Spike_P681H
NS3_T223I
E_T9I
M_Q19E
M_A63T
N_P13L
N_R203K
N_G204R
N_S413R
region
ORF1ab
Spike
NS3
E
M
N

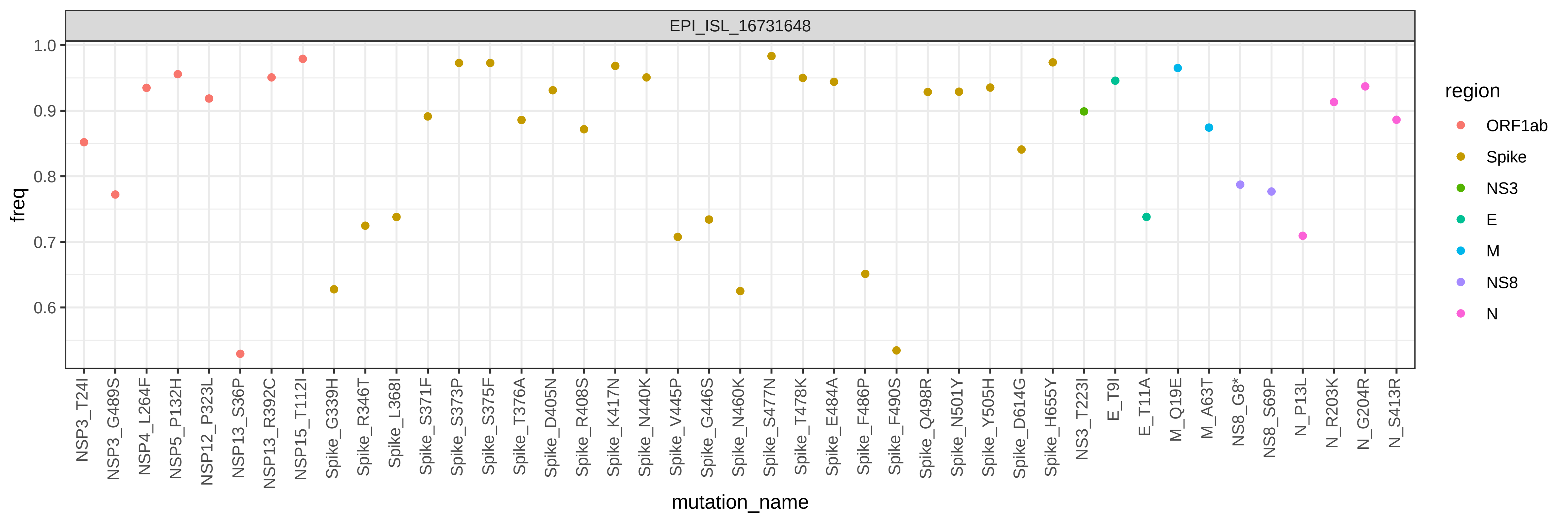

EPI_ISL_16731648
freq
1.0
0.9
0.8
0.7
0.6
mutation_name
NSP3_T24I
NSP3_G489S
NSP4_L264F
NSP5_P132H
NSP12_P323L
NSP13_S36P
NSP13_R392C
NSP15_T112I
Spike_G339H
Spike_R346T
Spike_L368I
Spike_S371F
Spike_S373P
Spike_S375F
Spike_T376A
Spike_D405N
Spike_R408S
Spike_K417N
Spike_N440K
Spike_V445P
Spike_G446S
Spike_N460K
Spike_S477N
Spike_T478K
Spike_E484A
Spike_F486P
Spike_F490S
Spike_Q498R
Spike_N501Y
Spike_Y505H
Spike_D614G
Spike_H655Y
NS3_T223I
E_T9I
E_T11A
M_Q19E
M_A63T
NS8_G8*
NS8_S69P
N_P13L
N_R203K
N_G204R
N_S413R
region
ORF1ab
Spike
NS3
E
M
NS8
N

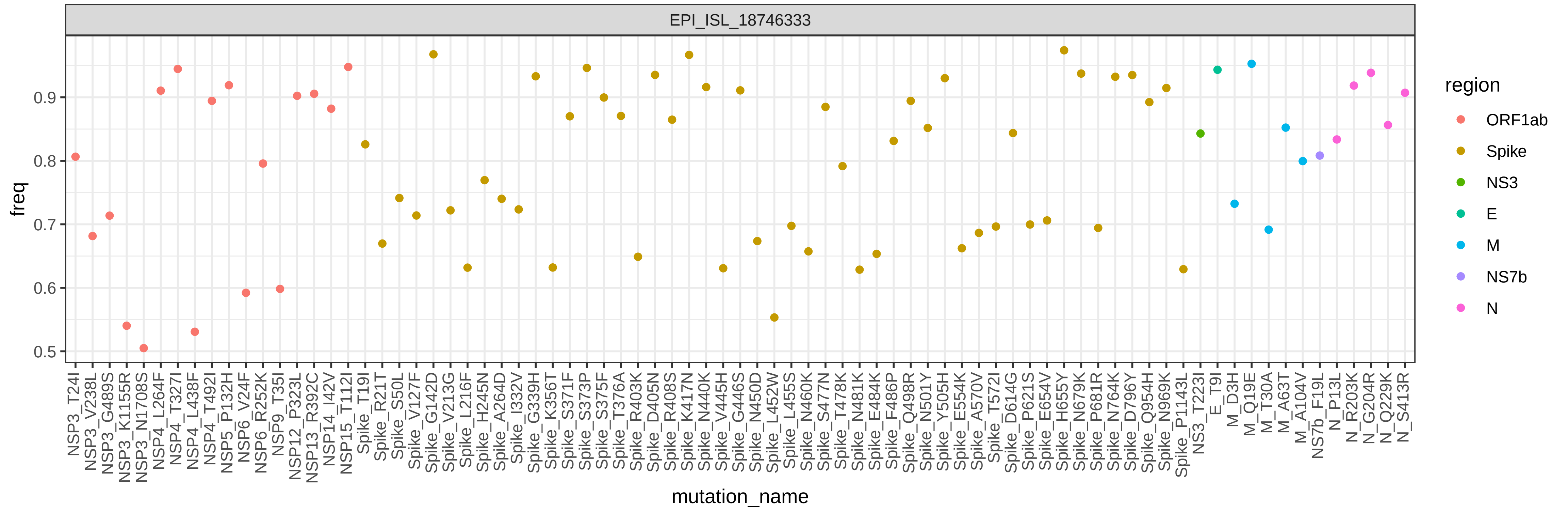
EPI_ISL_18746333
freq
mutation_name
region
ORF1ab
Spike
NS3
E
M
NS7b
N

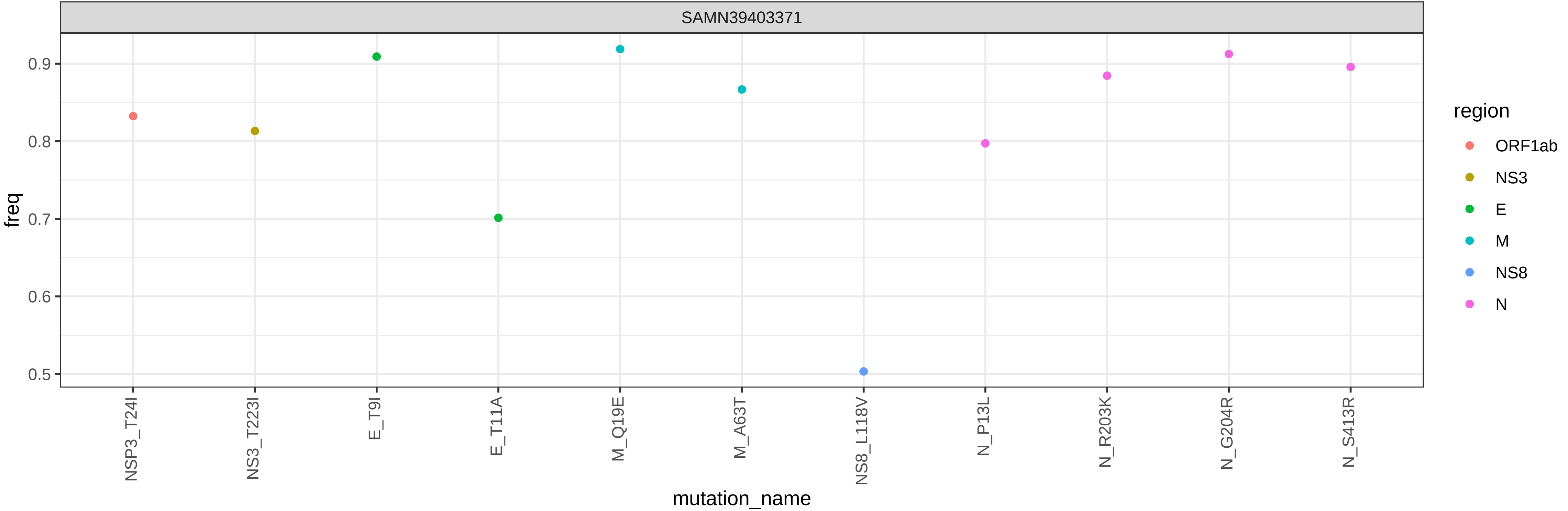

SAMN39403371
freq
0.9
0.8
0.7
0.6
0.5
NSP3_T24I
NS3_T223I
E_T9I
E_T11A
M_Q19E
M_A63T
NS8_L118V
N_P13L
N_R203K
N_G204R
N_S413R
mutation_name
region
ORF1ab
NS3
E
M
NS8
N

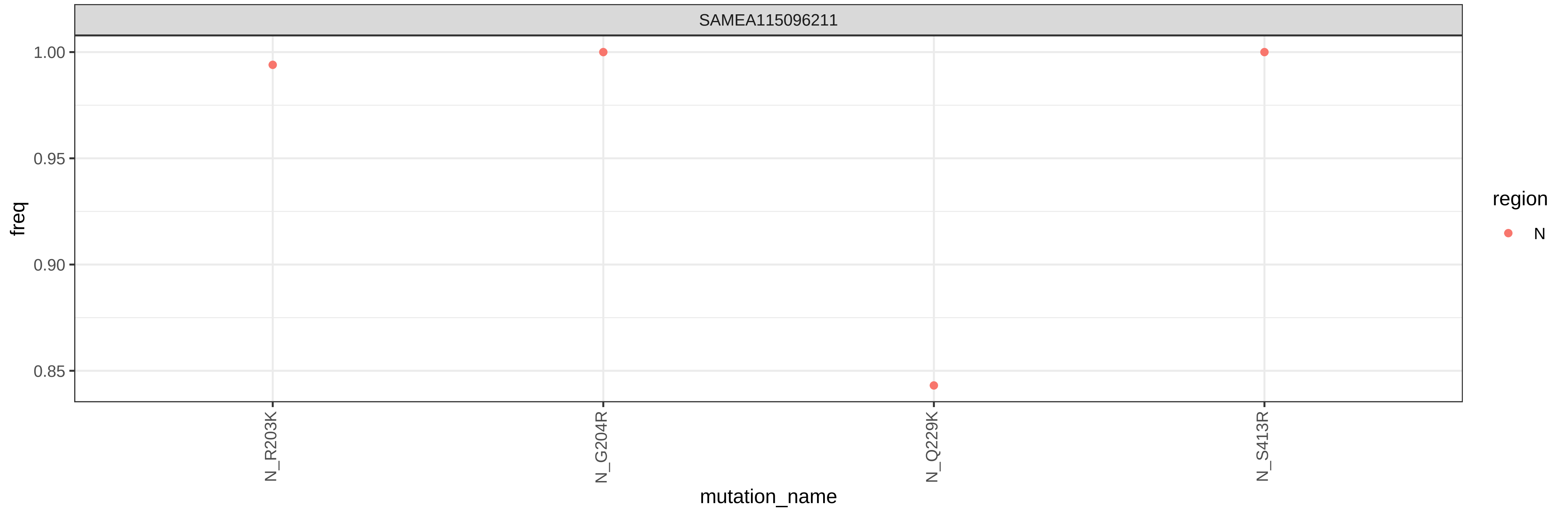

SAMEA115096211
1.00
0.95
0.90
0.85
freq
N_R203K
N_G204R
N_Q229K
N_S413R
mutation_name
region
N
