## Supplementary Data 4 for "Early evolution of BA.2.86 sheds light on the origins of highly divergent SARS-CoV-2 lineages"

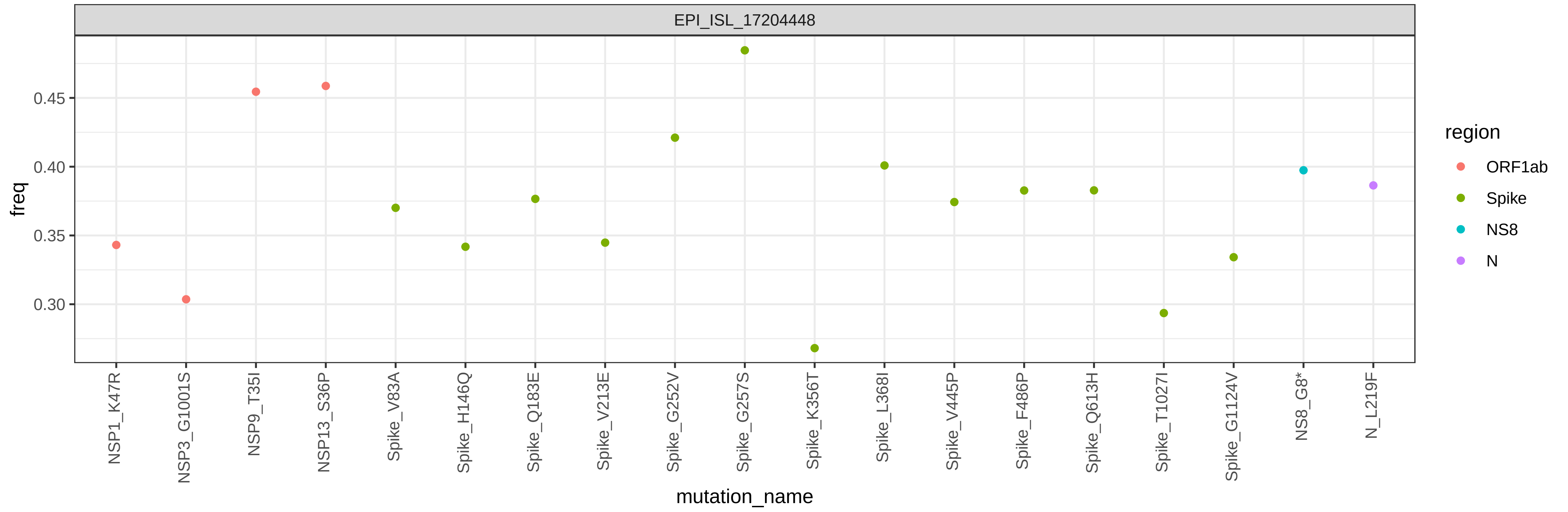
EPI_ISL_17204448
freq
0.45
0.40
0.35
0.30
mutation_name
NSP1_K47R
NSP3_G1001S
NSP9_T35I
NSP13_S36P
Spike_V83A
Spike_H146Q
Spike_Q183E
Spike_V213E
Spike_G252V
Spike_G257S
Spike_K356T
Spike_L368I
Spike_V445P
Spike_F486P
Spike_Q613H
Spike_T1027I
Spike_G1124V
NS8_G8*
N_L219F
region
ORF1ab
Spike
NS8
N

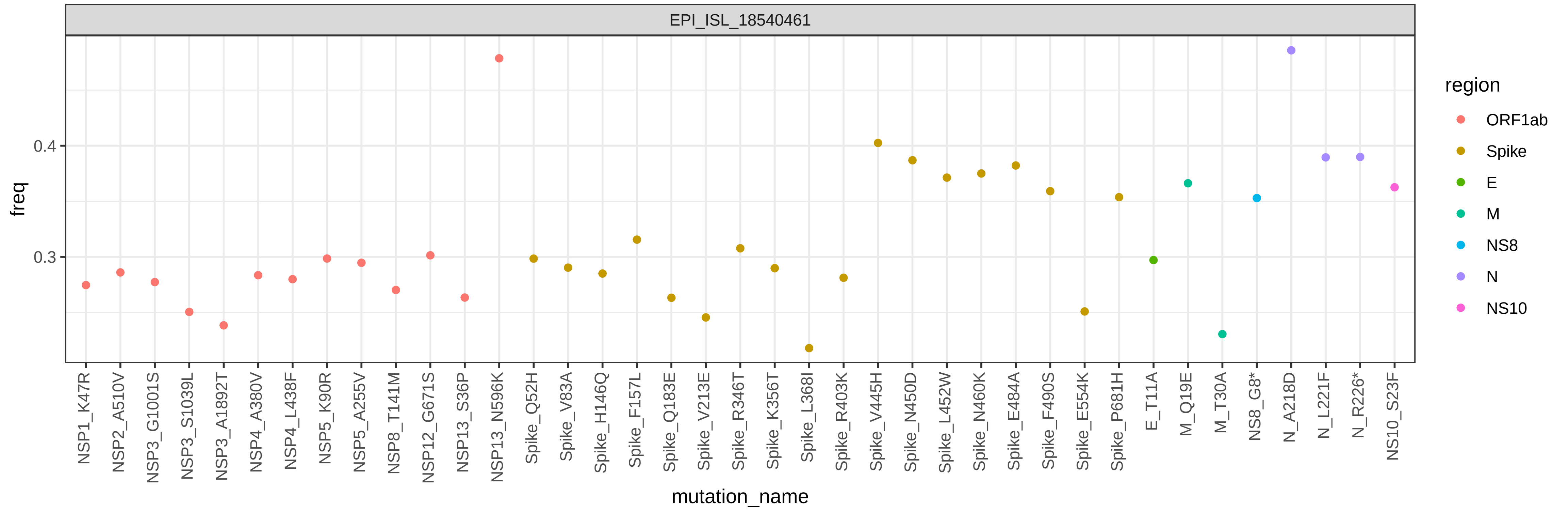

EPI_ISL_18540461
freq
0.4
0.3
mutation_name
NSP1_K47R
NSP2_A510V
NSP3_G1001S
NSP3_S1039L
NSP3_A1892T
NSP4_A380V
NSP4_L438F
NSP5_K90R
NSP5_A255V
NSP8_T141M
NSP12_G671S
NSP13_S36P
NSP13_N596K
Spike_Q52H
Spike_V83A
Spike_H146Q
Spike_F157L
Spike_Q183E
Spike_V213E
Spike_R346T
Spike_K356T
Spike_L368I
Spike_R403K
Spike_V445H
Spike_N450D
Spike_L452W
Spike_N460K
Spike_E484A
Spike_F490S
Spike_E554K
Spike_P681H
E_T11A
M_Q19E
M_T30A
NS8_G8*
N_A218D
N_L221F
N_R226*
NS10_S23F
region
ORF1ab
Spike
E
M
NS8
N
NS10

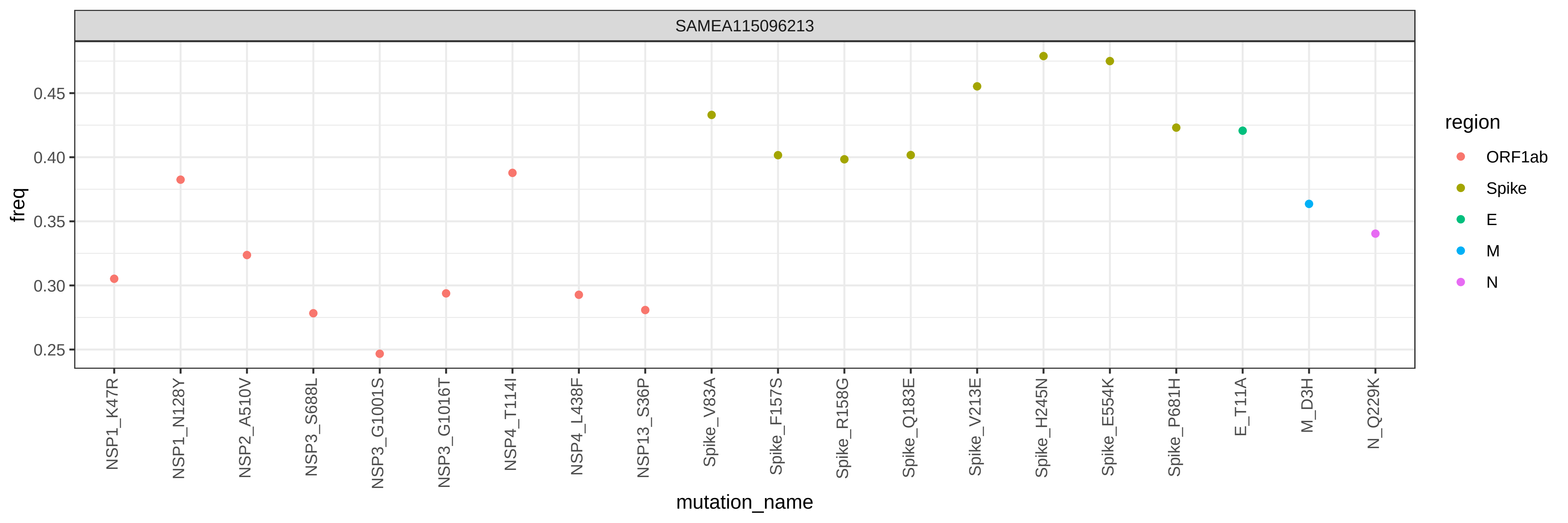
SAMEA115096213
freq
0.45
0.40
0.35
0.30
0.25
mutation_name
NSP1_K47R
NSP1_N128Y
NSP2_A510V
NSP3_S688L
NSP3_G1001S
NSP3_G1016T
NSP4_T114I
NSP4_L438F
NSP13_S36P
Spike_V83A
Spike_F157S
Spike_R158G
Spike_Q183E
Spike_V213E
Spike_H245N
Spike_E554K
Spike_P681H
E_T11A
M_D3H
N_Q229K
region
ORF1ab
Spike
E
M
N

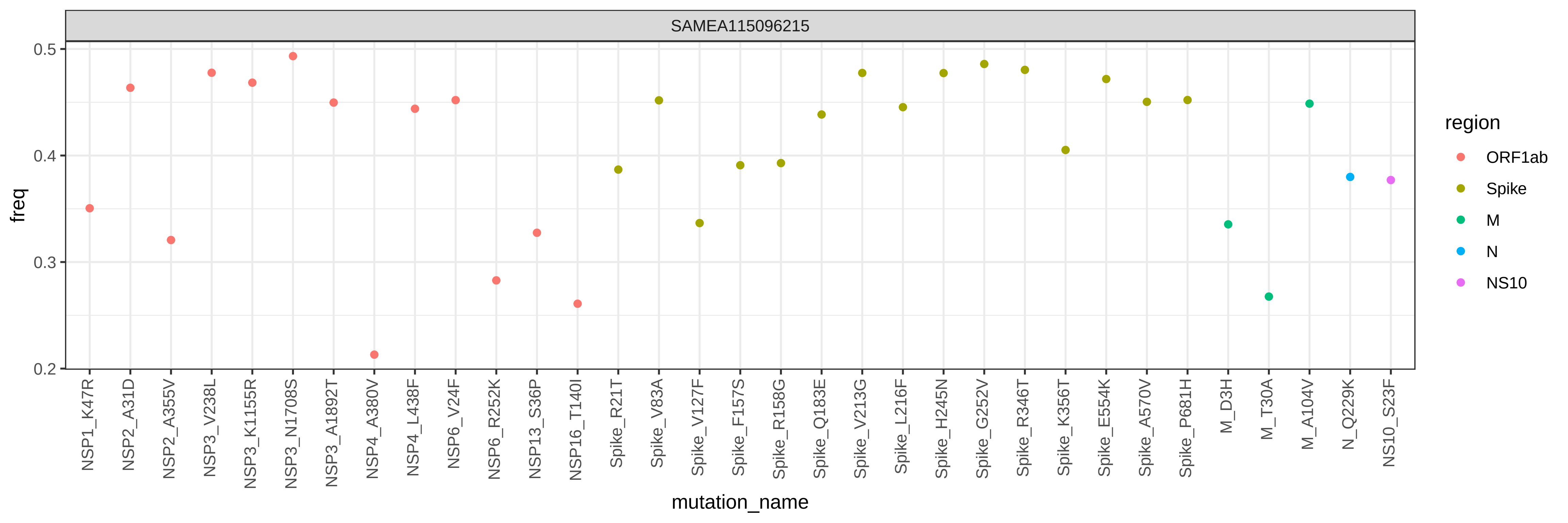

SAMEA115096215
freq
0.5
0.4
0.3
0.2
mutation_name
NSP1_K47R
NSP2_A31D
NSP2_A355V
NSP3_V238L
NSP3_K1155R
NSP3_N1708S
NSP3_A1892T
NSP4_A380V
NSP4_L438F
NSP6_V24F
NSP6_R252K
NSP13_S36P
NSP16_T140I
Spike_R21T
Spike_V83A
Spike_V127F
Spike_F157S
Spike_R158G
Spike_Q183E
Spike_V213G
Spike_L216F
Spike_H245N
Spike_G252V
Spike_R346T
Spike_K356T
Spike_E554K
Spike_A570V
Spike_P681H
M_D3H
M_T30A
M_A104V
N_Q229K
NS10_S23F
region
ORF1ab
Spike
M
N
NS10

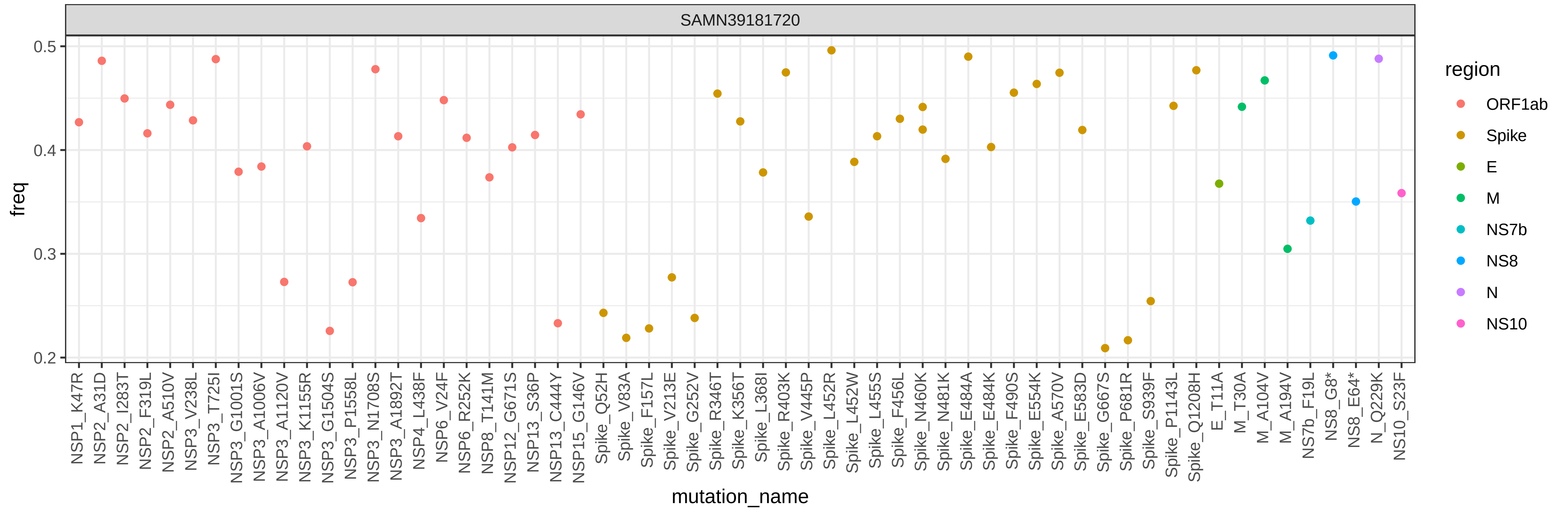
SAMN39181720
freq
0.5
0.4
0.3
0.2
mutation_name
NSP1_K47R
NSP2_A31D
NSP2_I283T
NSP2_F319L
NSP2_A510V
NSP3_V238L
NSP3_T725I
NSP3_G1001S
NSP3_A1006V
NSP3_A1120V
NSP3_K1155R
NSP3_G1504S
NSP3_P1558L
NSP3_N1708S
NSP3_A1892T
NSP4_L438F
NSP6_V24F
NSP6_R252K
NSP8_T141M
NSP12_G671S
NSP13_S36P
NSP13_C444Y
NSP15_G146V
Spike_Q52H
Spike_V83A
Spike_F157L
Spike_V213E
Spike_G252V
Spike_R346T
Spike_K356T
Spike_L368I
Spike_R403K
Spike_V445P
Spike_L452R
Spike_L452W
Spike_L455S
Spike_F456L
Spike_N460K
Spike_N481K
Spike_E484A
Spike_E484K
Spike_F490S
Spike_E554K
Spike_A570V
Spike_E583D
Spike_G667S
Spike_P681R
Spike_S939F
Spike_P1143L
Spike_Q1208H
E_T11A
M_T30A
M_A104V
M_A194V
NS7b_F19L
NS8_G8*
NS8_E64*
N_Q229K
NS10_S23F
region
ORF1ab
Spike
E
M
NS7b
NS8
N
NS10

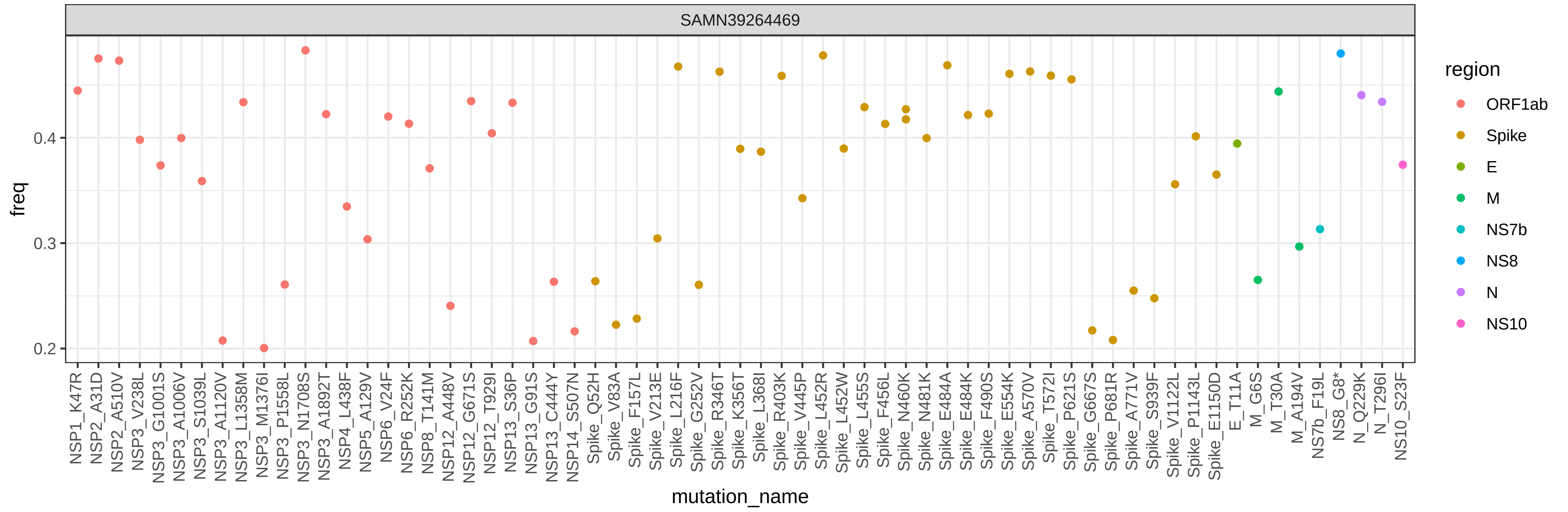
SAMN39264469
freq
0.2
0.3
0.4
mutation_name
NSP1_K47R
NSP2_A31D
NSP2_A510V
NSP3_V238L
NSP3_G1001S
NSP3_A1006V
NSP3_S1039L
NSP3_A1120V
NSP3_L1358M
NSP3_M1376I
NSP3_P1558L
NSP3_N1708S
NSP3_A1892T
NSP4_L438F
NSP5_A129V
NSP6_V24F
NSP6_R252K
NSP8_T141M
NSP12_A448V
NSP12_G671S
NSP12_T929I
NSP13_S36P
NSP13_G91S
NSP13_C444Y
NSP14_S507N
Spike_Q52H
Spike_V83A
Spike_F157L
Spike_V213E
Spike_L216F
Spike_G252V
Spike_R346T
Spike_K356T
Spike_L368I
Spike_R403K
Spike_V445P
Spike_L452R
Spike_L452W
Spike_L455S
Spike_F456L
Spike_N460K
Spike_N481K
Spike_E484A
Spike_E484K
Spike_F490S
Spike_E554K
Spike_A570V
Spike_T572I
Spike_P621S
Spike_G667S
Spike_P681R
Spike_A771V
Spike_S939F
Spike_V1122L
Spike_P1143L
Spike_E1150D
E_T11A
M_G6S
M_T30A
M_A194V
NS7b_F19L
NS8_G8*
N_Q229K
N_T296I
NS10_S23F
region
ORF1ab
Spike
E
M
NS7b
NS8
N
NS10

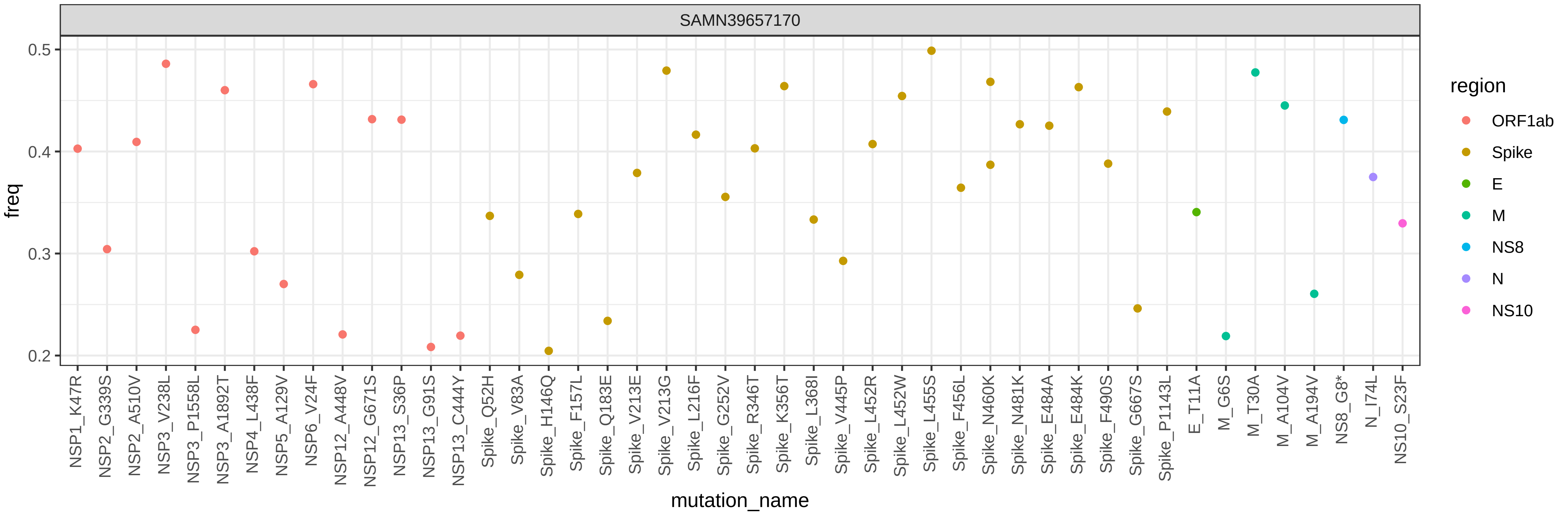
SAMN39657170
freq
0.5
0.4
0.3
0.2
mutation_name
NSP1_K47R
NSP2_G339S
NSP2_A510V
NSP3_V238L
NSP3_P1558L
NSP3_A1892T
NSP4_L438F
NSP5_A129V
NSP6_V24F
NSP12_A448V
NSP12_G671S
NSP13_S36P
NSP13_G91S
NSP13_C444Y
Spike_Q52H
Spike_V83A
Spike_H146Q
Spike_F157L
Spike_Q183E
Spike_V213E
Spike_V213G
Spike_L216F
Spike_G252V
Spike_R346T
Spike_K356T
Spike_L368I
Spike_V445P
Spike_L452R
Spike_L452W
Spike_L455S
Spike_F456L
Spike_N460K
Spike_N481K
Spike_E484A
Spike_E484K
Spike_F490S
Spike_G667S
Spike_P1143L
E_T11A
M_G6S
M_T30A
M_A104V
M_A194V
NS8_G8*
N_I74L
NS10_S23F
region
ORF1ab
Spike
E
M
NS8
N
NS10

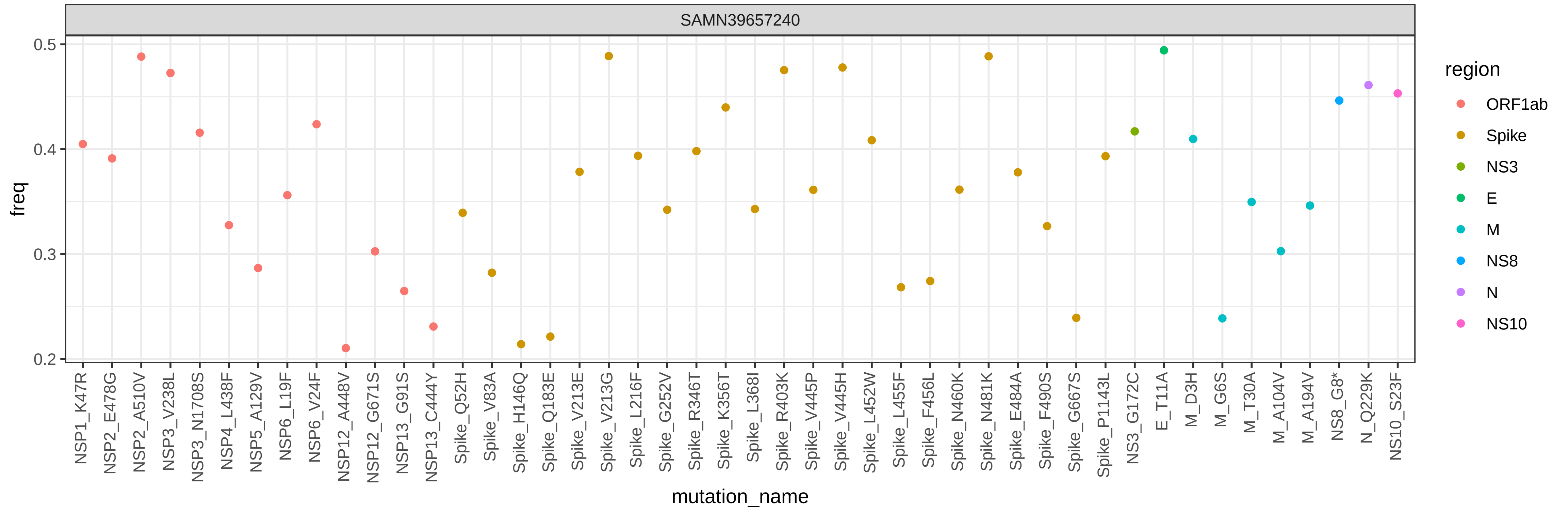
SAMN39657240
freq
0.5
0.4
0.3
0.2
mutation_name
NSP1_K47R
NSP2_E478G
NSP2_A510V
NSP3_V238L
NSP3_N1708S
NSP4_L438F
NSP5_A129V
NSP6_L19F
NSP6_V24F
NSP12_A448V
NSP12_G671S
NSP13_G91S
NSP13_C444Y
Spike_Q52H
Spike_V83A
Spike_H146Q
Spike_Q183E
Spike_V213E
Spike_V213G
Spike_L216F
Spike_G252V
Spike_R346T
Spike_K356T
Spike_L368I
Spike_R403K
Spike_V445P
Spike_V445H
Spike_L452W
Spike_L455F
Spike_F456L
Spike_N460K
Spike_N481K
Spike_E484A
Spike_F490S
Spike_G667S
Spike_P1143L
NS3_G172C
E_T11A
M_D3H
M_G6S
M_T30A
M_A104V
M_A194V
NS8_G8*
N_Q229K
NS10_S23F
region
ORF1ab
Spike
NS3
E
M
NS8
N
NS10

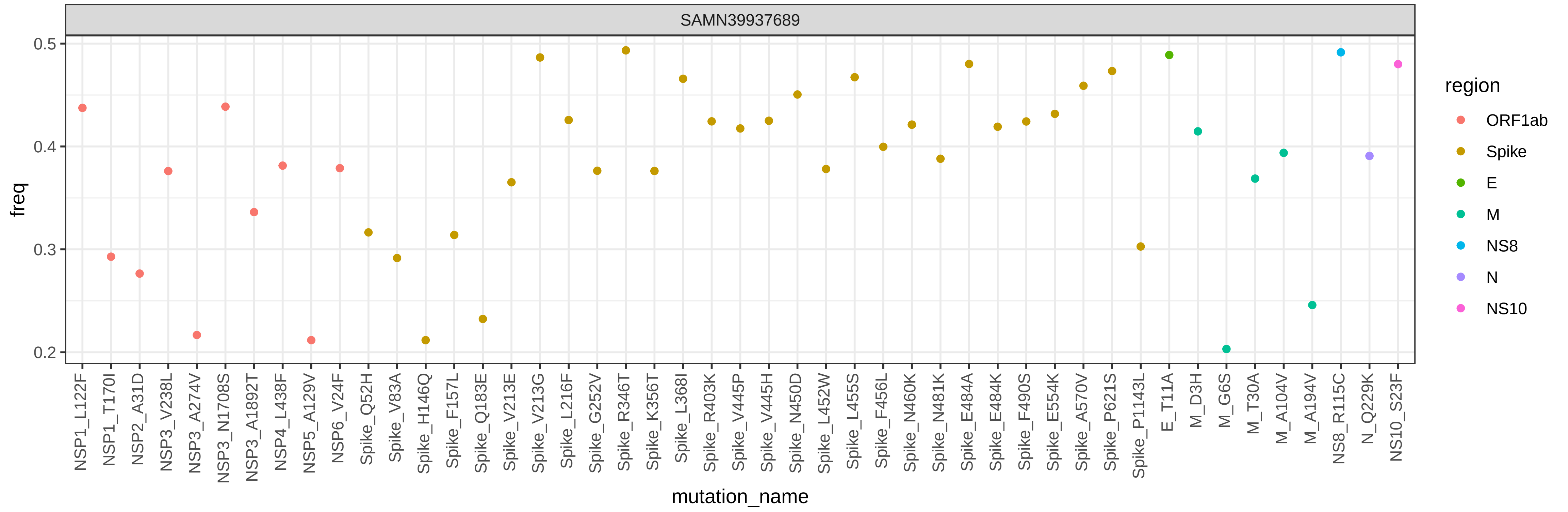
SAMN39937689
freq
0.5
0.4
0.3
0.2
mutation_name
NSP1_L122F
NSP1_T170I
NSP2_A31D
NSP3_V238L
NSP3_A274V
NSP3_N1708S
NSP3_A1892T
NSP4_L438F
NSP5_A129V
NSP6_V24F
Spike_Q52H
Spike_V83A
Spike_H146Q
Spike_F157L
Spike_Q183E
Spike_V213E
Spike_V213G
Spike_L216F
Spike_G252V
Spike_R346T
Spike_K356T
Spike_L368I
Spike_R403K
Spike_V445P
Spike_V445H
Spike_N450D
Spike_L452W
Spike_L455S
Spike_F456L
Spike_N460K
Spike_N481K
Spike_E484A
Spike_E484K
Spike_F490S
Spike_E554K
Spike_A570V
Spike_P621S
Spike_P1143L
E_T11A
M_D3H
M_G6S
M_T30A
M_A104V
M_A194V
NS8_R115C
N_Q229K
NS10_S23F
region
ORF1ab
Spike
E
M
NS8
N
NS10

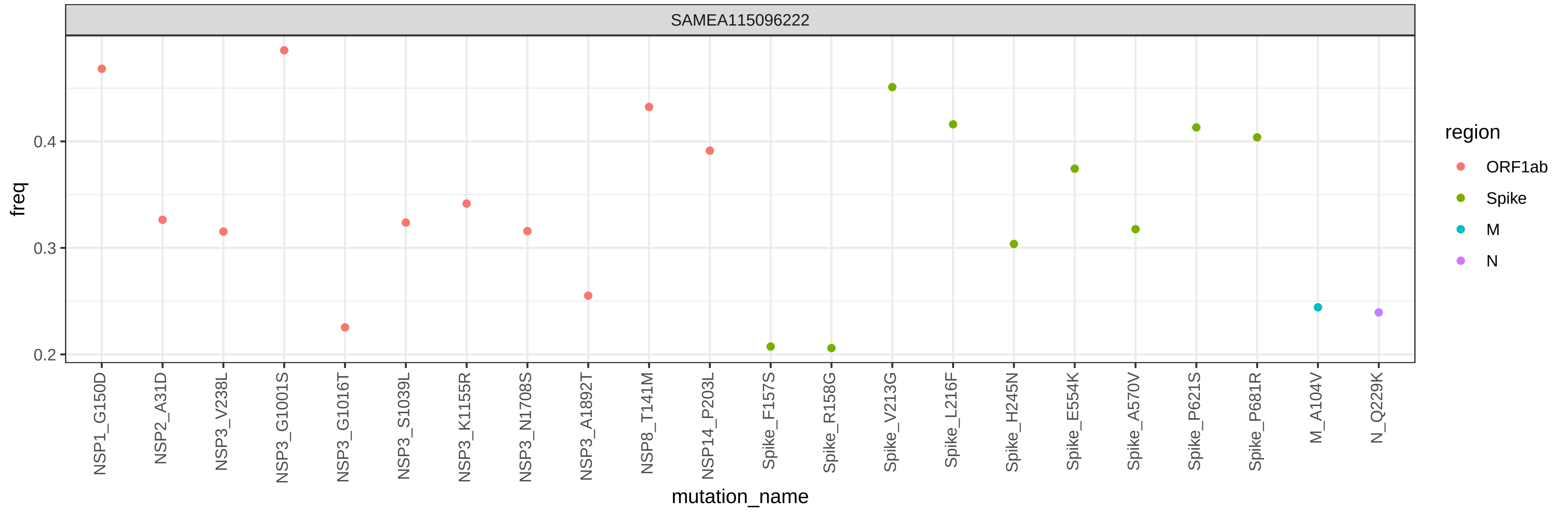
SAMEA115096222
freq
0.2
0.3
0.4
mutation_name
NSP1_G150D
NSP2_A31D
NSP3_V238L
NSP3_G1001S
NSP3_G1016T
NSP3_S1039L
NSP3_K1155R
NSP3_N1708S
NSP3_A1892T
NSP8_T141M
NSP14_P203L
Spike_F157S
Spike_R158G
Spike_V213G
Spike_L216F
Spike_H245N
Spike_E554K
Spike_A570V
Spike_P621S
Spike_P681R
M_A104V
N_Q229K
region
ORF1ab
Spike
M
N

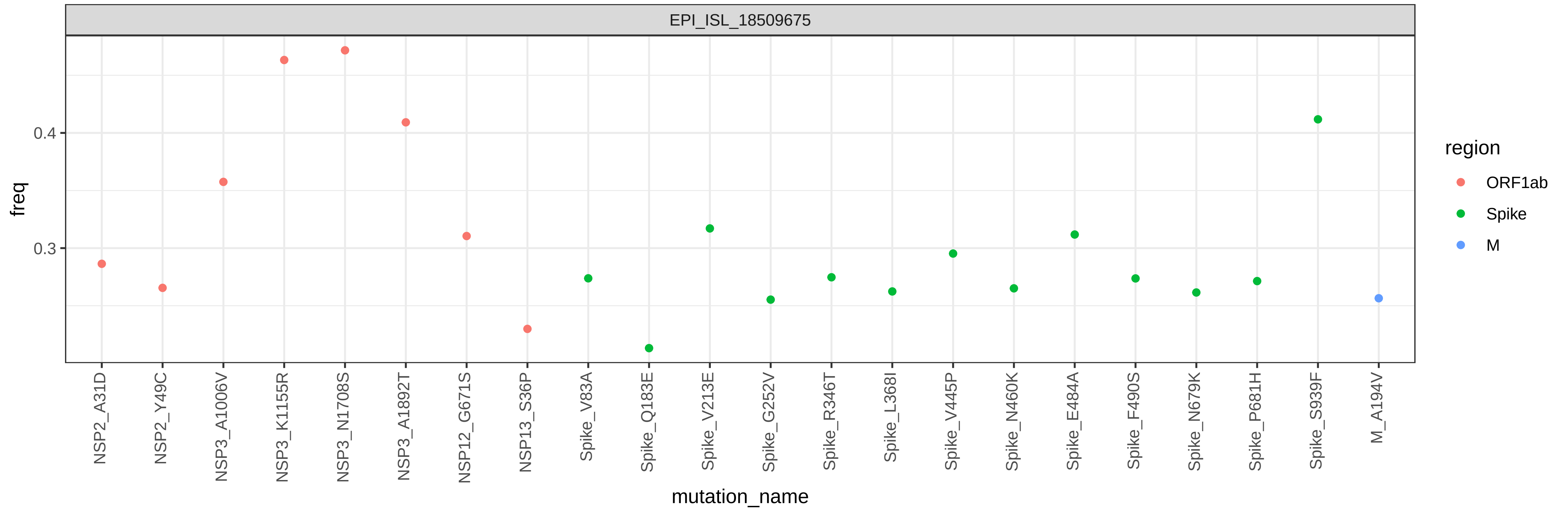

EPI_ISL_18509675
freq
0.4
0.3
mutation_name
NSP2_A31D
NSP2_Y49C
NSP3_A1006V
NSP3_K1155R
NSP3_N1708S
NSP3_A1892T
NSP12_G671S
NSP13_S36P
Spike_V83A
Spike_Q183E
Spike_V213E
Spike_G252V
Spike_R346T
Spike_L368I
Spike_V445P
Spike_N460K
Spike_E484A
Spike_F490S
Spike_N679K
Spike_P681H
Spike_S939F
M_A194V
region
ORF1ab
Spike
M

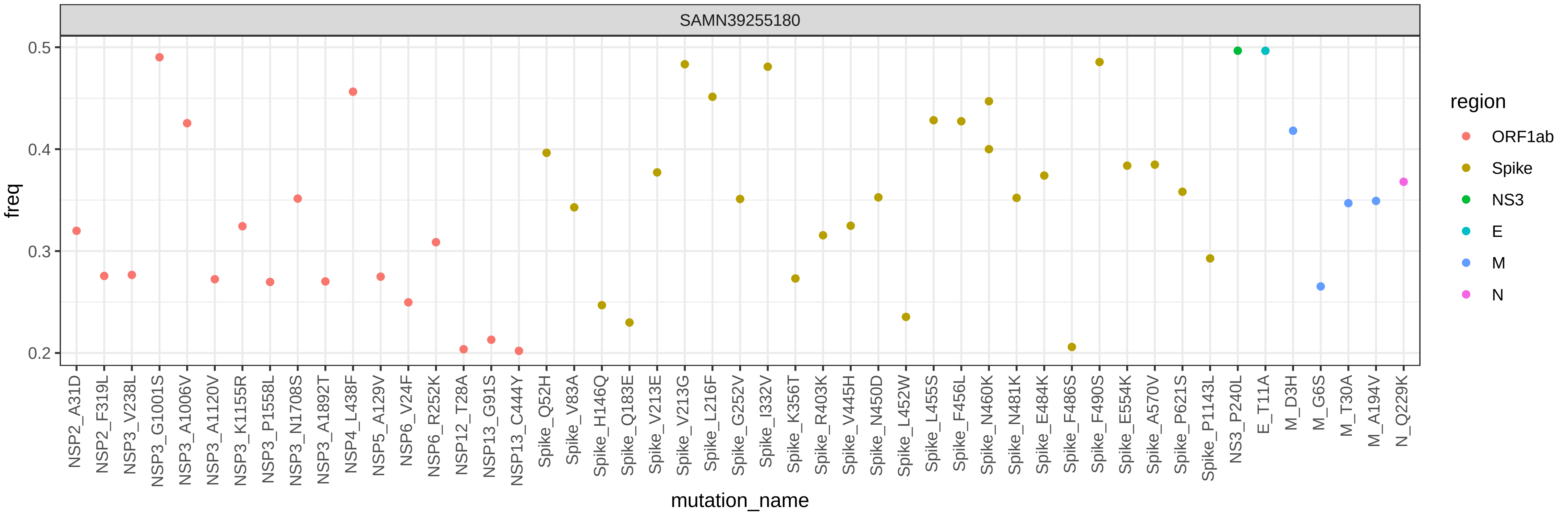

SAMN39255180
freq
0.2
0.3
0.4
0.5
mutation_name
NSP2_A31D
NSP2_F319L
NSP3_V238L
NSP3_G1001S
NSP3_A1006V
NSP3_A1120V
NSP3_K1155R
NSP3_P1558L
NSP3_N1708S
NSP3_A1892T
NSP4_L438F
NSP5_A129V
NSP6_V24F
NSP6_R252K
NSP12_T28A
NSP13_G91S
NSP13_C444Y
Spike_Q52H
Spike_V83A
Spike_H146Q
Spike_Q183E
Spike_V213E
Spike_V213G
Spike_L216F
Spike_G252V
Spike_I332V
Spike_K356T
Spike_R403K
Spike_V445H
Spike_N450D
Spike_L452W
Spike_L455S
Spike_F456L
Spike_N460K
Spike_N481K
Spike_E484K
Spike_F486S
Spike_F490S
Spike_E554K
Spike_A570V
Spike_P621S
Spike_P1143L
NS3_P240L
E_T11A
M_D3H
M_G6S
M_T30A
M_A194V
N_Q229K
region
ORF1ab
Spike
NS3
E
M
N

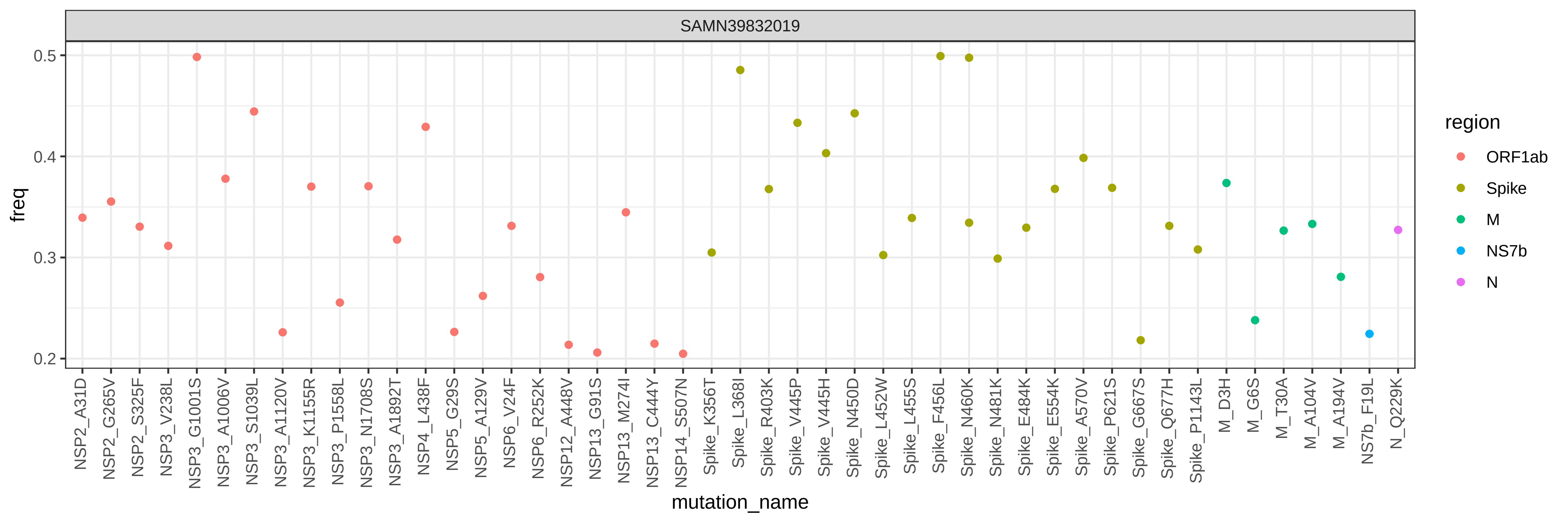

SAMN39832019
freq
0.5
0.4
0.3
0.2
mutation_name
NSP2_A31D
NSP2_G265V
NSP2_S325F
NSP3_V238L
NSP3_G1001S
NSP3_A1006V
NSP3_S1039L
NSP3_A1120V
NSP3_K1155R
NSP3_P1558L
NSP3_N1708S
NSP3_A1892T
NSP4_L438F
NSP5_G29S
NSP5_A129V
NSP6_V24F
NSP6_R252K
NSP12_A448V
NSP13_G91S
NSP13_M274I
NSP13_C444Y
NSP14_S507N
Spike_K356T
Spike_L368I
Spike_R403K
Spike_V445P
Spike_V445H
Spike_N450D
Spike_L452W
Spike_L455S
Spike_F456L
Spike_N460K
Spike_N481K
Spike_E484K
Spike_E554K
Spike_A570V
Spike_P621S
Spike_G667S
Spike_Q677H
Spike_P1143L
M_D3H
M_G6S
M_T30A
M_A104V
M_A194V
NS7b_F19L
N_Q229K
region
ORF1ab
Spike
M
NS7b
N

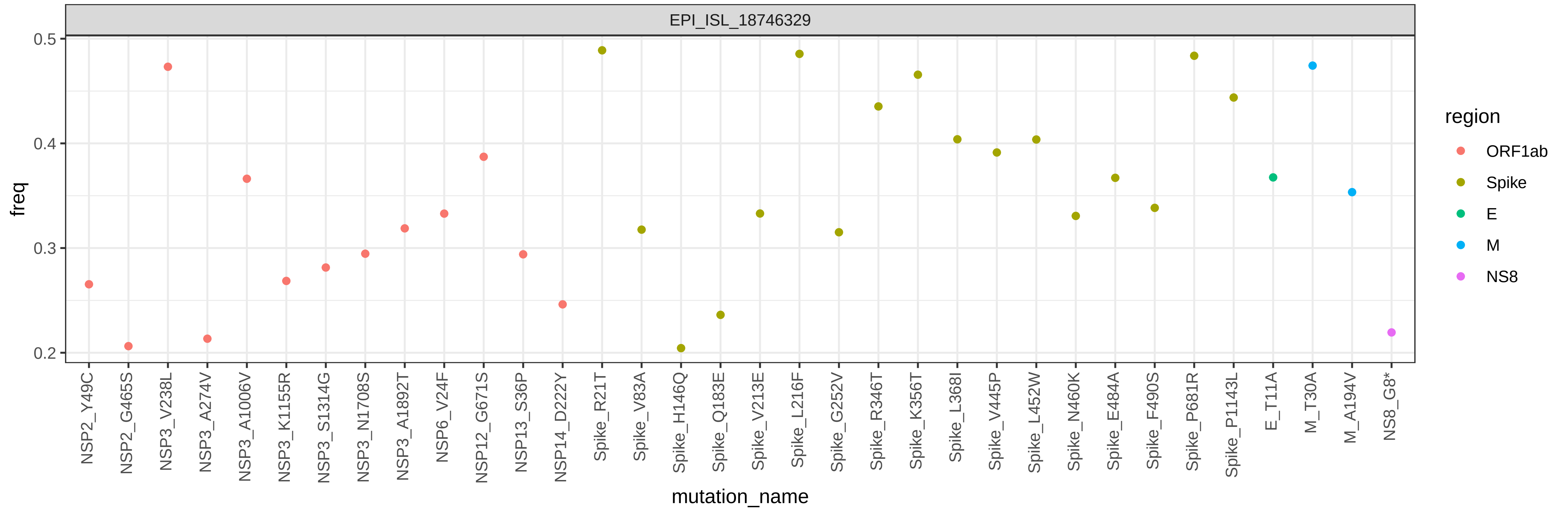

EPI_ISL_18746329
freq
0.2
0.3
0.4
0.5
region
ORF1ab
Spike
E
M
NS8
mutation_name
NSP2_Y49C
NSP2_G465S
NSP3_V238L
NSP3_A274V
NSP3_A1006V
NSP3_K1155R
NSP3_S1314G
NSP3_N1708S
NSP3_A1892T
NSP6_V24F
NSP12_G671S
NSP13_S36P
NSP14_D222Y
Spike_R21T
Spike_V83A
Spike_H146Q
Spike_Q183E
Spike_V213E
Spike_L216F
Spike_G252V
Spike_R346T
Spike_K356T
Spike_L368I
Spike_V445P
Spike_L452W
Spike_N460K
Spike_E484A
Spike_F490S
Spike_P681R
Spike_P1143L
E_T11A
M_T30A
M_A194V
NS8_G8*

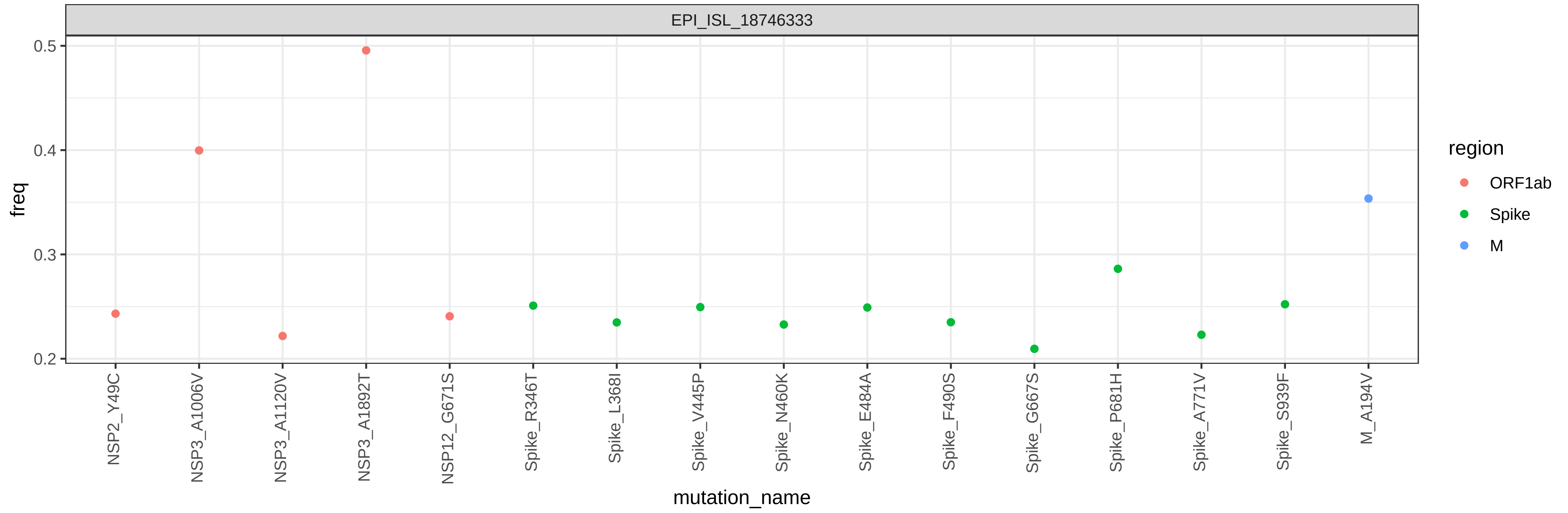
EPI_ISL_18746333
freq
0.2
0.3
0.4
0.5
NSP2_Y49C
NSP3_A1006V
NSP3_A1120V
NSP3_A1892T
NSP12_G671S
Spike_R346T
Spike_L368I
Spike_V445P
Spike_N460K
Spike_E484A
Spike_F490S
Spike_G667S
Spike_P681H
Spike_A771V
Spike_S939F
M_A194V
mutation_name
region
ORF1ab
Spike
M

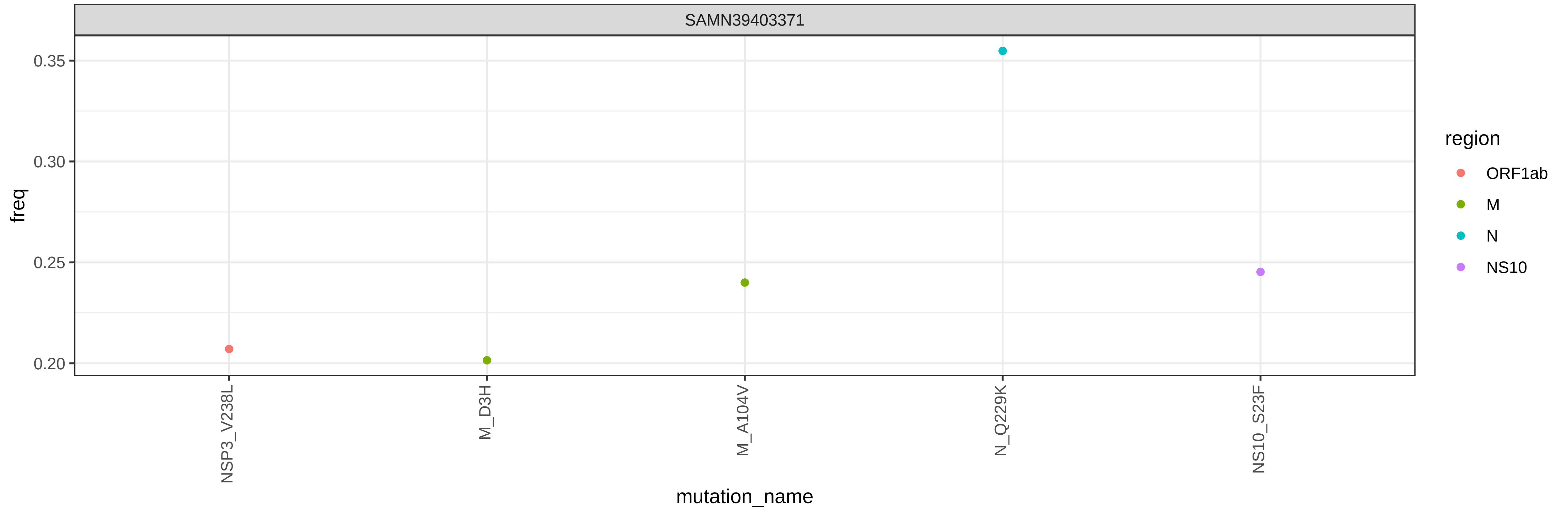
SAMN39403371
freq
0.35
0.30
0.25
0.20
NSP3_V238L
M_D3H
M_A104V
N_Q229K
NS10_S23F
mutation_name
region
ORF1ab
M
N
NS10

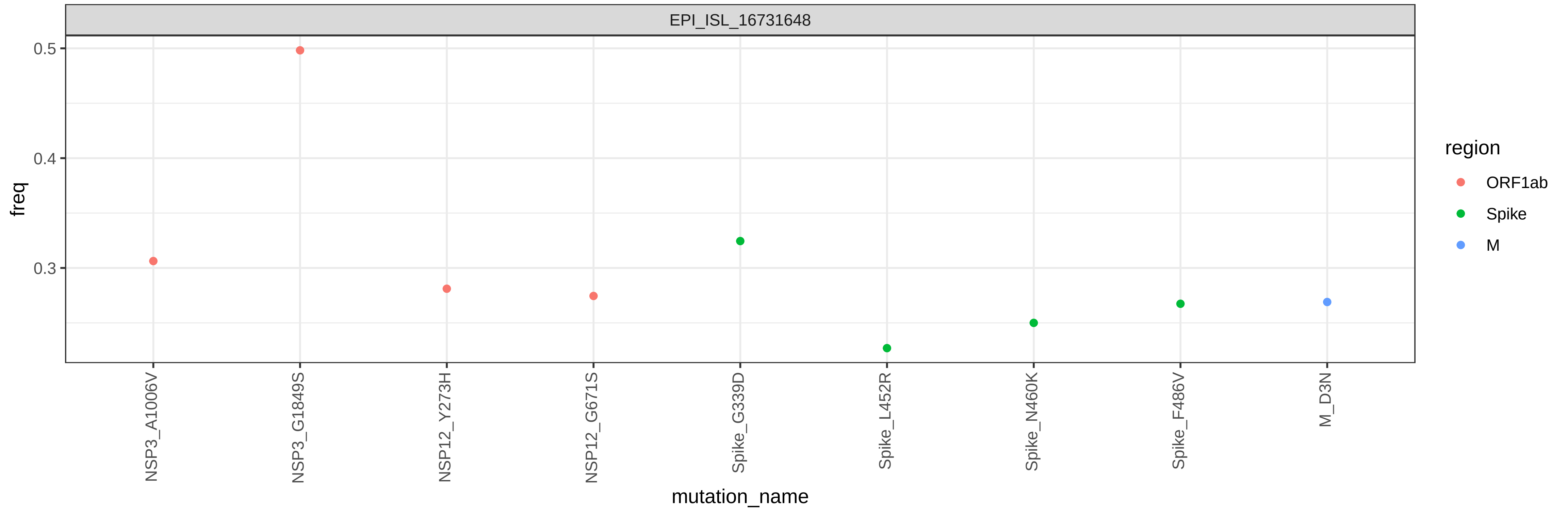
EPI_ISL_16731648
freq
0.5
0.4
0.3
NSP3_A1006V
NSP3_G1849S
NSP12_Y273H
NSP12_G671S
Spike_G339D
Spike_L452R
Spike_N460K
Spike_F486V
M_D3N
mutation_name
region
ORF1ab
Spike
M

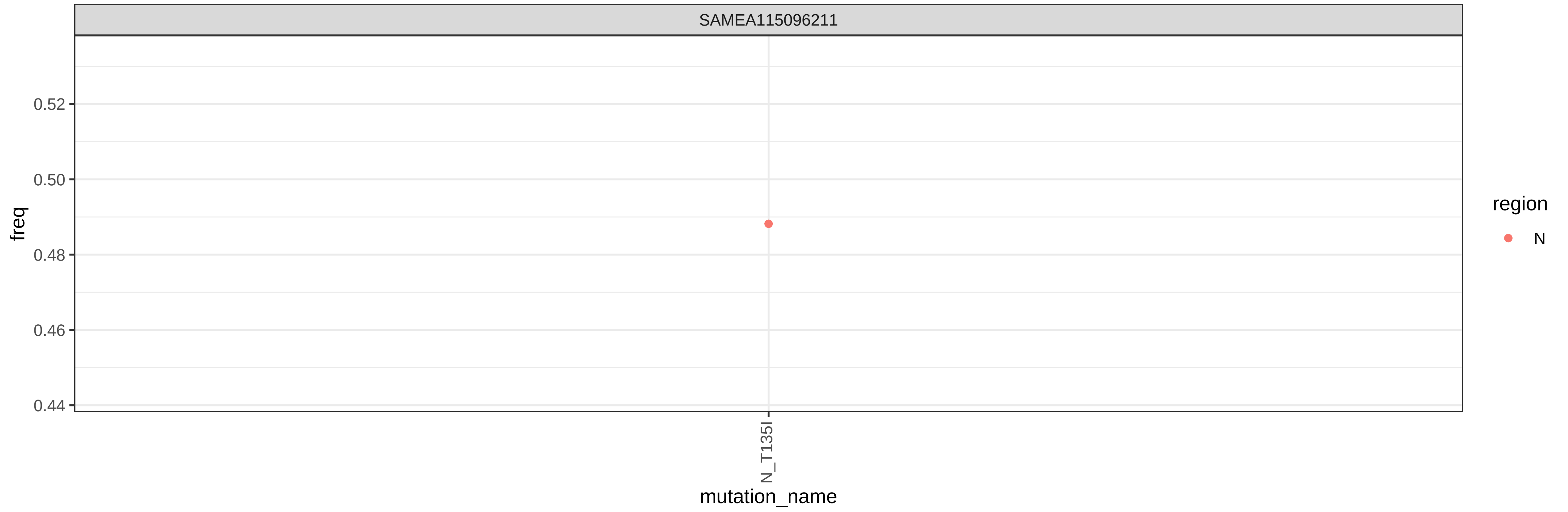

SAMEA115096211
freq
0.52
0.50
0.48
0.46
0.44
N_T135I
mutation_name
region
N

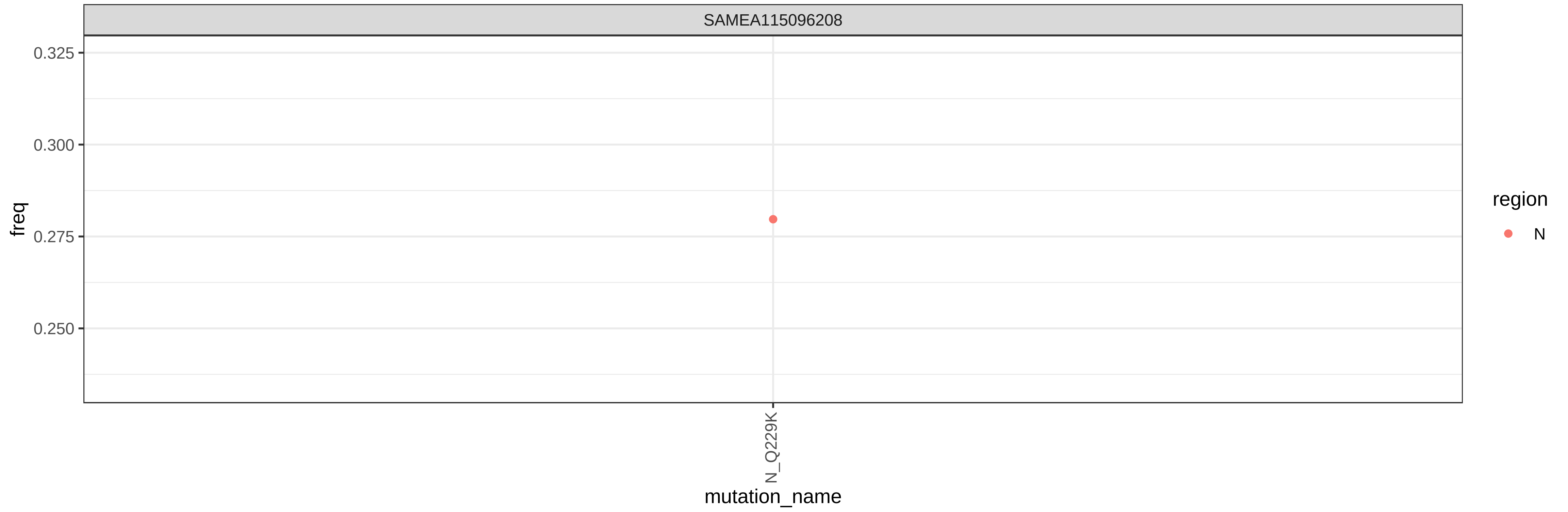
SAMEA115096208
0.325
0.300
0.275
0.250
freq
N_Q229K
mutation_name
region
N
